# Structural analogue-based protein structure domain assembly assisted by deep learning

**DOI:** 10.1101/2022.03.07.483151

**Authors:** Chun-Xiang Peng, Xiao-Gen Zhou, Yu-Hao Xia, Jun Liu, Ming-Hua Hou, Gui-Jun Zhang

## Abstract

**Motivation:** With the breakthrough of AlphaFold2, the protein structure prediction problem has made a remarkable progress through end-to-end deep learning techniques, in which correct folds could be built for nearly all single-domain proteins. However, the full-chain modelling appears to be lower on average accuracy than that for the constituent domains and requires higher demand on computing hardware, indicating the performance of full-chain modelling still needs to be improved. In this study, we investigate whether the predicted accuracy of full-chain model can be further improved by domain assembly assisted by deep learning.

**Results:** In this article, we developed a structural analogue-based protein structure domain assembly method assisted by deep learning, named SADA. In SADA, a multi-domain protein structure database (MPDB) was constructed for the full-chain analogue detection using individual domain models. Starting from the initial model constructed from the analogue, the domain assembly simulation was performed to generate the full-chain model through a two-stage differential evolution algorithm guided by the physics-based force field and an inter-residue distance potential predicted by deep learning. SADA was compared with the state-of-the-art domain assembly methods on 356 benchmark proteins, and the average TM-score of SADA models is 8.1% and 27.0% higher than that of DEMO and AIDA, respectively. We also assembled full-chain models of 20 human multi-domain proteins using individual domain models independently predicted by AlphaFold2, where the SADA full-chain models obtained a 4.8% higher average TM-score than full-chain models directly predicted by AlphaFold2 and fewer computing resources were required. In addition, we also find that the domains often interact in the similar way in the quaternary orientations if the domains have similar tertiary structures. Furthermore, homologous templates and structural analogues are complementary for multi-domain protein full-chain modelling.

**Availability:** The SADA web server are freely available at http://zhanglab-bioinf.com/SADA.

## 1 Introduction

Multi-domain proteins are an important class of proteins and consist of more than one unique folding units that are connected into a single chain. Many biological functions rely on the interaction of different domains (Zhou *et al*., 2022). For example, in ribose-binding protein in bacterial transport and chemotaxis, the ligand-binding site is located in a pocket formed by two domains. More than 80% of eukaryotic proteins and 67% of prokaryotic proteins contain multiple domains. However, only one third of structures in the Protein Data Bank (PDB) contain multiple domains, which indicates that it may be more difficult to experimentally determine the structure of multi-domain protein structure.

With the development of deep learning, protein structure prediction methods have been developed successively, such as trRosetta (Yang *et al*., 2020; Su *et al*., 2021; Du *et al*., 2021), RaptorX (Xu, 2019; Xu and Wang, 2019), RocketX (Liu *et al*., 2022), and D-I-TASSER (Zheng *et al*., 2021). Recently, end-to-end methods, such as AlphaFold2 (Jumper *et al*., 2021) and RoseTTAFold (Baek *et al*., 2021), have been proposed to accurately predict more complex proteins including some multi-domain proteins. The performance of these methods relies to some extent on the quality of the MSA or the homologous template (Pearce and Zhang, 2021). For example, the distributions of average confidence scores for AlphaFold2 models of human proteins with and without homologues available in the PDB, were different (Jones and Thornton, 2022). However, homologues available in the PDB may be fewer for multi-domain proteins, which may further affect the prediction performance of multi-domain proteins. Meanwhile, the end-to-end approach entails high demand on computing hardware. For relatively large proteins (i.e., >800 amino acids), most computer memory hardly satisfies its training and running requirements, which may not be helpful to the study of their intrinsic mechanisms. Therefore, based on the divide-and-conquer strategy, modelling the full-chain structure of multi-domain protein by domain assembly may be a lightweight alternative way. The way may be helpful for further improving the accuracy of end-to-end methods and revealing the folding mechanism, but they have been largely ignored by the community (Zhou *et al*., 2019).

In general, domain assembly method is mainly divided into two categories: *de novo*-based methods and template-based approaches. *De novo*-based methods focus mainly on construction of the linker models by some *de novo* or *ab initio* folding potentials. In Rosetta (Wollacott *et al*., 2007), the method of assembling structures of multi-domain proteins consists of an initial low-resolution search, in which the conformational space of the domain linker is explored using the Rosetta de novo structure prediction method, followed by a high-resolution search, in which all atoms are treated explicitly, and the backbone and side chain degrees of freedom are simultaneously optimised. In AIDA (Xu *et al*., 2015), domain assembly for assembling multi-domain protein structures is simulated by a fast-docking algorithm with an *ab initio* folding potential. The *de novo*- or *ab initio*-based methods may leave the domain structures largely randomly oriented in the final model (Zhou *et al*., 2019). Template based approaches often detect available templates to guide domain assembly. DEMO (Zhou *et al*., 2019) is used for constructing multi-domain protein structures by docking-based domain assembly simulations, in which inter-domain orientations are determined by the distance profiles from analogous templates as detected through domain-level structure alignments. However, the template-based approaches also have limitations because of the limited number of multi-domain proteins in PDB, and the difficulty of capturing the orientation between domains from the template maybe increase as the number of domains increases. Using deep learning technology may be helpful to capture the orientation information between domains that cannot be obtained from templates.

In this work, we proposed a new domain assembly method, SADA, in which the domain assembly simulation was performed to generate the full-chain model through a two-stage differential evolution algorithm guided by the physics-based force field and an inter-residue distance potential predicted by deep learning. SADA was test on a benchmark set of 356 proteins, for which the performance of SADA significantly outperformed most state-of-the-art domain assembly methods, starting from randomly oriented target native domain structures. Especially, on 40 benchmark proteins with the number of domains ≥ 4, the average TM-score of SADA is 0.60, which is 13.2% higher than that of the second-best method. Further, SADA was used to reassemble the full-chain model of AlphaFold2 on 20 human multi-domain proteins, where the average TM-score of the models was improved from 0.63 to 0.65, starting from domain models decomposed from full-chain model of AlphaFold2. In the 20 proteins, the average TM-score of SADA full-chain models achieved 0.66, starting from individual domain models independently predicted by AlphaFold2. Compared to the full-chain model of AlphaFold2, the quality of the full-chain models of half of the 20 proteins was improved after being assembled by SADA, and some cases achieved significantly better models. In addition, we also provided two additional function modules in webserver, including a culling module to filter the whole MPDB according to input criteria and a detection module to identify the structural analogues of full-chain according to input domain models.

## 2 Methods

SADA is a structural analogue-based protein structure domain assembly method assisted by deep learning, which involves five steps as follows: (1) Detects structural analogues of the full-chain from the constructed MPDB according to the input protein domain models; (2) Constructs an initial model based on the detected 1st-ranked analogue; (3) Utilizes a deep learning network to predict the inter-residue distance distribution; (4) Builds multi-domain protein specificity force field for guiding domain assembly based on the predicted residue distance distribution and the property of multi-domain protein; and (5) Assembles the domain models to generate final full-chain model by the proposed two-stage differential evolution algorithm from the initial model. The pipeline is displayed in Figure 1a.

**Fig. 1.**
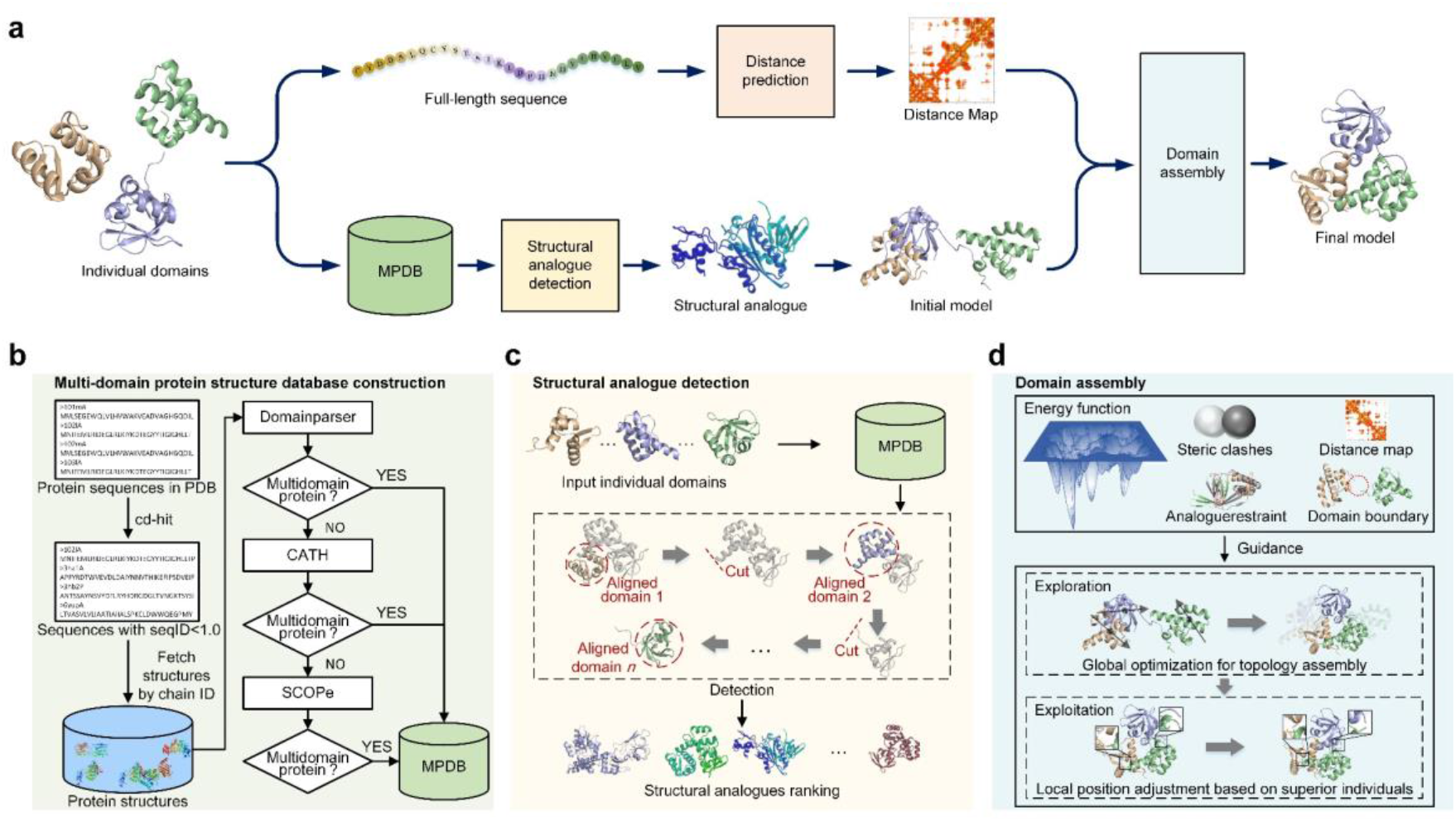
(a) Pipeline of SADA for domain assembly. (b) Flowchart of multi-domain protein structure database construction. (c) Illustration of structural analogues detection. (d) Illustration of two-stage differential evolution algorithm for domain assembly.

### 2.1 MPDB construction

The flowchart of MPDB construction is shown in Figure 1b. The collection and processing steps of multi-domain protein data in MPDB are as follows: (1) CD-HIT (Fu *et al*., 2012) was used to remove the redundancy of protein structures with a sequence identity cut-off of 100% in PDB, and then protein structures with sequence identity of less than 100% were obtained from PDB; (2) DomainParser (Xu *et al*., 2000) was next used to determine whether these proteins are multi-domain proteins or not; and (3) the single-domain proteins determined by DomainParser were further confirmed by the definition of CATH (Orengo *et al*., 1997; Lam *et al*., 2016) and SCOPe (Chandonia *et al*., 2017) on whether they were multi-domain proteins. All the multi-domain proteins selected in the three steps above were finally collected to construct the MPDB. The used parameters of CD-HIT are listed in Supplementary Test S1.

As of September 2021, MPDB contains 48,225 multi-domain proteins, in which 37,495 proteins have 2 domains, 7,539 proteins have 3 domains, 2,182 proteins have 4 domains, and 1,009 proteins have more than 4 domains. The statistics of the number of domains in the MPDB are shown in Figure S1.

For many purposes, it is useful to obtain a subset from MPDB. It is often the case that additional criteria are desirable, such as resolution, length, sequence identity, or number of domains cut-offs. For example, in protein structure prediction methods based on machine learning, the proteins that satisfy specific criteria are used to train or test. Inspired by PISCES (Wang and Dunbrack, 2003), we developed a module for retrieving multi-domain proteins in MPDB, which filters the whole MPDB according to user’s input criteria including protein chain length, resolution, number of domains, Rfactor, and sequence identity of multi-domain protein, and then provides the protein structures and related information that satisfy the criteria to users. The module is described in Supplementary Text S2.

### 2.2 Structural analogue detection

Homologues inherit similarities from their common ancestor, while analogues converge to similar structures due to a limited number of energetically favourable ways to pack secondary structural elements. In some cases, inferring structural analogues of the full-chain based on the domain structures may be more critical for research and modelling of multi-domain proteins.

Based on MPDB, we designed an algorithm to detect the multi-domain protein structural analogues. The structural analogue detection is illustrated in Figure 1c. To prevent structurally similar domains from being matched to the same part of the protein, the input individual domains were aligned on each protein in MPDB by using TM-align (Zhang and Skolnick, 2005, 2004), with no overlap allowed in the alignments of different domains. According to the structural similarity of individual domain models, a local similarity score *LS*_score_ was designed to evaluate the quality of structural analogues. *LS*_score_ is defined as follows:

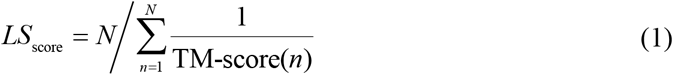

where TM-score(*n*) is the TM-score between the *n*-th domain of the target and the template protein in MPDB after aligning the domain on the template protein by TM-align. *N* is the number of query individual domains. The *LS*_score_ has a value range of (0, 1], where 1 indicates a perfect match between domain models and the structural analogues of the full-chain.

In addition, although structural analogues serve as important parts of protein structural modelling and research, limited studies have attempted to recognise structural analogues (Javier *et al*., 2020). Thus, we provide the function module to detect structural analogues from MPDB, and the module is described in Supplementary Text S3.

### 2.3 Force field for domain assembly

The force field was designed based on Anfinsen’s hypothesis, in which native-like protein conformations represent unique, low-energy, thermodynamically stable conformations. Domain assembly was guided by the force field, which was computed as a linear combination of four energy terms below,

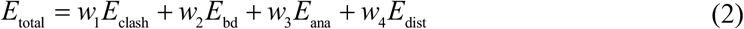

in which each energy function term is explained.

*E*_clash_ represents the C_α_ clashes between domains, which is used to prevent the domain model from getting too close during assembly to satisfy the physical concept. This term is computed using Equation (3):

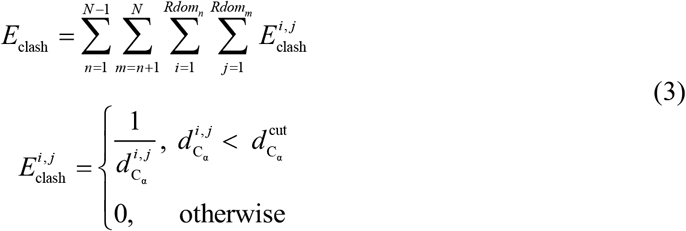

where 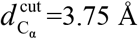, *N* is the number of domain models, *Rdom_n_* and *Rdom_m_* represent the number of residues in the *n*-th and *m*-th domain, respectively, and 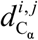 is the distance between C_α_ of the residue *i* and *j* in the evaluated decoy.

*E*_bd_ is the boundary distance energy, which is defined as follows:

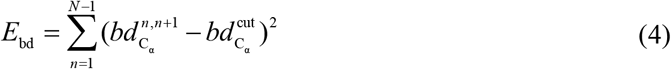

where the 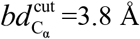, and 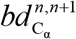 is the C_α_ atom distance between the C-terminal residue of *n*-th domain and *N*-terminal residue of the (*n* + 1)-th domain in the evaluated decoy. In the discontinuous domain proteins, the discontinuous domain is split into multiple segments because of the insertion of the continuous domains. Therefore, for discontinuous domain proteins, these discontinuous segments were treated as domains to calculate the energy.

*E*_ana_ is the structural analogue restrain energy. This term aims to prevent the assembly from deviating too much from the orientation obtained from initial model. *E*_ana_ is calculated as follows:

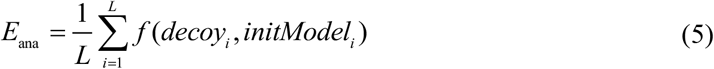

where *f*(*decoy_i_, initModel_i_*) represents the distance between the *i*-th C_α_ atom of the evaluated decoy and the *i*-th C_α_ atom of initial model, and *L* is the length of the full-chain model.

*E*_dist_ is the inter-domain distance potential. The predicted inter-residue distance provides abundant spatial constraint information for domain assembly, which can compensate for the effects of low-quality structural analogues on domain assembly.

In this study, we used GeomNet, a geometric constraints prediction network in our recently developed structure prediction server RocketX (Liu *et al*., 2022), to predict the inter-residue distance of full-chain. For the query sequence, MSA was generated by iterative search against UniRef30 (Mirdita *et al*., 2017) and BFD (Steinegger *et al*., 2019) databases by using HHblits (Steinegger *et al*., 2019) with gradually relaxed e-values of 1e^-30^, 1e^-10^, 1e^-6^, and 1e^-3^. The input features of GeomNet were extracted from MSA, including the inverse of the covariance matrix, the position specific scoring matrix, the residue position entropy, and the one-hot encoding of the query sequence. GeomNet contains 66 residual blocks, each of which consists of two 2D convolutional layers, two instance normalization layers, two ELU activation layers and a dropout layer. The predicted distances were divided into 36 equidistant bins from 2Å to 20Å.

The potential is defined as follows:

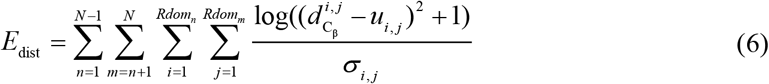

where 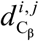 is the real distance between C_β_ atom (C_α_ for glycine) of the residue pair (*i, j*) in the evaluated decoy, and *u_ij_* and *σ_i,j_* are the mean and standard deviation obtained by Gaussian fitting of the residue pair (*i, j*) distance distribution, respectively.

The weighting parameters in Equation (2) were set as *w*_1_ = 0.52, *w*_2_ = 0.20, *w*_3_ = 0.50 and *w*_4_ = (1- *LS*_score_). If the local similarity score *LS*_score_ is greater than or equal to 0.82, *w*_3_ is set to 6.0. A high *LS*_score_ indicates that the interaction direction inferred from structural analogue is reliable. Therefore, *w*_3_ was increased, whereas *w*_4_ was decreased.

### 2.4 Two-stage differential evolution algorithm for domain assembly

The assembly engine for domain assembly was carried out through simultaneous rotation and translation of each domain. For each domain, its movement can be represented by a translation vector and three rotation angles. Therefore, for a multi-domain protein with *N* domains, the solution of domain assembly can be represented as a (6 * *N*)-dimensional target vector, and the solution *S* can be represented as follows:

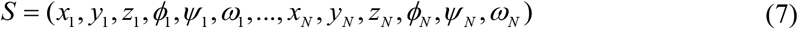

where *x_N_, y_N_* and *z_N_* represent the translation vector of the *N*-domain, and *ϕ_N_, ψ_N_* and *ω_N_* represent the three rotation angles of the *N*-domain.

Under the guidance of the force field, a two-stage differential evolution algorithm was proposed to determine the optimal solution. The exploration stage aims to prevent the algorithm from getting stuck in local optimal and generate multiple superior topology structures for the exploitation stage. Thus, the mutation strategy of slow convergence speed and strong exploration capability was used in this stage.

In the exploitation stage, the explored superior solutions rapidly converged to the minimum. Therefore, on the basis of the superior solutions generated in the previous stage, we further adjusted the local position of these solutions to generate the optimal solution, and the mutation strategy with fast convergence speed was used at this stage. The algorithm description and parameters setting of the two-stage differential evolution algorithm is described in Supplementary Text S4.

## 3 Results and discussion

### 3.1 Dataset

To fairly compare the performance of SADA with other methods, we used all the 356 proteins from DEMO benchmark dataset as the test targets (Zhou *et al*., 2019). The 356 proteins were generated by separately clustering the proteins with different domain types and structures from the template library of DEMO with a 30% sequence identity cut-off. This benchmark included 166 2-domain (2dom), 69 3-domain (3dom), 40 ≥ 4-domain (m4dom), and 81 discontinuous-domain (2dis) proteins. The maximum number of domains in m4dom is 7. Table S1 summarizes the details of the 356 test proteins, including PDB ID, protein length and the number of domains.

### 3.2 Results of benchmark set

In this test, we reassembled the individual domain structures excised from the experimental structure. The initial domain structure was randomly rotated and translated before assembly, and structural analogues with a sequence identity > 30% to the query were excluded. SADA was compared with two well-known domain assembly methods, namely, DEMO (Zhou *et al*., 2019) and AIDA (Xu *et al*., 2015). The average results of the final models of SADA, DEMO and AIDA on the benchmark set are shown in Table 1. The detailed results of each protein are shown in Supplementary Table S2 and S3.

**Table 1.** Summary of domain structure assembly by using experimentally solved domains on 356 test proteins. TM-score represents the average TM-score of final full-chain models. #TM-score ≥ 0.5 represents the number of models with TM-score ≥ 0.5. The values in the last column are the results of the Wilcoxon signed-rank test based on the comparison with the TM-score of SADA.

| Method | TM-score | #TM-score | *P*-value |
| --- | --- | --- | --- |
| SADA | 0.80 | 328 | - |
| DEMO | 0.74 | 318 | 1.28E-21 |
| AIDA | 0.63 | 288 | 8.91E-51 |

Overall, Overall, the average TM-score of SADA models is 0.80, which is 8.1% higher than that (0.74) of DEMO (with *P*-value = 1.28E-21) and 27.0% higher than that (0.63) of AIDA (with *P*-value = 8.91E-51) respectively. SADA correctly assembles (i.e., TM-score ≥ 0.5) 328 out of 356 targets, accounting for 92.1% of the total, which is 3.1% and 13.9% better than that of DEMO and AIDA, respectively. The respective *P*-values of DEMO and AIDA indicate statistically significant differences between the methods. For an intuitive comparison of the TM-score between different methods, their results are depicted in Figure 2.

**Fig. 2.**
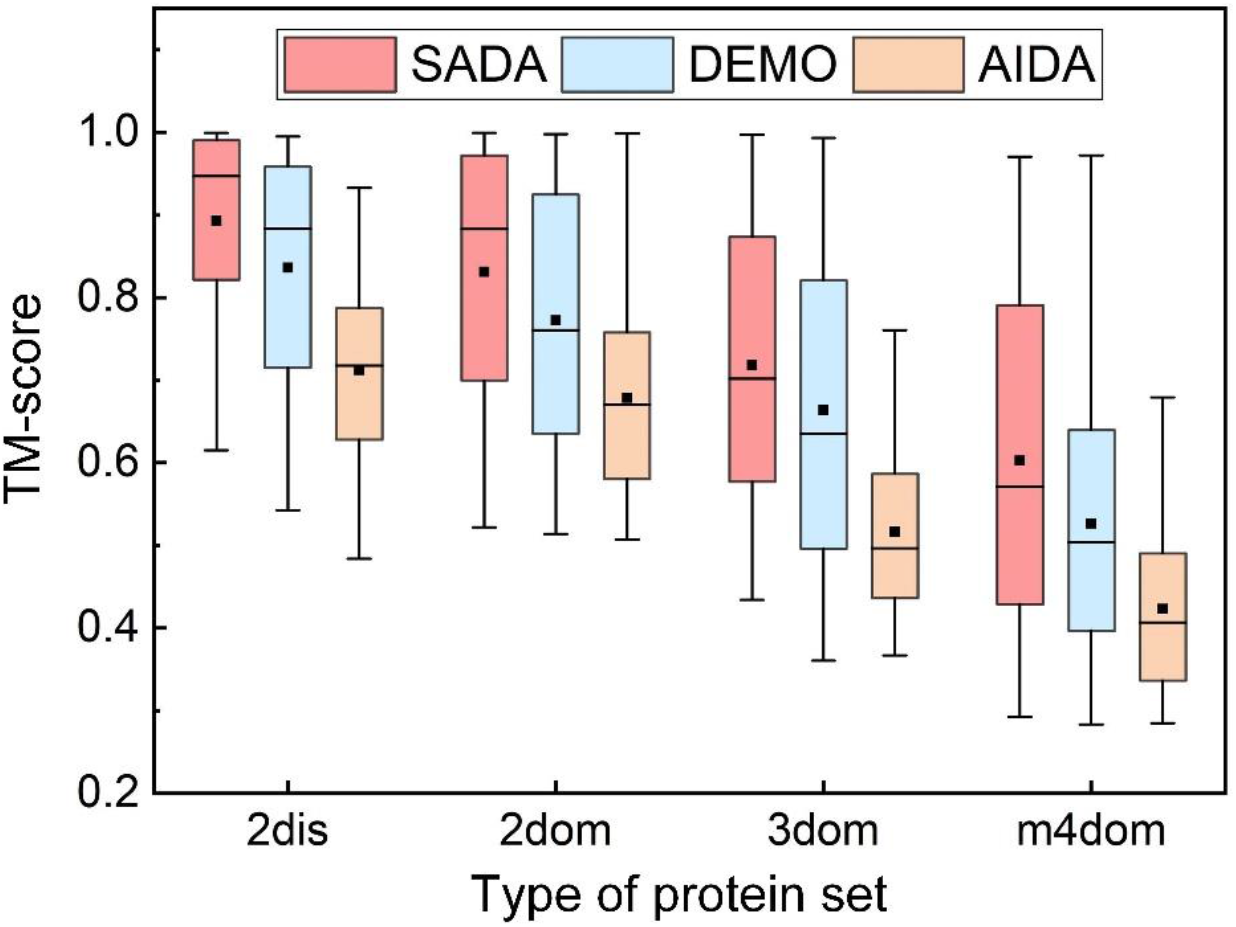
Boxplot for the TM-score of the assembly models by SADA, DEMO and AIDA. The square and horizontal lines in the box represent the mean and median TM-scores, and the horizontal lines on the top and bottom are the maximum and minimum TM-scores, respectively.

Figure 2 shows the average TM-score histograms in separate categories, indicating that SADA assembled more accurate full-chain models for the proteins of different types of domains. As the number of domains increased, the performance of these methods decreased, possibly because the search space of domain assembly simulations increased as the degrees of freedom increased for proteins of more domains. In addition, for SADA, 2dom and 3dom proteins account for the majority in MPDB, and with the increase of the number of domains, high-quality full-chain structural analogues were hard to identify, thus affecting the performance. However, on the 40 m4dom proteins, the average TM-score is 0.60, which is 13.2% higher than that of the second-best method (DEMO). It is because we used GeomNet to capture the distance information between domains, thus compensating for the quality decline of structural analogues. Generally, models of SADA have a higher TM-score than that of other methods, and the overall quality of the multi-domain models is acceptable, with an average TM-score of 0.68 for proteins with 3 or more domains, 74.3% of which had a TM-score ≥ 0.5.

### 3.3 Coverage of multi-domain proteins in the MPDB

On the benchmark set, we compared the coverage of MPDB and that of template library used in DEMO (Zhou *et al*., 2019). Here, the template library used in DEMO is denoted as DEMO-lib. In the MPDB and DEMO-lib, TM-align (Zhang and Skolnick, 2005) was used to calculate the TM-score (Zhang and Skolnick, 2004) between each test protein and the proteins in the two databases, respectively. For each test protein, the average TM-score of the top 10 templates with the highest TM-score was considered as the coverage score of the test protein in the corresponding database.

At a sequence identity cut-off of 30%, the average coverage score of MPDB is 0.57, which is 7.5% higher than that of DEMO-lib (0.53) on the 166 2dom proteins. The average coverage score of MPDB is 0.56, which is 12.0% higher than that of DEMO-lib (0.50) on the 69 3dom proteins. On the m4dom proteins, the average coverage score of MPDB is 0.48, which is 6.7% higher than that of DEMO-lib (0.45). The average coverage score of MPDB is 0.56 on the 81 2dis proteins, which is 5.7% higher than that of DEMO-lib (0.53). The *P*-value between the coverage scores of MPDB and that of DEMO-lib is 5.54E-53, indicating that there is a significant difference between the coverage score of MPDB and DEMO-lib. The head-to-head comparisons between the coverage score of each test protein in DEMO-lib and MPDB are shown in Figure 3. At a sequence identity cut-off of 50%, the average coverage score of MPDB is 0.60, which is 9.1% higher than that of DEMO-lib (0.55) for the 356 test proteins. At a sequence identity cut-off of 70%, the average coverage score of MPDB is 0.61, which is 10.9% higher than that of DEMO-lib (0.55). The detailed coverage score for each test protein and the summaries of different databases are listed in Supplementary Tables S4-S9. At sequence identity cut-off values of 50% and 70%, the head-to-head comparisons between the coverage score of each test protein in DEMO-lib and MPDB are shown in Supplementary Figure S2.

**Fig. 3.**
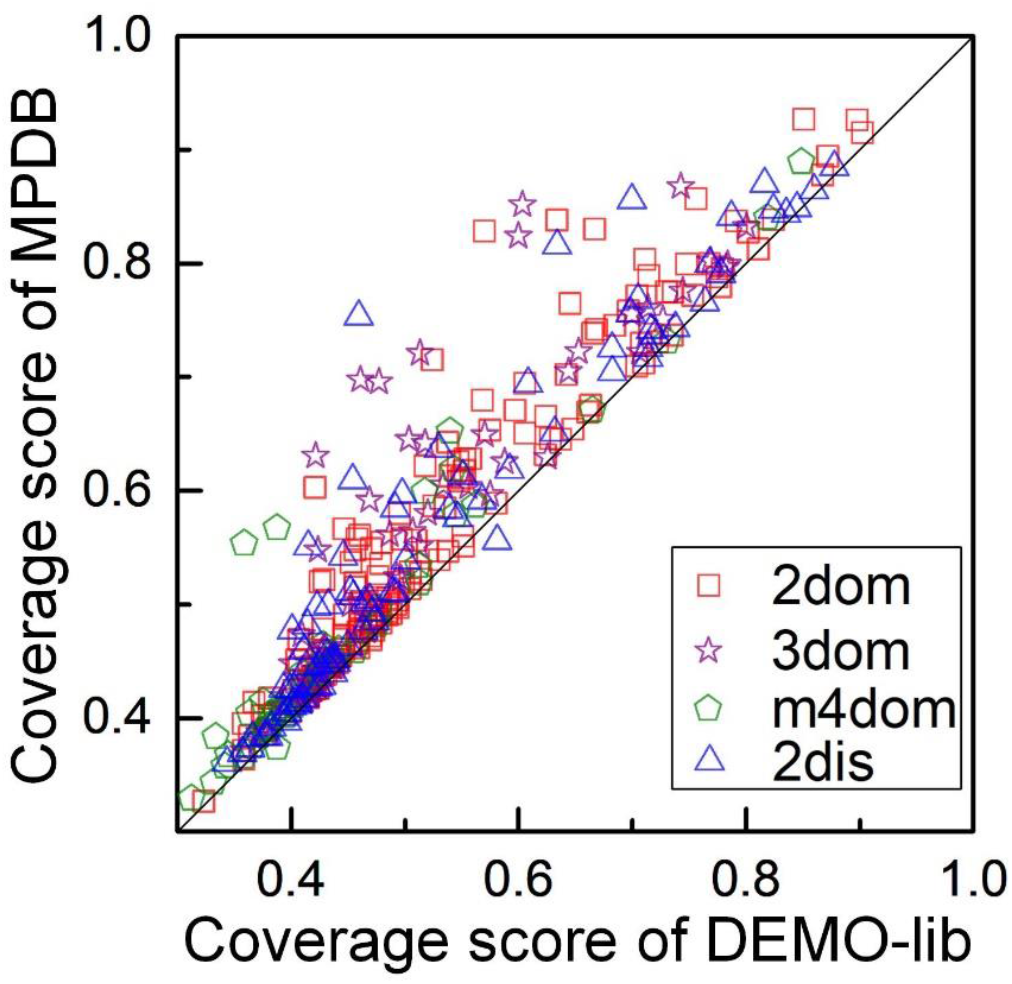
Head-to-head comparison between the coverage score of the test proteins in DEMO-lib and MPDB when the sequence identity between the test protein and the proteins in MPDB and DEMO-lib is less than 30%. The x-axis represents the coverage score of the test proteins in the DEMO-lib. The y-axis represents the coverage score of the test proteins in the MPDB.

These results show that the constructed MPDB covers remarkably more multi-domain protein structures than DEMO-lib, and there is the significantly difference between the coverage score of MPDB and that of DEMO-lib.

### 3.4 Performance of the structural analogue detection

There are some proteins that the sequence identity between them is low, but they have similar topologies. These proteins may be important for structural modelling of multi-domain proteins.

Structural analogue serves as the basis for SADA, and in this study, we tested the performance of the structural analogue detection algorithm. On the 356 test proteins, the proposed structural analogue detection algorithm was used to detect structural analogues of the full-chain according to the individual domain models, and the structural analogue with the highest *LS*_score_ was used to calculate TM-score between the analogue and native structure of target protein. When the sequence identity was less than 30%, the average TM-score of the 356 structural analogues with the highest *LS*_score_ was 0.56, where 192 cases have the highly similar topologies to the native structures with TM-score ≥ 0.5. When the sequence identities were less than 50% and 70%, the average TM-score of the structural analogues is increased to 0.64 and 0.65, respectively. The number of structural analogues with TM-score ≥ 0.5 to the full-chain native structures were 239 and 247, respectively. The detailed TM-score of structural analogues for each test protein at different sequence identity cut-off values are shown in Supplementary Tables S10 and S11. At different sequence identity cut-off values, the average TM-scores of structural analogues are always more than 0.5, indicating that the detected structural analogues can describe the global topological structure of the full-chain for the majority of multi-domain proteins in the test set. For the 356 test proteins, the *LS*_score_ of the top 1 structural analogue is shown in Table S12. Figure 4 shows the distributions between *LS*_score_ and TM-score at a sequence identity cut-off of 30%. At sequence identity cut-off values of 50% and 70%, the distributions between *LS*_score_ and TM-score are shown in Supplementary Figure S3.

**Fig. 4.**
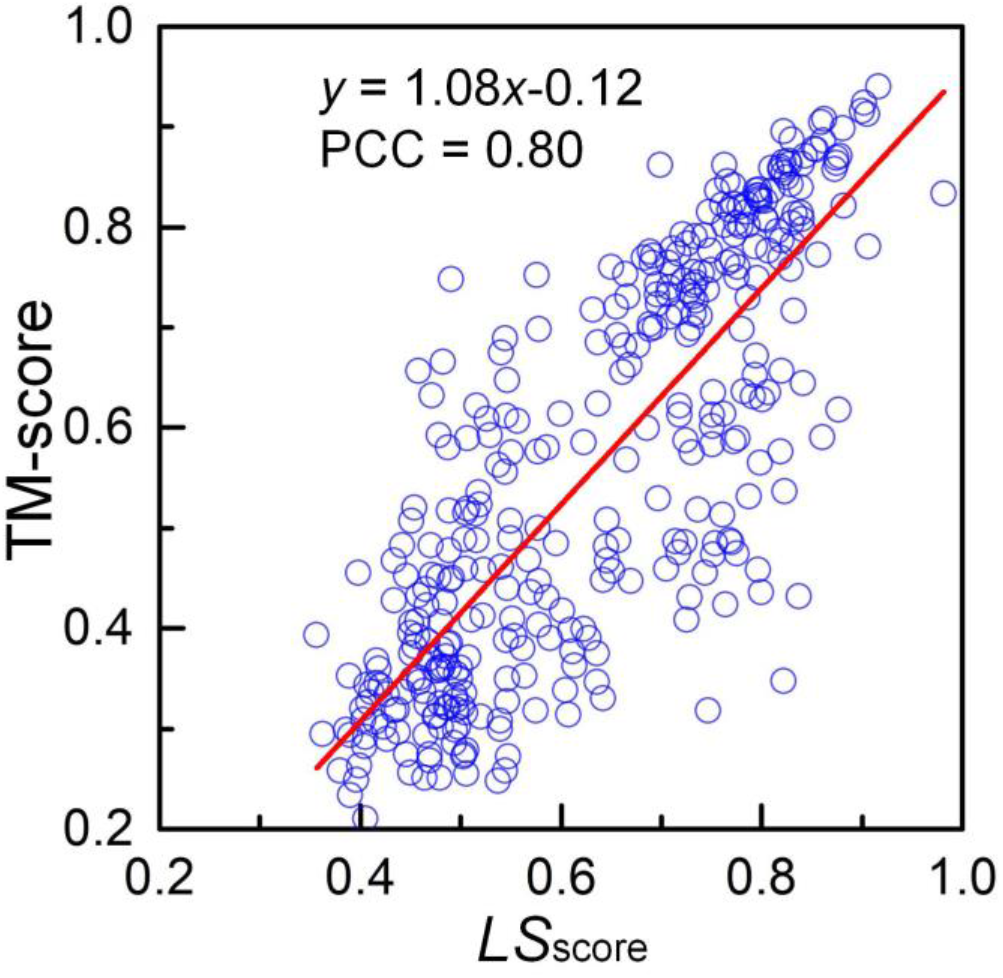
Distribution of *LS*_score_ and TM-score under sequence identity of less than 30%. For each blue circle, the x-axis represents the *LS*_score_ of the structural analogue with the highest *LS*_score_ in the corresponding test protein, and the y-axis represents the TM-score between the native structure of the test protein and the detected structural analogue. The least-squares linear fit and Pearson correlation coefficient (PCC) are listed in the figure.

To further prove the effectiveness of the proposed structural analogue detection algorithm, the Pearson correlation coefficient between *LS*_score_ of the top 1 structural analogue and TM-score of the structural analogue to the native structure was calculated. The Pearson correlation coefficients between the *LS*_score_ and TM-score are 0.80, 0.80 and 0.81 at sequence identity cut-off values of 30%, 50% and 70%, respectively.

The results show that the proposed structural analogue detection algorithm can effectively detect the structural analogues of full-chain and verify that domains often interact in the similar way in the quaternary orientations if the domains have similar tertiary structures.

### 3.5 Effect of using predicted distance information

In SADA, an inter-residue distance potential predicted by GeomNet and the physics-based force field was used to guide domain assembly. The inter-residue distance information was not used in SADA to discuss the effect of the predicted distance information, and experiments were conducted on 356 test proteins. The SADA without using predicted distance information, namely, SADA-w/o-D, was used for comparative experiments to explore the contribution of the distance information predicted by GeomNet.

The domain assembly results of SADA and SADA-w/o-D on the benchmark are shown in Supplementary Table S13 and summarized in Supplementary Table S14. Based on the results, SADA achieves better TM-score than SADA-w/o-D in 275 out of 356 proteins. The average TM-score of SADA is 0.80 on 356 test proteins, which is 8.1% higher than that of SADA-w/o-D (0.74). On 166 2dom proteins and 81 2dis proteins, the average TM-scores of SADA (0.83 for 2dom and 0.89 for 2dis) were 5.1% and 4.7% higher than that of SADA-w/o-D (0.79 for 2dom and 0.85 for 2dis), respectively. Especially, on 69 3dom proteins and 40 m4dom proteins, the average TM-scores of SADA (0.72 for 3dom and 0.60 for m4dom) were 10.8% and 25.0% higher than that of SADA-w/o-D (0.65 for 3dom and 0.48 for m4dom), respectively. For an intuitive comparison of the effect of the predicted distance information, the average TM-score histogram in separate categories is depicted in Figure 5.

**Fig. 5.**
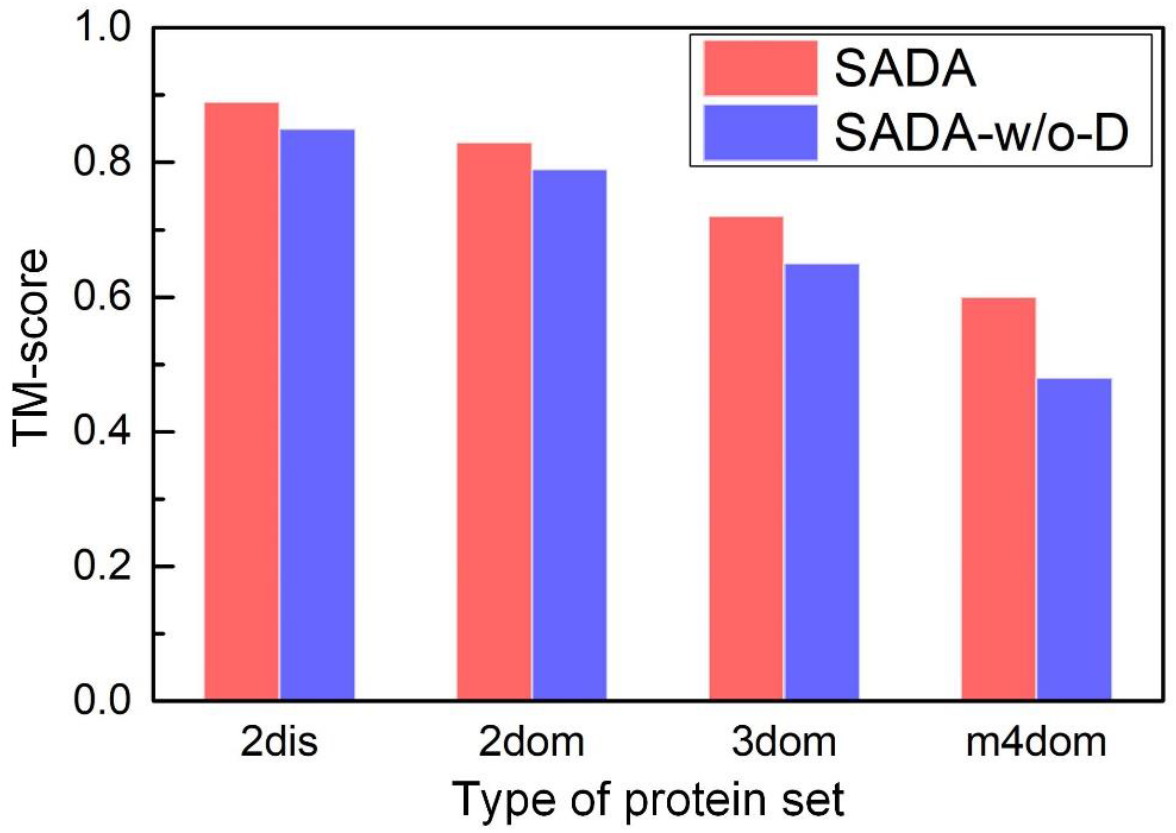
Average TM-score of the final model assembled by SADA and SADA-w/o-D on 356 test proteins.

Based on the results of the above-mentioned comparative experiment, inter-residue distance information predicted by GeomNet can capture the orientation between domains that cannot be obtained by templates, thus improving the accuracy of protein domain assembly.

### 3.6 Complementarity analysis between structural analogue and homologue

Template, as an important part of protein structure modelling, is mainly derived from homologous proteins. However, homologous templates for individual domains can often be detected, but templates that can be used to model the entire query protein are often unavailable. Therefore, structural analogues are suitable for multi-domain protein modelling in some cases, because some proteins are formed by duplication, divergence and recombination of domains.

Among the 356 test proteins, we analysed the quality of the detected structural analogues and the homologues searched by JackHMMER (Eddy, 1998). Here, the parameters of JackHMMER were set as default, and each domain model of test proteins was a native model. When the sequence identity cut-offs were 30%, 50% and 70%, the structural analogue with the highest *LS*_score_ and the top 1 homologue were used to calculate the TM-score between the analogue (homologue) and native structure of target protein, respectively. The detailed results of each protein are shown in Supplementary Tables S10 and S15, respectively. At a sequence identity cut-off of 30%, the head-to-head analysis between the TM-score of the analogue with the highest *LS*_score_ and that of top 1 homologue are shown in Figure 6. At a sequence identity cut-off of 30%, the TM-score of the structural analogues is greater than or equal to that of homologues in 169 out of the 256 test proteins, where the 100 out of the 356 test proteins that JackHMMER cannot search for homologues in MPDB. Especially for the 100 proteins without homologous templates, there are 24 structural analogues with TM-score ≥ 0.5. The results are summarised in Supplementary Table S16.

**Fig. 6.**
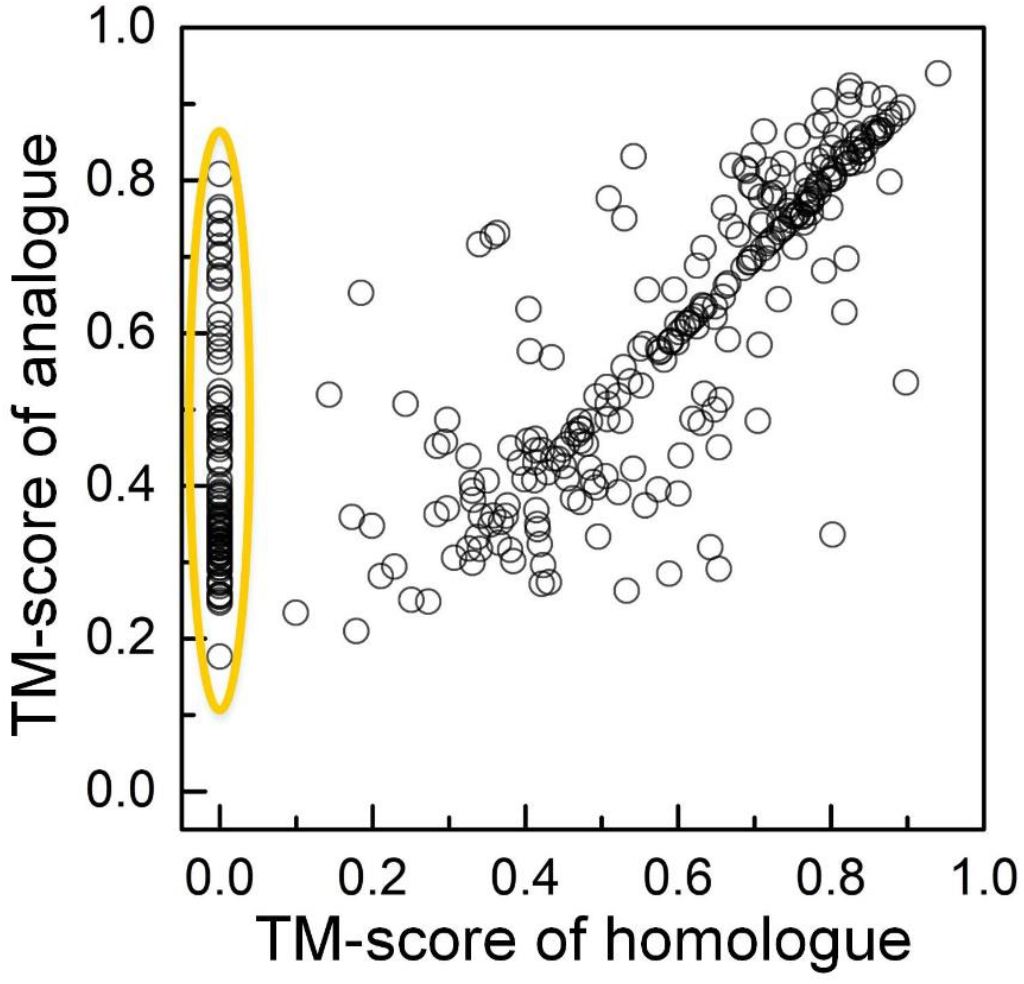
Head-to-head analysis between the TM-score of analogue and that of homologue under the sequence identity cut-off of 30%. For each black circle, the x-axis represents the TM-score between the native structure of the test protein and the top 1 homologous template, and the y-axis represents the TM-score between the native structure of the test protein and the structural analogue with the highest *LS*_score_. The black circles in the yellow area represent that the homologous templates of these test proteins cannot be detected by JackHMMER.

At a sequence identity cut-off of 50%, JackHMMER cannot search for homologues in MPDB in 71 out of the 356 test proteins, where the TM-score of the structural analogues is greater than or equal to that of homologues in 213 out of the 285 test proteins. The results are summarised in Supplementary Table S17. Among the 285 test proteins, the average TM-score of the top 1 homologue is 0.66 and the average TM-score of the analogues with the highest *LS*_score_ is 0.68. Among the 71 test proteins, 22 structural analogues have TM-score ≥ 0.5. At a sequence identity cutoff of 70%, JackHMMER cannot search for homologues in MPDB in 65 of the 356 test proteins, where the TM-score of the structural analogues is greater than or equal to that of homologues in 218 out of the 291 test proteins. The results are summarised in Supplementary Table S18. On the 291 test proteins, the average TM-score of the top 1 homologue is 0.68 and the average TM-score of the analogues with the highest *LS*_score_ is 0.70. On the 65 test proteins, 20 structural analogues have TM-score ≥ 0.5. The head-to-head analysis at identity cut-offs of 50% and 70% are shown in Supplementary Figure S4.

These results show that the structural analogues and homologues are complementary. For the test proteins in which the JackHMMER cannot search for homologues, the proposed structural analogues detection algorithm can detect some structural analogues with a TM-score ≥ 0.5.

### 3.7 Assembly multi-domain proteins using analogue and homologue

To further study the difference of analogues and homologues on the accuracy of domain assembly, we used SADA-w/o-D to assemble the full-chain structure of multi-domain proteins according to the homologous templates and analogous templates, respectively. In this study, SADA-w/o-D was used to assemble domains, because the influence of distance information predicted by deep learning needs to be excluded.

At a sequence identity cut-off of 30%, the initial models were generated based on the analogue with the highest *LS*_score_ and top 1 homologue, respectively. The final results are shown in Supplementary Table S19. Among the 256 test proteins with homologues detected by JackHMMER, the average TM-scores were 0.75 and 0.76 for the full-chain models generated by SADA-w/o-D using homologues and analogues, respectively. SADA-w/o-D using analogues obtained full-chain models with TM-score ≥ 0.5 in 218 out of 256 and accounts for 85.2% of the 256 proteins. SADA-w/o-D using homologues obtained models with TM-score ≥ 0.5 in 214 out of 256 and accounts for 83.6% of the 256 proteins. The full-chain models generated by SADA-w/o-D using analogue are better than that using homologue in 132 out of the 256 test proteins. Especially, on the 100 test proteins without homologous templates, 88 full-chain models have a TM-score ≥ 0.5. For an intuitive analysis of the quality of full-chain models assembled by using different template on benchmark target proteins, the head-to-head comparison is shown in Figure 7.

**Fig. 7.**
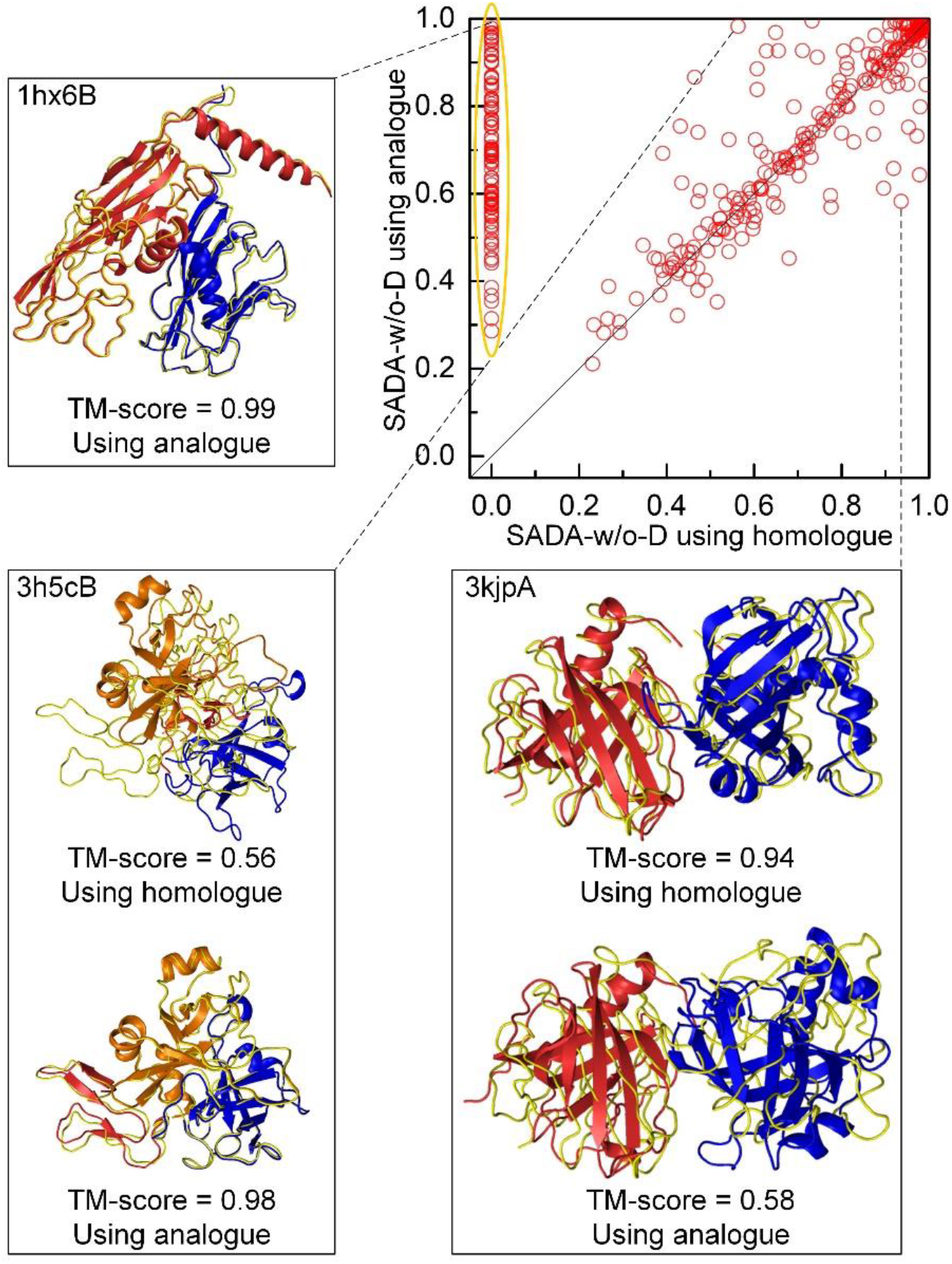
Head-to-head comparison of the models assembled by SADA-w/o-D using homologue and analogue in TM-score. The red circles in the yellow region represent that the homologous templates of these test proteins cannot be detected by JackHMMER. The different colors in protein structures represent different domains, and the yellow lines represent native structures.

Three illustrative examples are shown from Figure 7. For the 2dom protein 1hx6B, JackHMMER cannot detect homologous template at a sequence identity cut-off of 30%, while SADA-w/o-D assembled the full-chain model with TM-score of 0.99 by using analogous template. For the 3dom protein 3h5cB, the full-chain model with TM-score of 0.56 was generated using homologous template, while SADA-w/o-D using analogous template assembled the full-chain model with TM-score of 0.98. For the 2dom protein 3kjpA, SADA-w/o-D using homologous and analogous template assembled the full-chain model with TM-scores of 0.94 and 0.58, respectively.

These results further demonstrate that structural analogues and homologues are complementary, and the combination of analogues and homologues may improve the modelling accuracy of multi-domain proteins.

### 3.8 Assembly human proteins predicted by AlphaFold2

Although AlphaFold2 predicted the structures of many protein targets at or near experimental resolution, for some multi-domain proteins, the quality of its predicted structure may be further improved by SADA. In this section, we reassembled 20 human multi-domain proteins selected from AlphaFold DB, whose full-chain models predicted by AlphaFold2 were all less than 0.80 in TM-score (Tunyasuvunakool *et al*., 2021). Here, we conducted two sets of experiments respectively. In the first set of experiments, the individual domain models for SADA assembly were decomposed from the full-chain structures of AlphaFold2 (deposited in AlphaFold DB). In the second set of experiments, the individual domain models for SADA assembly were independently predicted by AlphaFold2. In both sets of experiments, structural analogues with a sequence identity > 70% to the target were excluded. The detailed results of each protein are shown in Supplementary Table S20, and the head-to-head comparison between them are shown in Figure 8.

**Fig. 8.**
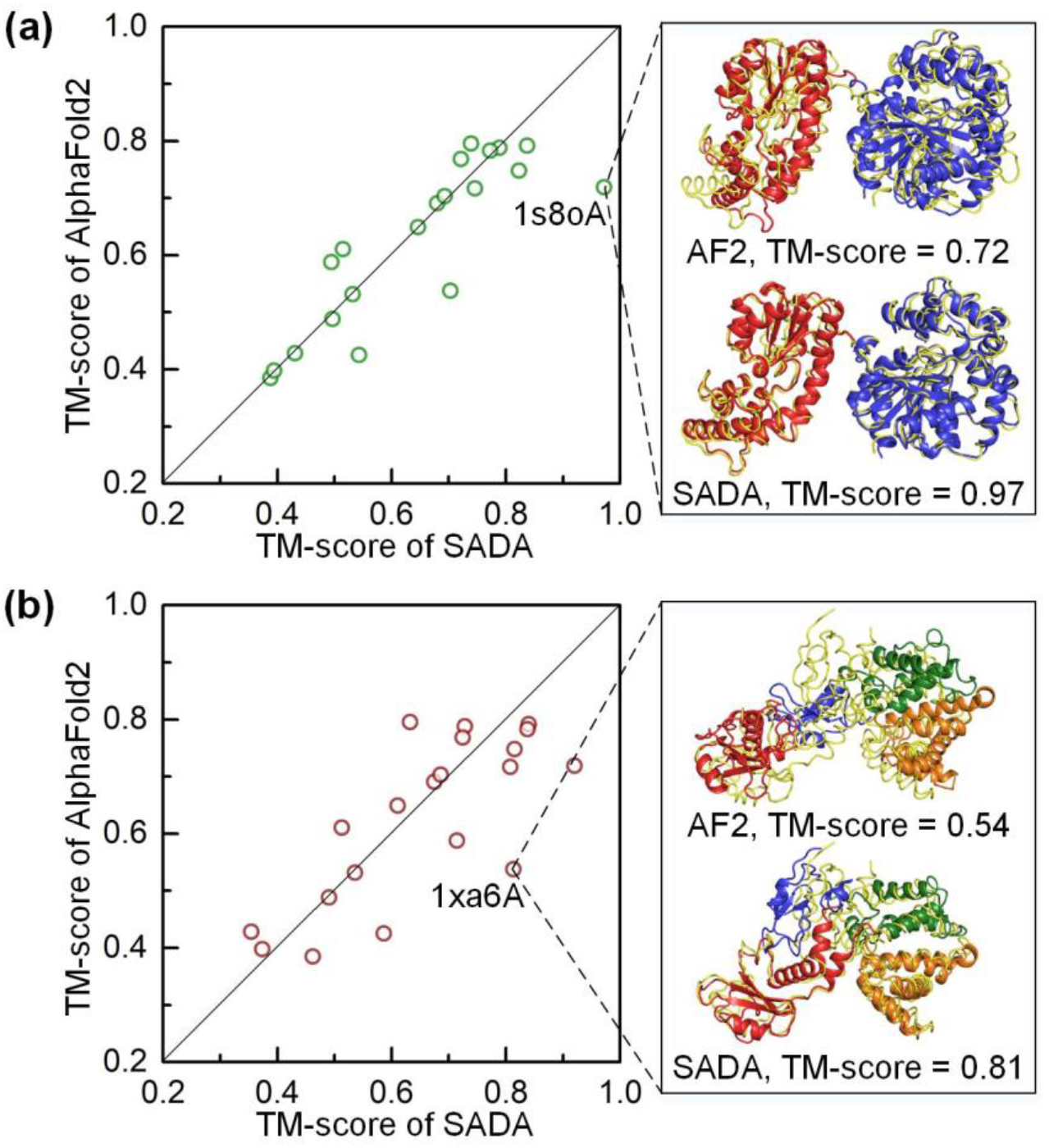
The head-to-head comparison between full-chain structures of AlphaFold2 and full-chain structures assembled by SADA. (a) The domain models for SADA were decomposed from the full-chain structures of AlphaFold2. (b) The domain models for SADA were predicted by AlphaFold2. The different colors represent different domains and the yellow lines are native structures.

In the first set of experiments, the TM-score of 10 out of 20 full-chain structures was increased after SADA reassembly, and the average TM-score increased from 0.63 to 0.65. In the second set of experiments, the TM-score of 11 out of 20 full-chain structures was increased after SADA reassembly, and the average TM-score increased from 0.63 to 0.66. In Figure 8, we selected 2 representative examples.

For the protein 1s8oA in Figure 8(a), it was composed of two domains, where the TM-scores for the models decomposed from the full-chain structures of AlphaFold2 for domains 1 and 2 were 0.98 and 0.97, respectively, but the full-length structure achieved a TM score of 0.72. However, the TM-score of the full-chain model generated by SADA achieved 0.97. For the protein 1xa6A in Figure 8(b), it was composed of four domains, where the TM-scores for the models predicted by AlphaFold2 for domains 1, 2, 3 and 4 were 0.93, 0.90, 0.93 and 0.95, respectively, and full-chain model assembled by SADA achieved a TM-score of 0.81. However, the full-chain structure of AlphaFold2 achieved a TM-score of 0.54, although the TM-scores for the models decomposed from the full-chain structures of AlphaFold2 for domains 1, 2, 3 and 4 were 0.92, 0.88, 0.93 and 0.95, respectively. This may be because 46 residues are missing in the native structure of domain 2, so the structure of these 46 residues in the predicted full-chain model affects the orientation between domains, resulting in poor quality of the full-chain model. The domain models predicted by AlphaFold2 may have a closer structure to the real structure, especially on domain 2. The structures of domain 2 are shown in Supplementary Figure S5. Thus, full-chain model assembled by SADA achieved a TM-score of 0.81, where the domain models were predicted by AlphaFold2.

For some multi-domain proteins, the correct orientation between domains may not be captured by AlphaFold2, and the accuracy of the full-chain modelling may sharply decline due to partial structure prediction errors. These results further indicate that the domain assembly may be an effective strategy for improving the accuracy of full-chain modelling.

## 4 Conclusion

We developed a structural analogue-based protein structure domain assembly method assisted by deep learning, named SADA. In SADA, a multi-domain protein structure database was constructed using DomainParser and the domain knowledge defined in CATH and SCOPe databases, named MPDB. Based on MPDB, a structural analogue detection algorithm was proposed, to identify the structural analogues of full-chain from MPDB through structural alignment of individual domain models. Based on the detected analogue, the initial full-chain model was generated. Under the guidance of the physics-based force field and an inter-residue distance potential predicted by GeomNet, domain assembly was simulated by two-stage differential evolution algorithm to generate final full-chain model. In the exploration stage, some superior topology structures were generated. In the exploitation stage, the local position was optimised based on superior topology structures. Results on 356 tested proteins show that the proposed SADA significantly outperforms most state-of-the-art domain assembly methods. In addition, SADA was also applied to assemble 20 multi-domain human proteins from the AlphaFold2 prediction, and the quality of the full-chain models of half of the 20 proteins improved. Among them, some cases achieved significantly better models. The proposed protein domain assembly method may be an effective complement to the end-to-end approaches.

In this study, the Pearson correlation coefficient between the local similarity score *LS*_score_ and TM-score is 0.80 at sequence identity cut-off of 30%. The results indicate that domains often interact in the similar way in the quaternary orientations if they have similar tertiary structures. Moreover, the analysis of homologues searched by JackHMMER and structural analogues detected by the proposed algorithm shows that they are complementary. If more suitable templates can be selected from homologues and analogues, the accuracy of multi-domain protein structure modelling can be further improved. Meanwhile, similar idea of SADA may be extended to the detection of structural analogue of protein complex, which may contribute to the protein complex modelling, and this is also the next direction of our group.

## 5 Acknowledgments

We thank Zhongze Yu and Fenqi Ge for helpful discussions. This work has been supported by the “New Generation Artificial Intelligence” major project of Science and Technology Innovation 2030 of the Ministry of Science and Technology of the People’s Republic of China [No. 2021ZD0150100], the National Nature Science Foundation of China [No. 62173304, 61773346], and the Key Project of Zhejiang Provincial Natural Science Foundation of China [No. LZ20F030002].

## References

Baek, M. et al. (2021) Accurate prediction of protein structures and interactions using a three-track neural network. Science, 373, 871–876.

Chandonia, J.M. et al. (2017) SCOPe: Manual Curation and Artifact Removal in the Structural Classification of Proteins-extended Database. Journal of Molecular Biology, 429, 348–355.

Du, Z.Y. et al. (2021) The trRosetta server for fast and accurate protein structure prediction. Nature Protocol, 16, 5634–5651.

Eddy, S.R. (1998) Profile hidden Markov models. Bioinformatics, 14, 755–763.

Fu, L. et al. (2012) CD-HIT: Accelerated for clustering the next-generation sequencing data. Bioinformatics, 28, 3150–3152.

Javier, C.D. et al. (2020) Deep learning enables the design of functional de novo antimicrobial proteins. bioRxiv, DOI: https://doi.org/10.1101/2020.08.26.266940.

Jones, D.T. and Thornton, J.M. (2022) The impact of AlphaFold2 one year on. Nature Methods, 19, 15–20.

Jumper, J. et al. (2021) Highly accurate protein structure prediction with AlphaFold. Nature, 596, 583–589.

Lam, S.D. et al. (2016) Gene3D: expanding the utility of domain assignments. Nucleic Acids Research, 44, D404–D409.

Liu, J. et al. (2022) De novo protein structure prediction by incremental inter-residue geometries prediction and model quality assessment using deep learning. bioRxiv, DOI: 10.1101/2022.01.11.475831.

Mirdita, M. et al. (2017) Uniclust databases of clustered and deeply annotated protein sequences and alignments. Nucleic Acids Research, 45, D170–D176.

Orengo, C.A. et al. (1997) CATH-a hierarchic classification of protein domain structures. Structure, 5, 1093–1109.

Pearce, R. and Zhang, Y. (2021) Toward the solution of the protein structure prediction problem. Journal of Biological Chemistry, 297, 100870.

Steinegger, M. et al. (2019) Protein-level assembly increases protein sequence recovery from metagenomic samples manyfold. Nature methods, 16, 603–606.

Steinegger, M. et al. (2019) HH-suite3 for fast remote homology detection and deep protein annotation. BMC bioinformatics, 20, 1–15.

Su, H. et al. (2021) Improved protein structure prediction using a new multi-scale network and homologous templates. Advanced Science, 8, 2102592–2102602.

Tunyasuvunakool, K. et al. (2021) Highly accurate protein structure prediction for the human proteome. Nature, 596, 590–596.

Wang, G.L. and Dunbrack, R.L. (2003) PISCES: a protein sequence culling server. Bioinformatics, 19, 1589–1591.

Wollacott, A.M. et al. (2007) Prediction of structures of multidomain proteins from structures of the individual domains. Protein Science, 16, 165–175.

Xu, D. et al. (2015) AIDA: Ab initio domain assembly for automated multidomain protein structure prediction and domain-domain interaction prediction. Bioinformatics, 31, 2098–2105.

Xu, J.B. (2019) Distance-based protein folding powered by deep learning. Proceedings of the National Academy of Sciences, 116, 16856–16865.

Xu, J.B. and Wang, S. (2019) Analysis of distance-based protein structure prediction by deep learning in CASP13. Proteins, Structure, Function, and Bioinformatics, 87, 1069–1081.

Xu, Y. et al. (2000) Protein domain decomposition using a graph-theoretic approach. Bioinformatics, 16, 1091–1104.

Yang, J.Y. et al. (2020) Improved protein structure prediction using predicted interresidue orientations. Proceedings of the National Academy of Sciences, 117, 1496–1503.

Zhang, Y. and Skolnick, J. (2004) Scoring function for automated assessment of protein structure template quality. Proteins, 57, 702–710.

Zhang, Y. and Skolnick, J. (2005) TM-align: a protein structure alignment algorithm based on the TM-score. Nucleic Acids Research, 33, 2302–2309.

Zheng, W. et al. (2021) Protein structure prediction using deep learning distance and hydrogen-bonding restraints in CASP14. Proteins, DOI: 10.1002/prot.26193.

Zhou, X.G. et al. (2019) Assembling multidomain protein structures through analogous global structural alignments. Proceedings of the National Academy of Sciences, 116, 15930–15938.

Zhou, X.G. et al. (2022) Progressive and accurate assembly of multi-domain protein structures from cryo-EM density maps. Nature Computational Science, DOI: 10.1101/2020.10.15.340455, in press.

