## Supplementary Information for "Structural analogue-based protein structure domain assembly assisted by deep learning"

#### Supplementary Figures

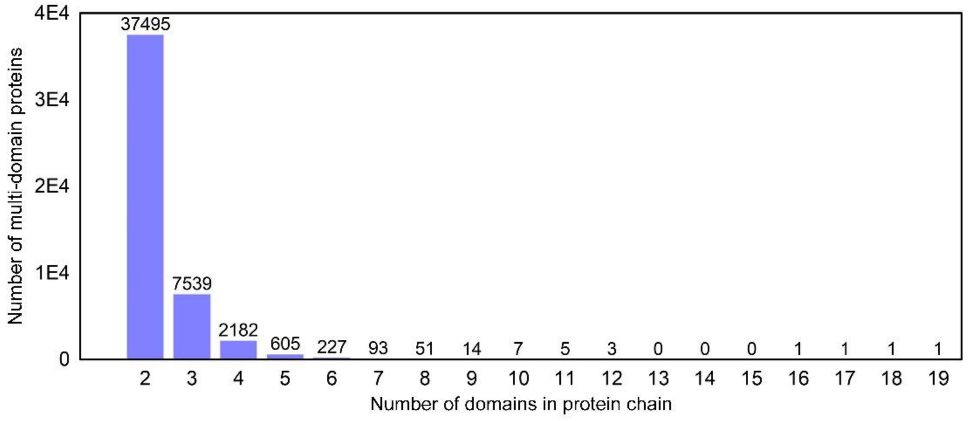


**Fig. S1.** Distribution of different multi-domain proteins in MPDB.

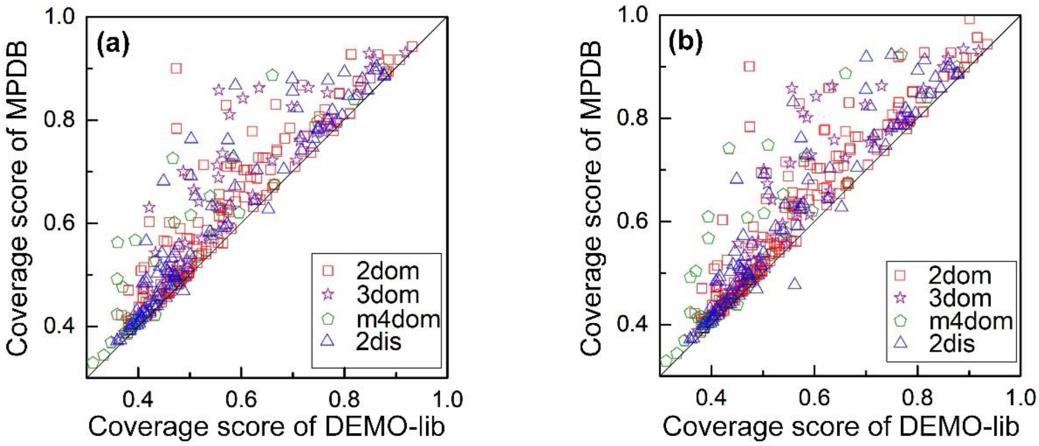


**Fig. S2.** Head-to-head comparison between the coverage score of the test proteins in DEMO-lib and MPDB. (a) and (b) represent the sequence identity between the test protein and the proteins in MPDB and DEMO-lib is less than 50% and 70%, respectively.

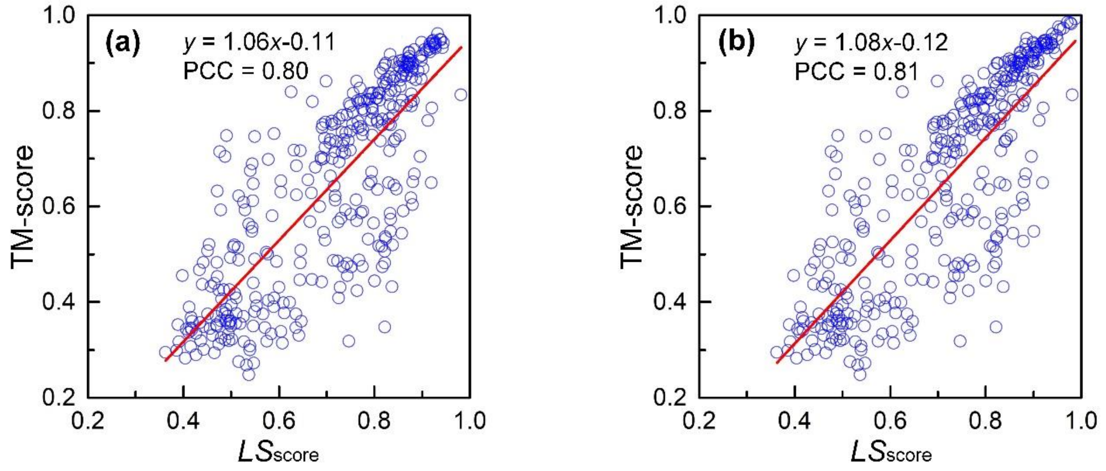


**Fig. S3.** Distribution of $LS_{score}$ and TM-score. (a) and (b) represent the distribution $LS_{score}$ and TM-score under sequence identity less than 50% and 70%, respectively.

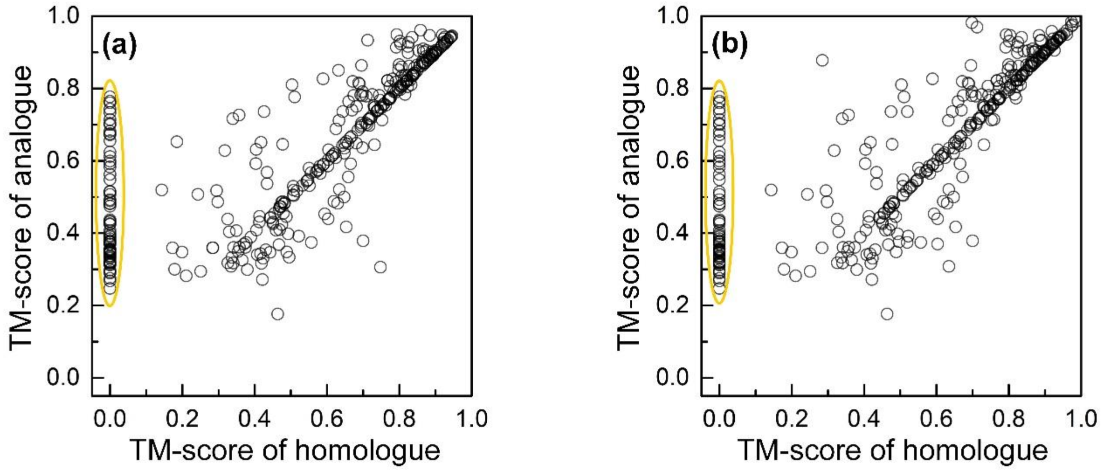


**Fig. S4.** Head-to-head analysis between the TM-score of analogues and that of homologues. (a) and (b) represent the results under sequence identity less than 50% and 70%, respectively.

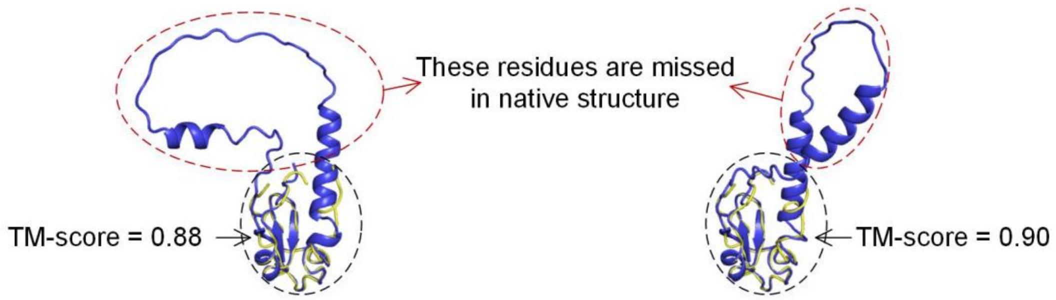


**Fig. S5.** Structure of domain 2 of 1xa6A. Left is the domain structure decomposed from the full-chain structures of AlphaFold2. Right is the domain structure predicted by AlphaFold2. The blue color represents the predicted structure, and the yellow lines are native structures.

#### Supplementary Tables

**Table S1.** Protein length and number of domains on each test protein. #Domains represents the number of domains. 2dis represents the protein with 2 discontinuous domains.

| PDB ID | Length | #Domains | PDB ID | Length | #Domains | PDB ID | Length | #Domains |
|---|---|---|---|---|---|---|---|---|
| 1cjyA | 630 | 2 | 3rwxA | 239 | 2 | 1kfqA | 567 | 4 |
| 1efdN | 262 | 2 | 3sb4A | 323 | 2 | 1ldjA | 725 | 7 |
| 1fjrA | 188 | 2 | 3swjA | 246 | 2 | 1nyqB | 645 | 4 |
| 1g87B | 613 | 2 | 3t58B | 509 | 2 | 1ug9A | 1019 | 4 |
| 1hx6B | 396 | 2 | 3t7jA | 237 | 2 | 1z1wA | 780 | 4 |
| 1iwaA | 441 | 2 | 3u07C | 382 | 2 | 2au3A | 403 | 4 |
| 1m5qH | 123 | 2 | 3u0oB | 335 | 2 | 2ii2A | 304 | 4 |
| 1mkfA | 371 | 2 | 3u9gA | 217 | 2 | 2olsA | 725 | 4 |
| 1mkmB | 247 | 2 | 3ub1D | 251 | 2 | 2ra1A | 413 | 5 |
| 1nh2D | 102 | 2 | 3uitD | 257 | 2 | 2v5dA | 722 | 4 |
| 1pprM | 312 | 2 | 3uo3A | 184 | 2 | 2xt6A | 1055 | 4 |
| 1prrA | 173 | 2 | 3v7oB | 203 | 2 | 2zpaB | 651 | 4 |
| 1q19A | 500 | 2 | 3vr8B | 249 | 2 | 3apoA | 688 | 7 |
| 1qwrA | 321 | 2 | 3wkuA | 405 | 2 | 3b43A | 569 | 6 |
| 1r71B | 116 | 2 | 3zvmA | 385 | 2 | 3gf5B | 380 | 7 |
| 1rh1A | 490 | 2 | 4acoA | 452 | 2 | 3hjlA | 321 | 4 |
| 1rktA | 207 | 2 | 4ap5A | 373 | 2 | 3kq4B | 457 | 4 |
| 1s6lA | 181 | 2 | 4axdA | 437 | 2 | 3kwlA | 493 | 4 |
| 1sp3A | 436 | 2 | 4bfiB | 202 | 2 | 3ob8A | 1024 | 5 |
| 1vz6A | 365 | 2 | 4bt9B | 228 | 2 | 3opfB | 493 | 4 |
| 1w3aA | 312 | 2 | 4cczA | 611 | 2 | 3p53A | 494 | 4 |
| 1wv3A | 181 | 2 | 4d0nB | 371 | 2 | 3pvlA | 598 | 5 |
| 1x7pA | 265 | 2 | 4d1iG | 544 | 2 | 3r05A | 1008 | 7 |
| 1x9yA | 346 | 2 | 4dj3A | 298 | 2 | 3ubhA | 409 | 4 |
| 1y11A | 356 | 2 | 4dqaA | 358 | 2 | 3w2wA | 618 | 4 |
| 1yiqA | 684 | 2 | 4eo3A | 328 | 2 | 3zniA | 391 | 4 |
| 1zbuB | 289 | 2 | 4eogA | 460 | 2 | 4aimA | 698 | 4 |
| 1ze1A | 308 | 2 | 4etxA | 300 | 2 | 4ak1A | 612 | 6 |
| 2ablA | 163 | 2 | 4fguA | 407 | 2 | 4aq1A | 721 | 6 |
| 2ahvA | 512 | 2 | 4fkcA | 370 | 2 | 4fe9A | 450 | 4 |
| 2bkpA | 202 | 2 | 4fxkC | 290 | 2 | 4h2aA | 708 | 4 |
| 2c1yA | 231 | 2 | 4gbyA | 475 | 2 | 4i5sB | 395 | 4 |
| 2cxcA | 138 | 2 | 4ggmX | 270 | 2 | 4iggB | 771 | 6 |
| 2d1cA | 495 | 2 | 4gslA | 598 | 2 | 4j9vA | 456 | 4 |
| 2d7iA | 536 | 2 | 4gyjA | 618 | 2 | 4k3bA | 772 | 6 |
| 2e9hA | 157 | 2 | 4h3tA | 518 | 2 | 4kwuA | 1030 | 6 |
| 2e9xB | 175 | 2 | 4hmoA | 310 | 2 | 4m00A | 495 | 4 |
| 2evrA | 240 | 2 | 4ie6A | 404 | 2 | 1bp1A | 456 | 2dis |
| 2ew9A | 149 | 2 | 4l5gA | 162 | 2 | 1cjsA | 212 | 2dis |

| | | | | | | | | |
|---|---|---|---|---|---|---|---|---|
| 2fd5A | 180 | 2 | 4lpqA | 206 | 2 | 1ck1A | 239 | 2dis |
| 2gh8A | 544 | 2 | 4m8rA | 398 | 2 | 1ecrA | 305 | 2dis |
| 2gt1A | 323 | 2 | 4n06B | 340 | 2 | 1f5qD | 248 | 2dis |
| 2gzaC | 336 | 2 | 4nj5A | 482 | 2 | 1fa9A | 833 | 2dis |
| 2hjqA | 99 | 2 | 4opaB | 197 | 2 | 1gu7A | 364 | 2dis |
| 2hwjA | 184 | 2 | 4qkuB | 427 | 2 | 1itwA | 740 | 2dis |
| 2ijd1 | 644 | 2 | 4up9A | 514 | 2 | 1jkiA | 525 | 2dis |
| 2iu7A | 159 | 2 | 4w7sA | 444 | 2 | 1n80A | 328 | 2dis |
| 2iw2A | 479 | 2 | 1bf2A | 750 | 3 | 1nzjA | 273 | 2dis |
| 2jz4A | 299 | 2 | 1bhgA | 611 | 3 | 1qhdA | 397 | 2dis |
| 2kdyA | 261 | 2 | 1f7uA | 606 | 3 | 1qz9A | 404 | 2dis |
| 2kn4A | 158 | 2 | 1fx7A | 230 | 3 | 1sb7B | 340 | 2dis |
| 2mbgA | 265 | 2 | 1griA | 211 | 3 | 1vk1A | 223 | 2dis |
| 2nsfA | 240 | 2 | 1h88C | 152 | 3 | 1vrmA | 304 | 2dis |
| 2nykA | 235 | 2 | 1m8pB | 571 | 3 | 1xvuA | 402 | 2dis |
| 2o6yA | 514 | 2 | 1ni5A | 428 | 3 | 1yy3A | 315 | 2dis |
| 2owbA | 262 | 2 | 1q25A | 428 | 3 | 1z87A | 263 | 2dis |
| 2qfiA | 286 | 2 | 1uzjA | 162 | 3 | 2a1sC | 393 | 2dis |
| 2qp2A | 513 | 2 | 1zpuA | 504 | 3 | 2a3lA | 616 | 2dis |
| 2qygA | 429 | 2 | 1zy9A | 522 | 3 | 2bt1A | 254 | 2dis |
| 2r5wB | 345 | 2 | 2b5uA | 465 | 3 | 2bydA | 283 | 2dis |
| 2uu7A | 370 | 2 | 2ewfA | 574 | 3 | 2c43A | 280 | 2dis |
| 2w4bA | 455 | 2 | 2piaA | 321 | 3 | 2dfyC | 158 | 2dis |
| 2x7iA | 296 | 2 | 2r7dA | 452 | 3 | 2dlaA | 222 | 2dis |
| 2x8kC | 243 | 2 | 2uwnA | 187 | 3 | 2g3pA | 201 | 2dis |
| 2yilA | 131 | 2 | 2v0nA | 459 | 3 | 2gg6A | 445 | 2dis |
| 2yrqA | 173 | 2 | 2vgmA | 354 | 3 | 2gsyE | 433 | 2dis |
| 2zxcA | 643 | 2 | 2wqrB | 322 | 3 | 2gzoA | 187 | 2dis |
| 3a1iA | 508 | 2 | 2y25B | 314 | 3 | 2j2cA | 470 | 2dis |
| 3a45A | 288 | 2 | 2yk0A | 698 | 3 | 2kfwA | 196 | 2dis |
| 3a56A | 291 | 2 | 2zzqA | 469 | 3 | 2l9yA | 167 | 2dis |
| 3ajvA | 174 | 2 | 3bt1U | 273 | 3 | 2ntyB | 337 | 2dis |
| 3aqkA | 390 | 2 | 3c1yA | 349 | 3 | 2r3vA | 390 | 2dis |
| 3arbA | 274 | 2 | 3cw2C | 251 | 3 | 2r58A | 212 | 2dis |
| 3aujG | 139 | 2 | 3f83A | 508 | 3 | 2w4mA | 250 | 2dis |
| 3b2zF | 293 | 2 | 3fc3A | 190 | 3 | 2x0cA | 308 | 2dis |
| 3b7wA | 543 | 2 | 3gbgA | 260 | 3 | 2y51A | 529 | 2dis |
| 3bt3A | 129 | 2 | 3h5cB | 263 | 3 | 2yb0E | 264 | 2dis |
| 3c4tA | 241 | 2 | 3ibjA | 661 | 3 | 2z86C | 603 | 2dis |
| 3craA | 239 | 2 | 3ippB | 420 | 3 | 3afoA | 360 | 2dis |
| 3d30A | 212 | 2 | 3jymB | 307 | 3 | 3bu2A | 189 | 2dis |
| 3eo5A | 169 | 2 | 3kbgA | 187 | 3 | 3cvzA | 248 | 2dis |
| 3errA | 527 | 2 | 3mc8A | 257 | 3 | 3dupA | 279 | 2dis |
| 3g79A | 475 | 2 | 3npfA | 306 | 3 | 3eswA | 332 | 2dis |

| | | | | | | | | |
|---|---|---|---|---|---|---|---|---|
| 3h2tA | 326 | 2 | 3orjA | 415 | 3 | 3eukH | 459 | 2dis |
| 3hcsA | 157 | 2 | 3plaA | 375 | 3 | 3fi7A | 177 | 2dis |
| 3hyiA | 286 | 2 | 3qe9Y | 345 | 3 | 3fvvA | 223 | 2dis |
| 3i2dA | 279 | 2 | 3qjoA | 491 | 3 | 3gmsA | 331 | 2dis |
| 3iam2 | 179 | 2 | 3qphA | 335 | 3 | 3hzzB | 447 | 2dis |
| 3ifrA | 483 | 2 | 3qyeA | 317 | 3 | 3m1uA | 410 | 2dis |
| 3isqA | 384 | 2 | 3rimA | 697 | 3 | 3mw8A | 233 | 2dis |
| 3j7aK | 129 | 2 | 3rrpA | 455 | 3 | 3mwcA | 388 | 2dis |
| 3k1rA | 193 | 2 | 3soaA | 436 | 3 | 3nsjA | 530 | 2dis |
| 3k2iA | 411 | 2 | 3tixD | 424 | 3 | 3ntkA | 169 | 2dis |
| 3kh5A | 279 | 2 | 3tp9A | 467 | 3 | 3oaaG | 284 | 2dis |
| 3kjpA | 293 | 2 | 3ua3A | 648 | 3 | 3ptyA | 284 | 2dis |
| 3kt1A | 557 | 2 | 3uj0A | 502 | 3 | 3rfyA | 356 | 2dis |
| 3ktmE | 171 | 2 | 3vn4A | 375 | 3 | 3seoB | 226 | 2dis |
| 3kzwA | 494 | 2 | 3vsmA | 638 | 3 | 3spgA | 332 | 2dis |
| 3l76A | 585 | 2 | 3w1bA | 589 | 3 | 3u0kA | 396 | 2dis |
| 3ld1A | 359 | 2 | 3zh9B | 339 | 3 | 3vlaA | 428 | 2dis |
| 3lsgA | 101 | 2 | 4alzA | 212 | 3 | 3vstA | 638 | 2dis |
| 3me4A | 169 | 2 | 4ax8A | 456 | 3 | 4aqfB | 474 | 2dis |
| 3ml4C | 203 | 2 | 4b3iA | 736 | 3 | 4b21A | 207 | 2dis |
| 3mx2B | 517 | 2 | 4bd9B | 165 | 3 | 4dt4A | 160 | 2dis |
| 3mzfA | 362 | 2 | 4c0aB | 355 | 3 | 4dtfA | 359 | 2dis |
| 3njaB | 107 | 2 | 4c0sA | 449 | 3 | 4ewtA | 389 | 2dis |
| 3nqiA | 240 | 2 | 4dimA | 382 | 3 | 4f23A | 509 | 2dis |
| 3nt8A | 401 | 2 | 4indA | 383 | 3 | 4fzbC | 207 | 2dis |
| 3og5A | 155 | 2 | 4jdzB | 447 | 3 | 4g1pA | 479 | 2dis |
| 3oh0A | 593 | 2 | 4kc3B | 278 | 3 | 4gfqA | 186 | 2dis |
| 3pcsB | 355 | 2 | 4kikB | 648 | 3 | 4hvzA | 214 | 2dis |
| 3po3S | 164 | 2 | 4lmfA | 276 | 3 | 4il6B | 505 | 2dis |
| 3pxpA | 288 | 2 | 4lziA | 278 | 3 | 4jxkA | 327 | 2dis |
| 3qavA | 219 | 2 | 4m9pA | 292 | 3 | 4m8mB | 574 | 2dis |
| 3qf4B | 588 | 2 | 4pt5A | 319 | 3 | 4mzyA | 495 | 2dis |
| 3qjjA | 243 | 2 | 4uwhA | 553 | 3 | 4onyA | 583 | 2dis |
| 3qtdA | 435 | 2 | 1c1zA | 325 | 5 | 4pyhA | 293 | 2dis |
| 3r6bA | 115 | 2 | 1d2pA | 373 | 4 | 4rg1A | 285 | 2dis |
| 3rh7A | 289 | 2 | 1k7tA | 170 | 4 | | | |

**Table S2.** Detailed results of domain structure assembly using experimentally solved domains on 356 test proteins.

| PDB ID | TM-score | | | PDB ID | TM-score | | | PDB ID | TM-score | | |
|---|---|---|---|---|---|---|---|---|---|---|---|
| | SADA | DEMO | AIDA | | SADA | DEMO | AIDA | | SADA | DEMO | AIDA |
| 1cjyA | 0.81 | 0.81 | 0.81 | 3rwxA | 0.60 | 0.55 | 0.58 | 1kfqA | 0.97 | 0.97 | 0.68 |
| 1efdN | 0.97 | 1.00 | 0.71 | 3sb4A | 0.61 | 0.57 | 0.56 | 1ldjA | 0.62 | 0.55 | 0.28 |
| 1fjrA | 0.86 | 0.75 | 0.68 | 3swjA | 0.71 | 0.71 | 0.73 | 1nyqB | 0.87 | 0.76 | 0.58 |
| 1g87B | 0.76 | 0.78 | 0.79 | 3t58B | 0.58 | 0.55 | 0.53 | 1ug9A | 0.65 | 0.65 | 0.58 |
| 1hx6B | 0.98 | 0.93 | 0.65 | 3t7jA | 0.97 | 0.98 | 0.52 | 1z1wA | 0.93 | 0.95 | 0.50 |
| 1iwaA | 0.99 | 1.00 | 0.82 | 3u07C | 0.88 | 0.80 | 0.72 | 2au3A | 0.81 | 0.66 | 0.41 |
| 1m5qH | 0.58 | 0.57 | 0.56 | 3u0oB | 0.99 | 0.99 | 0.57 | 2ii2A | 0.87 | 0.78 | 0.66 |
| 1mkfA | 0.99 | 0.95 | 0.61 | 3u9gA | 0.77 | 0.73 | 0.71 | 2olsA | 0.49 | 0.50 | 0.44 |
| 1mkmB | 0.70 | 0.70 | 0.70 | 3ub1D | 0.62 | 0.58 | 0.56 | 2ra1A | 0.49 | 0.33 | 0.44 |
| 1nh2D | 0.58 | 0.54 | 0.53 | 3uitD | 0.57 | 0.52 | 0.55 | 2v5dA | 0.69 | 0.82 | 0.59 |
| 1pprM | 0.57 | 0.99 | 0.93 | 3uo3A | 0.99 | 0.90 | 0.83 | 2xt6A | 0.54 | 0.41 | 0.38 |
| 1prrA | 0.60 | 0.53 | 0.55 | 3v7oB | 0.65 | 0.65 | 0.62 | 2zpaB | 0.90 | 0.44 | 0.41 |
| 1q19A | 0.99 | 0.96 | 0.95 | 3vr8B | 1.00 | 0.66 | 0.61 | 3apoA | 0.45 | 0.41 | 0.29 |
| 1qwrA | 1.00 | 0.98 | 0.75 | 3wkuA | 0.76 | 0.78 | 0.77 | 3b43A | 0.36 | 0.44 | 0.32 |
| 1r71B | 0.88 | 0.73 | 0.55 | 3zvmA | 0.59 | 0.55 | 0.52 | 3gf5B | 0.41 | 0.46 | 0.31 |
| 1rh1A | 0.88 | 0.55 | 0.58 | 4acoA | 0.91 | 0.80 | 0.76 | 3hjlA | 0.36 | 0.39 | 0.38 |
| 1rktA | 0.97 | 0.87 | 0.79 | 4ap5A | 0.99 | 0.74 | 0.62 | 3kq4B | 0.36 | 0.35 | 0.41 |
| 1s6lA | 0.76 | 0.73 | 0.80 | 4axdA | 0.99 | 0.88 | 0.77 | 3kwlA | 0.58 | 0.53 | 0.38 |
| 1sp3A | 1.00 | 0.59 | 0.59 | 4bfiB | 0.83 | 0.96 | 0.59 | 3ob8A | 0.97 | 0.68 | 0.47 |
| 1vz6A | 0.95 | 0.82 | 0.75 | 4bt9B | 0.61 | 0.63 | 0.61 | 3opfB | 0.68 | 0.63 | 0.42 |
| 1w3aA | 0.62 | 0.57 | 0.54 | 4cczA | 0.96 | 0.61 | 0.93 | 3p53A | 0.37 | 0.35 | 0.37 |
| 1wv3A | 0.93 | 0.84 | 0.85 | 4d0nB | 0.91 | 0.93 | 0.69 | 3pvlA | 0.68 | 0.75 | 0.51 |
| 1x7pA | 0.99 | 0.78 | 0.63 | 4d1iG | 0.96 | 0.79 | 0.72 | 3r05A | 0.32 | 0.31 | 0.30 |
| 1x9yA | 0.63 | 0.98 | 0.57 | 4dj3A | 0.98 | 0.94 | 0.61 | 3ubhA | 0.68 | 0.51 | 0.33 |
| 1y11A | 0.60 | 0.55 | 0.57 | 4dqaA | 0.64 | 0.63 | 0.62 | 3w2wA | 0.85 | 0.51 | 0.53 |
| 1yiqA | 0.88 | 0.99 | 0.89 | 4eo3A | 0.58 | 0.57 | 0.59 | 3zniA | 0.41 | 0.35 | 0.40 |
| 1zbuB | 0.79 | 0.79 | 0.76 | 4eogA | 0.98 | 0.68 | 0.66 | 4aimA | 0.91 | 0.51 | 0.49 |
| 1ze1A | 0.99 | 0.96 | 0.79 | 4etxA | 0.55 | 0.55 | 0.60 | 4ak1A | 0.29 | 0.28 | 0.29 |
| 2ablA | 0.70 | 0.63 | 0.64 | 4fguA | 0.92 | 0.69 | 0.69 | 4aq1A | 0.45 | 0.46 | 0.35 |
| 2ahvA | 0.82 | 0.75 | 0.80 | 4fkcA | 0.94 | 0.94 | 0.73 | 4fe9A | 0.47 | 0.44 | 0.38 |
| 2bkpA | 0.83 | 0.59 | 0.79 | 4fxkC | 0.97 | 0.96 | 0.54 | 4h2aA | 0.61 | 0.53 | 0.43 |
| 2c1yA | 0.56 | 0.70 | 0.58 | 4gbyA | 1.00 | 0.99 | 1.00 | 4i5sB | 0.55 | 0.40 | 0.41 |
| 2cxcA | 0.96 | 0.90 | 0.51 | 4ggmX | 0.61 | 0.57 | 0.74 | 4iggB | 0.31 | 0.33 | 0.31 |
| 2d1cA | 0.84 | 0.79 | 0.78 | 4gslA | 0.55 | 0.54 | 0.53 | 4j9vA | 0.87 | 0.61 | 0.46 |
| 2d7iA | 0.74 | 0.77 | 0.84 | 4gyjA | 1.00 | 1.00 | 0.67 | 4k3bA | 0.56 | 0.54 | 0.49 |
| 2e9hA | 0.97 | 0.96 | 0.81 | 4h3tA | 0.98 | 0.89 | 0.79 | 4kwuA | 0.78 | 0.82 | 0.46 |
| 2e9xB | 0.99 | 0.86 | 0.88 | 4hmoA | 0.99 | 0.94 | 0.55 | 4m00A | 0.52 | 0.51 | 0.52 |
| 2evrA | 0.99 | 0.69 | 0.68 | 4ie6A | 0.70 | 0.70 | 0.70 | 1bp1A | 0.99 | 0.98 | 0.73 |
| 2ew9A | 0.54 | 0.57 | 0.57 | 4l5gA | 0.97 | 0.67 | 0.64 | 1cjsA | 0.84 | 0.97 | 0.64 |
| 2fd5A | 0.99 | 0.97 | 0.75 | 4lpqA | 0.91 | 0.70 | 0.66 | 1ck1A | 0.99 | 0.95 | 0.64 |

| | | | | | | | | | | | |
|---|---|---|---|---|---|---|---|---|---|---|---|
| 2gh8A | 0.72 | 0.66 | 0.78 | 4m8rA | 0.77 | 0.79 | 0.76 | 1ecrA | 0.88 | 0.87 | 0.85 |
| 2gt1A | 0.97 | 0.96 | 0.64 | 4n06B | 0.99 | 0.90 | 0.81 | 1f5qD | 0.99 | 0.95 | 0.77 |
| 2gzaC | 0.98 | 0.92 | 0.67 | 4nj5A | 0.69 | 0.69 | 0.61 | 1fa9A | 1.00 | 0.94 | 0.68 |
| 2hjqA | 0.55 | 0.58 | 0.53 | 4opaB | 0.71 | 0.68 | 0.67 | 1gu7A | 1.00 | 0.94 | 0.61 |
| 2hwjA | 0.83 | 0.68 | 0.70 | 4qkuB | 1.00 | 0.73 | 0.72 | 1itwA | 0.99 | 0.61 | 0.62 |
| 2ijd1 | 0.77 | 0.75 | 0.76 | 4up9A | 0.96 | 0.98 | 0.97 | 1jkiA | 1.00 | 0.91 | 0.82 |
| 2iu7A | 0.60 | 0.58 | 0.58 | 4w7sA | 0.64 | 0.63 | 0.64 | 1n80A | 1.00 | 0.84 | 0.78 |
| 2iw2A | 0.98 | 0.96 | 0.67 | 1bf2A | 0.99 | 0.99 | 0.71 | 1nzjA | 0.99 | 0.98 | 0.82 |
| 2jz4A | 0.59 | 0.51 | 0.51 | 1bhgA | 1.00 | 0.97 | 0.58 | 1qhdA | 0.90 | 0.88 | 0.60 |
| 2kdyA | 0.59 | 0.84 | 0.62 | 1f7uA | 0.97 | 0.76 | 0.64 | 1qz9A | 0.99 | 0.99 | 0.77 |
| 2kn4A | 0.65 | 0.61 | 0.60 | 1fx7A | 0.87 | 0.78 | 0.63 | 1sb7B | 0.87 | 0.97 | 0.64 |
| 2mbgA | 0.78 | 0.82 | 0.76 | 1griA | 0.68 | 0.52 | 0.52 | 1vk1A | 0.85 | 0.62 | 0.62 |
| 2nsfA | 0.94 | 0.72 | 0.68 | 1h88C | 0.71 | 0.62 | 0.46 | 1vrmA | 0.99 | 0.90 | 0.71 |
| 2nykA | 0.95 | 0.81 | 0.74 | 1m8pB | 0.68 | 0.70 | 0.38 | 1xvuA | 0.98 | 0.91 | 0.82 |
| 2o6yA | 0.99 | 0.98 | 0.67 | 1ni5A | 0.75 | 0.67 | 0.42 | 1yy3A | 0.70 | 0.70 | 0.70 |
| 2owbA | 0.98 | 1.00 | 0.70 | 1q25A | 0.49 | 0.44 | 0.41 | 1z87A | 0.67 | 0.54 | 0.52 |
| 2qfiA | 0.87 | 0.72 | 0.72 | 1uzjA | 0.78 | 0.52 | 0.49 | 2a1sC | 0.86 | 0.82 | 0.81 |
| 2qp2A | 0.74 | 0.73 | 0.75 | 1zpuA | 0.99 | 0.99 | 0.50 | 2a3lA | 0.97 | 0.99 | 0.86 |
| 2qygA | 0.97 | 0.99 | 0.68 | 1zy9A | 0.92 | 0.96 | 0.45 | 2bt1A | 0.99 | 0.93 | 0.66 |
| 2r5wB | 0.80 | 0.60 | 0.58 | 2b5uA | 0.48 | 0.45 | 0.51 | 2bydA | 0.98 | 0.58 | 0.63 |
| 2uu7A | 0.99 | 0.96 | 0.75 | 2ewfA | 0.49 | 0.55 | 0.47 | 2c43A | 0.93 | 0.57 | 0.63 |
| 2w4bA | 0.83 | 0.79 | 0.77 | 2piaA | 0.86 | 0.75 | 0.74 | 2dfyC | 0.66 | 0.62 | 0.48 |
| 2x7iA | 0.98 | 0.96 | 0.58 | 2r7dA | 0.98 | 0.92 | 0.75 | 2dlaA | 1.00 | 0.97 | 0.93 |
| 2x8kC | 0.98 | 0.71 | 0.83 | 2uwnA | 0.77 | 0.48 | 0.37 | 2g3pA | 0.63 | 0.87 | 0.57 |
| 2yilA | 0.63 | 0.63 | 0.64 | 2v0nA | 0.45 | 0.41 | 0.41 | 2gg6A | 0.99 | 0.99 | 0.72 |
| 2yrqA | 0.52 | 0.51 | 0.53 | 2vgmA | 0.76 | 0.66 | 0.40 | 2gsyE | 0.92 | 0.70 | 0.68 |
| 2zxcA | 0.82 | 0.81 | 0.80 | 2wqrB | 0.49 | 0.67 | 0.37 | 2gzoA | 0.93 | 0.57 | 0.53 |
| 3a1iA | 0.93 | 0.90 | 0.89 | 2y25B | 0.57 | 0.42 | 0.40 | 2j2cA | 0.96 | 0.72 | 0.66 |
| 3a45A | 0.99 | 0.99 | 0.54 | 2yk0A | 0.72 | 0.74 | 0.43 | 2kfwA | 0.78 | 0.77 | 0.73 |
| 3a56A | 0.66 | 0.76 | 0.66 | 2zzqA | 0.46 | 0.72 | 0.72 | 2l9yA | 0.69 | 0.74 | 0.72 |
| 3ajvA | 0.98 | 0.76 | 0.55 | 3bt1U | 0.61 | 0.40 | 0.42 | 2ntyB | 0.82 | 0.71 | 0.72 |
| 3aqkA | 0.96 | 0.96 | 0.73 | 3c1yA | 0.63 | 0.49 | 0.49 | 2r3vA | 1.00 | 0.96 | 0.71 |
| 3arbA | 0.94 | 0.95 | 0.85 | 3cw2C | 0.70 | 0.65 | 0.66 | 2r58A | 0.65 | 0.99 | 0.53 |
| 3aujG | 0.99 | 0.70 | 0.71 | 3f83A | 0.46 | 0.39 | 0.41 | 2w4mA | 0.89 | 0.78 | 0.77 |
| 3b2zF | 0.76 | 0.76 | 0.81 | 3fc3A | 0.54 | 0.47 | 0.45 | 2x0cA | 0.66 | 0.63 | 0.58 |
| 3b7wA | 0.83 | 0.83 | 0.83 | 3gbgA | 0.64 | 0.60 | 0.63 | 2y51A | 0.99 | 0.98 | 0.83 |
| 3bt3A | 0.95 | 0.92 | 0.62 | 3h5cB | 0.97 | 0.86 | 0.60 | 2yb0E | 0.97 | 0.89 | 0.83 |
| 3c4tA | 0.74 | 0.74 | 0.74 | 3ibjA | 0.58 | 0.50 | 0.49 | 2z86C | 0.69 | 0.76 | 0.57 |
| 3craA | 0.61 | 0.58 | 0.54 | 3ippB | 0.74 | 0.44 | 0.55 | 3afoA | 0.67 | 0.78 | 0.63 |
| 3d30A | 0.96 | 0.91 | 0.57 | 3jymB | 0.43 | 0.49 | 0.44 | 3bu2A | 0.91 | 0.68 | 0.70 |
| 3eo5A | 0.64 | 0.60 | 0.53 | 3kbgA | 0.62 | 0.76 | 0.46 | 3cvzA | 0.87 | 0.61 | 0.61 |
| 3errA | 0.79 | 0.64 | 0.66 | 3mc8A | 0.83 | 0.87 | 0.51 | 3dupA | 0.98 | 0.83 | 0.68 |
| 3g79A | 0.90 | 0.84 | 0.60 | 3npfA | 0.93 | 0.96 | 0.54 | 3eswA | 0.89 | 0.89 | 0.91 |
| 3h2tA | 0.81 | 0.82 | 0.86 | 3orjA | 0.66 | 0.62 | 0.57 | 3eukH | 0.92 | 0.91 | 0.88 |

| | | | | | | | | | | | |
|---|---|---|---|---|---|---|---|---|---|---|---|
| 3hcsA | 0.80 | 0.70 | 0.69 | 3plaA | 0.65 | 0.63 | 0.63 | 3fi7A | 0.99 | 0.99 | 0.75 |
| 3hyiA | 0.71 | 0.69 | 0.68 | 3qe9Y | 0.94 | 0.93 | 0.67 | 3fvvA | 0.98 | 0.93 | 0.72 |
| 3i2dA | 0.86 | 0.78 | 0.56 | 3qjoA | 0.80 | 0.99 | 0.76 | 3gmsA | 1.00 | 0.99 | 0.93 |
| 3iam2 | 0.92 | 0.64 | 0.60 | 3qphA | 0.48 | 0.57 | 0.58 | 3hzzB | 1.00 | 0.93 | 0.79 |
| 3ifrA | 0.99 | 0.98 | 0.65 | 3qyeA | 0.99 | 0.95 | 0.53 | 3m1uA | 0.98 | 0.60 | 0.73 |
| 3isqA | 1.00 | 0.99 | 0.62 | 3rimA | 0.98 | 0.97 | 0.54 | 3mw8A | 0.75 | 0.94 | 0.55 |
| 3j7aK | 0.96 | 0.99 | 0.71 | 3rrpA | 0.98 | 0.97 | 0.66 | 3mwcA | 1.00 | 0.98 | 0.91 |
| 3k1rA | 0.66 | 0.61 | 0.58 | 3soaA | 0.73 | 0.73 | 0.53 | 3nsjA | 0.82 | 0.84 | 0.78 |
| 3k2iA | 0.78 | 0.73 | 0.70 | 3tixD | 0.63 | 0.61 | 0.39 | 3ntkA | 0.98 | 0.97 | 0.61 |
| 3kh5A | 0.94 | 0.98 | 0.56 | 3tp9A | 0.87 | 0.57 | 0.59 | 3oaaG | 0.98 | 0.98 | 0.56 |
| 3kjpA | 0.91 | 0.71 | 0.53 | 3ua3A | 0.58 | 0.51 | 0.45 | 3ptyA | 0.75 | 0.73 | 0.74 |
| 3kt1A | 0.91 | 0.58 | 0.58 | 3uj0A | 0.94 | 0.56 | 0.50 | 3rfyA | 0.95 | 0.73 | 0.76 |
| 3ktmE | 0.63 | 0.59 | 0.62 | 3vn4A | 0.47 | 0.44 | 0.46 | 3seoB | 0.77 | 0.65 | 0.61 |
| 3kzwA | 1.00 | 0.99 | 0.68 | 3vsmA | 0.80 | 0.98 | 0.56 | 3spgA | 0.98 | 0.86 | 0.66 |
| 3l76A | 0.89 | 0.65 | 0.72 | 3w1bA | 0.72 | 0.72 | 0.59 | 3u0kA | 0.61 | 0.61 | 0.60 |
| 3ld1A | 0.70 | 0.70 | 0.70 | 3zh9B | 0.62 | 0.58 | 0.58 | 3vlaA | 1.00 | 0.98 | 0.86 |
| 3lsgA | 0.86 | 0.86 | 0.61 | 4alzA | 0.69 | 0.38 | 0.46 | 3vstA | 1.00 | 0.98 | 0.83 |
| 3me4A | 0.90 | 0.63 | 0.60 | 4ax8A | 0.64 | 0.69 | 0.52 | 4aqfB | 0.88 | 0.92 | 0.84 |
| 3ml4C | 0.65 | 0.93 | 0.55 | 4b3iA | 0.94 | 0.68 | 0.50 | 4b21A | 0.99 | 0.93 | 0.80 |
| 3mx2B | 0.92 | 0.89 | 0.73 | 4bd9B | 0.61 | 0.51 | 0.39 | 4dt4A | 0.98 | 0.96 | 0.69 |
| 3mzfA | 0.93 | 0.86 | 0.75 | 4c0aB | 0.57 | 0.52 | 0.49 | 4dtfA | 0.95 | 0.85 | 0.81 |
| 3njaB | 0.61 | 0.61 | 0.58 | 4c0sA | 0.63 | 0.64 | 0.62 | 4ewtA | 0.84 | 0.88 | 0.72 |
| 3nqiA | 0.94 | 0.93 | 0.74 | 4dimA | 0.89 | 0.90 | 0.62 | 4f23A | 1.00 | 0.67 | 0.73 |
| 3nt8A | 0.76 | 0.58 | 0.52 | 4indA | 0.46 | 0.42 | 0.41 | 4fzbC | 0.99 | 0.92 | 0.74 |
| 3og5A | 0.97 | 0.92 | 0.78 | 4jdzB | 0.70 | 0.63 | 0.45 | 4g1pA | 0.93 | 0.96 | 0.77 |
| 3oh0A | 0.97 | 0.78 | 0.83 | 4kc3B | 0.76 | 0.67 | 0.37 | 4gfqA | 0.76 | 0.66 | 0.62 |
| 3pcsB | 0.86 | 0.80 | 0.76 | 4kikB | 0.73 | 0.85 | 0.67 | 4hvzA | 0.75 | 0.55 | 0.59 |
| 3po3S | 0.96 | 0.58 | 0.56 | 4lmfA | 0.49 | 0.47 | 0.44 | 4il6B | 0.73 | 0.71 | 0.70 |
| 3pxpA | 0.74 | 0.70 | 0.69 | 4lziA | 0.67 | 0.46 | 0.42 | 4jxkA | 0.98 | 0.97 | 0.76 |
| 3qavA | 0.98 | 0.91 | 0.93 | 4m9pA | 0.51 | 0.36 | 0.38 | 4m8mB | 0.87 | 0.80 | 0.80 |
| 3qf4B | 0.93 | 0.74 | 0.62 | 4pt5A | 0.91 | 0.82 | 0.46 | 4mzyA | 1.00 | 0.95 | 0.79 |
| 3qjjA | 0.97 | 0.88 | 0.57 | 4uwhA | 0.99 | 0.94 | 0.49 | 4onyA | 0.81 | 0.82 | 0.59 |
| 3qtdA | 1.00 | 0.99 | 0.62 | 1c1zA | 0.46 | 0.41 | 0.30 | 4pyhA | 0.77 | 0.93 | 0.78 |
| 3r6bA | 0.54 | 0.65 | 0.51 | 1d2pA | 0.36 | 0.29 | 0.30 | 4rg1A | 0.77 | 0.85 | 0.65 |
| 3rh7A | 0.71 | 0.61 | 0.80 | 1k7tA | 0.67 | 0.38 | 0.39 | | | | |

**Table S3.** Summary of domain structure assembly by using experimentally solved domains on 356 test proteins.

| Domain | Method | TM-score | #TM-score⩾0.5 | *P*-value |
|---|---|---|---|---|
| 2dom (*N* = 166) | SADA | 0.83 | 166 | - |
| | DEMO | 0.78 | 166 | 7.92E-13 |
| | AIDA | 0.68 | 166 | 8.86E-22 |
| 3dom (*N* = 69) | SADA | 0.72 | 57 | - |
| | DEMO | 0.67 | 51 | 2.07E-04 |
| | AIDA | 0.52 | 33 | 1.24E-11 |
| m4dom (*N* = 40) | SADA | 0.60 | 24 | - |
| | DEMO | 0.53 | 20 | 7.20E-04 |
| | AIDA | 0.42 | 9 | 3.41E-07 |
| 2dis (*N* = 81) | SADA | 0.89 | 81 | - |
| | DEMO | 0.84 | 81 | 3.80E-05 |
| | AIDA | 0.71 | 80 | 1.22E-14 |
| All (*N* = 356) | SADA | 0.80 | 328 | - |
| | DEMO | 0.74 | 318 | 1.28E-21 |
| | AIDA | 0.63 | 288 | 8.91E-51 |

**Table S4.** Coverage scores of each test protein in MPDB and DEMO-lib under the sequence identity cutoff of 30%.

| PDB ID | Coverage score | | PDB ID | Coverage score | | PDB ID | Coverage score | |
|---|---|---|---|---|---|---|---|---|
| | MPDB | DEMO-lib | | MPDB | DEMO-lib | | MPDB | DEMO-lib |
| 1cjyA | 0.38 | 0.36 | 3rwxA | 0.45 | 0.44 | 1kfqA | 0.73 | 0.73 |
| 1efdN | 0.83 | 0.80 | 3sb4A | 0.53 | 0.51 | 1ldjA | 0.44 | 0.43 |
| 1fjrA | 0.45 | 0.43 | 3swjA | 0.51 | 0.48 | 1nyqB | 0.59 | 0.56 |
| 1g87B | 0.61 | 0.55 | 3t58B | 0.42 | 0.38 | 1ug9A | 0.53 | 0.51 |
| 1hx6B | 0.67 | 0.60 | 3t7jA | 0.65 | 0.63 | 1z1wA | 0.89 | 0.85 |
| 1iwaA | 0.84 | 0.79 | 3u07C | 0.45 | 0.40 | 2au3A | 0.65 | 0.54 |
| 1m5qH | 0.48 | 0.47 | 3u0oB | 0.73 | 0.71 | 2ii2A | 0.84 | 0.82 |
| 1mkfA | 0.43 | 0.42 | 3u9gA | 0.44 | 0.43 | 2olsA | 0.41 | 0.38 |
| 1mkmB | 0.65 | 0.57 | 3ub1D | 0.41 | 0.40 | 2ra1A | 0.41 | 0.39 |
| 1nh2D | 0.49 | 0.49 | 3uitD | 0.48 | 0.43 | 2v5dA | 0.62 | 0.54 |
| 1pprM | 0.45 | 0.43 | 3uo3A | 0.61 | 0.55 | 2xt6A | 0.50 | 0.48 |
| 1prrA | 0.47 | 0.45 | 3v7oB | 0.46 | 0.46 | 2zpaB | 0.37 | 0.36 |
| 1q19A | 0.56 | 0.46 | 3vr8B | 0.52 | 0.43 | 3apoA | 0.37 | 0.34 |
| 1qwrA | 0.83 | 0.67 | 3wkuA | 0.42 | 0.40 | 3b43A | 0.46 | 0.44 |
| 1r71B | 0.62 | 0.52 | 3zvmA | 0.42 | 0.41 | 3gf5B | 0.39 | 0.38 |
| 1rh1A | 0.40 | 0.37 | 4acoA | 0.41 | 0.37 | 3hjlA | 0.43 | 0.41 |
| 1rktA | 0.71 | 0.70 | 4ap5A | 0.61 | 0.54 | 3kq4B | 0.41 | 0.40 |
| 1s6lA | 0.44 | 0.43 | 4axdA | 0.40 | 0.36 | 3kwlA | 0.41 | 0.39 |
| 1sp3A | 0.39 | 0.38 | 4bfiB | 0.79 | 0.77 | 3ob8A | 0.52 | 0.51 |
| 1vz6A | 0.43 | 0.42 | 4bt9B | 0.52 | 0.50 | 3opfB | 0.37 | 0.39 |
| 1w3aA | 0.48 | 0.47 | 4cczA | 0.50 | 0.47 | 3p53A | 0.40 | 0.39 |
| 1wv3A | 0.49 | 0.48 | 4d0nB | 0.76 | 0.70 | 3pvlA | 0.60 | 0.52 |
| 1x7pA | 0.64 | 0.54 | 4d1iG | 0.65 | 0.64 | 3r05A | 0.33 | 0.31 |
| 1x9yA | 0.47 | 0.46 | 4dj3A | 0.56 | 0.46 | 3ubhA | 0.48 | 0.48 |
| 1y11A | 0.49 | 0.45 | 4dqaA | 0.45 | 0.43 | 3w2wA | 0.57 | 0.39 |
| 1yiqA | 0.49 | 0.47 | 4eo3A | 0.48 | 0.46 | 3zniA | 0.40 | 0.40 |
| 1zbuB | 0.55 | 0.55 | 4eogA | 0.60 | 0.42 | 4aimA | 0.36 | 0.34 |
| 1ze1A | 0.62 | 0.55 | 4etxA | 0.43 | 0.42 | 4ak1A | 0.38 | 0.37 |
| 2ablA | 0.55 | 0.52 | 4fguA | 0.43 | 0.42 | 4aq1A | 0.34 | 0.33 |
| 2ahvA | 0.74 | 0.67 | 4fkcA | 0.88 | 0.87 | 4fe9A | 0.39 | 0.38 |
| 2bkpA | 0.50 | 0.49 | 4fxkC | 0.55 | 0.47 | 4h2aA | 0.40 | 0.36 |
| 2c1yA | 0.56 | 0.50 | 4gbyA | 0.78 | 0.73 | 4i5sB | 0.46 | 0.43 |
| 2cxcA | 0.63 | 0.56 | 4ggmX | 0.44 | 0.44 | 4iggB | 0.55 | 0.36 |
| 2d1cA | 0.65 | 0.65 | 4gslA | 0.40 | 0.38 | 4j9vA | 0.58 | 0.54 |
| 2d7iA | 0.45 | 0.42 | 4gyjA | 0.80 | 0.78 | 4k3bA | 0.38 | 0.33 |
| 2e9hA | 0.47 | 0.47 | 4h3tA | 0.54 | 0.45 | 4kwuA | 0.67 | 0.66 |
| 2e9xB | 0.63 | 0.55 | 4hmoA | 0.74 | 0.74 | 4m00A | 0.38 | 0.37 |
| 2evrA | 0.58 | 0.54 | 4ie6A | 0.42 | 0.40 | 1bp1A | 0.44 | 0.41 |
| 2ew9A | 0.55 | 0.54 | 4l5gA | 0.50 | 0.48 | 1cjsA | 0.65 | 0.63 |
| 2fd5A | 0.78 | 0.78 | 4lpqA | 0.52 | 0.51 | 1ck1A | 0.77 | 0.71 |

| | | | | | | | | |
|---|---|---|---|---|---|---|---|---|
| 2gh8A | 0.52 | 0.46 | 4m8rA | 0.54 | 0.53 | 1ecrA | 0.38 | 0.38 |
| 2gt1A | 0.68 | 0.66 | 4n06B | 0.75 | 0.68 | 1f5qD | 0.80 | 0.77 |
| 2gzaC | 0.48 | 0.47 | 4nj5A | 0.37 | 0.36 | 1fa9A | 0.50 | 0.46 |
| 2hjqA | 0.50 | 0.49 | 4opaB | 0.45 | 0.44 | 1gu7A | 0.77 | 0.76 |
| 2hwjA | 0.47 | 0.46 | 4qkuB | 0.57 | 0.45 | 1itwA | 0.48 | 0.46 |
| 2ijd1 | 0.67 | 0.62 | 4up9A | 0.73 | 0.72 | 1jkiA | 0.58 | 0.49 |
| 2iu7A | 0.50 | 0.48 | 4w7sA | 0.53 | 0.50 | 1n80A | 0.37 | 0.36 |
| 2iw2A | 0.80 | 0.75 | 1bf2A | 0.75 | 0.73 | 1nzjA | 0.74 | 0.74 |
| 2jz4A | 0.48 | 0.46 | 1bhgA | 0.83 | 0.80 | 1qhdA | 0.43 | 0.39 |
| 2kdyA | 0.52 | 0.46 | 1f7uA | 0.70 | 0.64 | 1qz9A | 0.85 | 0.82 |
| 2kn4A | 0.48 | 0.48 | 1fx7A | 0.65 | 0.50 | 1sb7B | 0.45 | 0.44 |
| 2mbgA | 0.63 | 0.62 | 1griA | 0.46 | 0.43 | 1vk1A | 0.48 | 0.41 |
| 2nsfA | 0.47 | 0.45 | 1h88C | 0.49 | 0.48 | 1vrmA | 0.46 | 0.41 |
| 2nykA | 0.67 | 0.66 | 1m8pB | 0.59 | 0.47 | 1xvuA | 0.86 | 0.70 |
| 2o6yA | 0.72 | 0.52 | 1ni5A | 0.46 | 0.43 | 1yy3A | 0.41 | 0.40 |
| 2owbA | 0.93 | 0.90 | 1q25A | 0.40 | 0.38 | 1z87A | 0.38 | 0.37 |
| 2qfiA | 0.44 | 0.42 | 1uzjA | 0.42 | 0.42 | 2a1sC | 0.41 | 0.40 |
| 2qp2A | 0.50 | 0.46 | 1zpuA | 0.87 | 0.74 | 2a3lA | 0.44 | 0.44 |
| 2qygA | 0.89 | 0.87 | 1zy9A | 0.78 | 0.74 | 2bt1A | 0.50 | 0.43 |
| 2r5wB | 0.46 | 0.43 | 2b5uA | 0.37 | 0.36 | 2bydA | 0.51 | 0.45 |
| 2uu7A | 0.77 | 0.70 | 2ewfA | 0.39 | 0.37 | 2c43A | 0.51 | 0.45 |
| 2w4bA | 0.36 | 0.36 | 2piaA | 0.63 | 0.63 | 2dfyC | 0.43 | 0.42 |
| 2x7iA | 0.77 | 0.75 | 2r7dA | 0.70 | 0.46 | 2dlaA | 0.51 | 0.49 |
| 2x8kC | 0.51 | 0.49 | 2uwnA | 0.48 | 0.47 | 2g3pA | 0.40 | 0.40 |
| 2yilA | 0.54 | 0.48 | 2v0nA | 0.41 | 0.39 | 2gg6A | 0.87 | 0.82 |
| 2yrqA | 0.44 | 0.43 | 2vgmA | 0.56 | 0.51 | 2gsyE | 0.45 | 0.42 |
| 2zxcA | 0.33 | 0.32 | 2wqrB | 0.58 | 0.52 | 2gzoA | 0.42 | 0.40 |
| 3a1iA | 0.55 | 0.48 | 2y25B | 0.44 | 0.42 | 2j2cA | 0.43 | 0.40 |
| 3a45A | 0.80 | 0.77 | 2yk0A | 0.42 | 0.39 | 2kfwA | 0.43 | 0.43 |
| 3a56A | 0.43 | 0.41 | 2zzqA | 0.44 | 0.42 | 2l9yA | 0.41 | 0.41 |
| 3ajvA | 0.68 | 0.57 | 3bt1U | 0.42 | 0.42 | 2ntyB | 0.41 | 0.41 |
| 3aqkA | 0.74 | 0.66 | 3c1yA | 0.43 | 0.42 | 2r3vA | 0.70 | 0.68 |
| 3arbA | 0.84 | 0.82 | 3cw2C | 0.55 | 0.51 | 2r58A | 0.42 | 0.41 |
| 3aujG | 0.49 | 0.47 | 3f83A | 0.42 | 0.41 | 2w4mA | 0.74 | 0.72 |
| 3b2zF | 0.63 | 0.55 | 3fc3A | 0.44 | 0.43 | 2x0cA | 0.45 | 0.43 |
| 3b7wA | 0.93 | 0.85 | 3gbgA | 0.43 | 0.42 | 2y51A | 0.85 | 0.84 |
| 3bt3A | 0.74 | 0.72 | 3h5cB | 0.80 | 0.78 | 2yb0E | 0.46 | 0.45 |
| 3c4tA | 0.58 | 0.50 | 3ibjA | 0.44 | 0.42 | 2z86C | 0.37 | 0.36 |
| 3craA | 0.45 | 0.43 | 3ippB | 0.60 | 0.56 | 3afoA | 0.69 | 0.61 |
| 3d30A | 0.51 | 0.50 | 3jymB | 0.43 | 0.42 | 3bu2A | 0.43 | 0.42 |
| 3eo5A | 0.43 | 0.41 | 3kbgA | 0.42 | 0.42 | 3cvzA | 0.40 | 0.40 |
| 3errA | 0.69 | 0.61 | 3mc8A | 0.52 | 0.50 | 3dupA | 0.49 | 0.47 |
| 3g79A | 0.79 | 0.71 | 3npfA | 0.56 | 0.49 | 3eswA | 0.50 | 0.42 |
| 3h2tA | 0.39 | 0.38 | 3orjA | 0.47 | 0.41 | 3eukH | 0.59 | 0.57 |

| | | | | | | | | |
|---|---|---|---|---|---|---|---|---|
| 3hcsA | 0.47 | 0.47 | 3plaA | 0.45 | 0.40 | 3fi7A | 0.61 | 0.55 |
| 3hyiA | 0.50 | 0.49 | 3qe9Y | 0.72 | 0.65 | 3fvvA | 0.76 | 0.70 |
| 3i2dA | 0.50 | 0.46 | 3qjoA | 0.55 | 0.42 | 3gmsA | 0.88 | 0.88 |
| 3iam2 | 0.55 | 0.46 | 3qphA | 0.42 | 0.42 | 3hzzB | 0.72 | 0.71 |
| 3ifrA | 0.91 | 0.90 | 3qyeA | 0.75 | 0.70 | 3m1uA | 0.48 | 0.40 |
| 3isqA | 0.71 | 0.71 | 3rimA | 0.82 | 0.60 | 3mw8A | 0.62 | 0.59 |
| 3j7aK | 0.84 | 0.63 | 3rrpA | 0.80 | 0.77 | 3mwcA | 0.86 | 0.86 |
| 3k1rA | 0.46 | 0.45 | 3soaA | 0.63 | 0.59 | 3nsjA | 0.50 | 0.47 |
| 3k2iA | 0.59 | 0.58 | 3tixD | 0.38 | 0.37 | 3ntkA | 0.82 | 0.63 |
| 3kh5A | 0.83 | 0.57 | 3tp9A | 0.46 | 0.43 | 3oaaG | 0.64 | 0.53 |
| 3kjpA | 0.52 | 0.43 | 3ua3A | 0.42 | 0.42 | 3ptyA | 0.45 | 0.44 |
| 3kt1A | 0.43 | 0.42 | 3uj0A | 0.70 | 0.48 | 3rfyA | 0.38 | 0.38 |
| 3ktmE | 0.45 | 0.44 | 3vn4A | 0.39 | 0.38 | 3seoB | 0.45 | 0.44 |
| 3kzwA | 0.86 | 0.76 | 3vsmA | 0.76 | 0.71 | 3spgA | 0.75 | 0.46 |
| 3l76A | 0.52 | 0.48 | 3w1bA | 0.64 | 0.52 | 3u0kA | 0.55 | 0.41 |
| 3ld1A | 0.42 | 0.41 | 3zh9B | 0.52 | 0.49 | 3vlaA | 0.72 | 0.68 |
| 3lsgA | 0.77 | 0.74 | 4alzA | 0.46 | 0.46 | 3vstA | 0.49 | 0.47 |
| 3me4A | 0.56 | 0.51 | 4ax8A | 0.45 | 0.43 | 4aqfB | 0.39 | 0.38 |
| 3ml4C | 0.49 | 0.48 | 4b3iA | 0.60 | 0.53 | 4b21A | 0.84 | 0.79 |
| 3mx2B | 0.44 | 0.41 | 4bd9B | 0.43 | 0.43 | 4dt4A | 0.58 | 0.55 |
| 3mzfA | 0.76 | 0.65 | 4c0aB | 0.50 | 0.45 | 4dtfA | 0.40 | 0.39 |
| 3njaB | 0.48 | 0.46 | 4c0sA | 0.72 | 0.51 | 4ewtA | 0.79 | 0.78 |
| 3nqiA | 0.56 | 0.55 | 4dimA | 0.80 | 0.78 | 4f23A | 0.60 | 0.50 |
| 3nt8A | 0.40 | 0.39 | 4indA | 0.41 | 0.39 | 4fzbC | 0.61 | 0.45 |
| 3og5A | 0.59 | 0.53 | 4jdzB | 0.60 | 0.58 | 4g1pA | 0.74 | 0.72 |
| 3oh0A | 0.65 | 0.61 | 4kc3B | 0.72 | 0.71 | 4gfqA | 0.51 | 0.49 |
| 3pcsB | 0.47 | 0.41 | 4kikB | 0.63 | 0.42 | 4hvzA | 0.46 | 0.43 |
| 3po3S | 0.45 | 0.44 | 4lmfA | 0.45 | 0.44 | 4il6B | 0.54 | 0.45 |
| 3pxpA | 0.42 | 0.41 | 4lziA | 0.41 | 0.41 | 4jxkA | 0.84 | 0.84 |
| 3qavA | 0.81 | 0.81 | 4m9pA | 0.44 | 0.44 | 4m8mB | 0.36 | 0.34 |
| 3qf4B | 0.80 | 0.71 | 4pt5A | 0.65 | 0.57 | 4mzyA | 0.56 | 0.58 |
| 3qjjA | 0.70 | 0.64 | 4uwhA | 0.85 | 0.60 | 4onyA | 0.73 | 0.71 |
| 3qtdA | 0.42 | 0.42 | 1c1zA | 0.44 | 0.41 | 4pyhA | 0.58 | 0.54 |
| 3r6bA | 0.48 | 0.47 | 1d2pA | 0.39 | 0.38 | 4rg1A | 0.54 | 0.50 |
| 3rh7A | 0.45 | 0.44 | 1k7tA | 0.46 | 0.46 | | | |

**Table S5.** Under the sequence identity cutoff of 30%, average coverage scores of 2dom proteins, 3dom proteins, m4dom proteins, 2dis proteins and all test protein in MPDB and DEMO-lib. The values in the last column are the results of the Wilcoxon signed-rank test based on comparison to the coverage scores of MPDB.

| Database | 2dom (166) | 3dom (69) | m4dom (40) | 2dis (81) | Total (356) | *P*-value |
|---|---|---|---|---|---|---|
| MPDB | 0.57 | 0.56 | 0.48 | 0.56 | 0.56 | NA |
| DEMO-lib | 0.53 | 0.50 | 0.45 | 0.53 | 0.52 | 5.54E-53 |

**Table S6.** Coverage scores of each test protein in MPDB and DEMO-lib under the sequence identity cutoff of 50%.

| PDB ID | Coverage score | | PDB ID | Coverage score | | PDB ID | Coverage score | |
|---|---|---|---|---|---|---|---|---|
| | MPDB | DEMO-lib | | MPDB | DEMO-lib | | MPDB | DEMO-lib |
| 1cjyA | 0.47 | 0.41 | 3rwxA | 0.45 | 0.44 | 1kfqA | 0.80 | 0.75 |
| 1efdN | 0.85 | 0.83 | 3sb4A | 0.54 | 0.52 | 1ldjA | 0.60 | 0.47 |
| 1fjrA | 0.45 | 0.43 | 3swjA | 0.51 | 0.48 | 1nyqB | 0.62 | 0.60 |
| 1g87B | 0.71 | 0.57 | 3t58B | 0.47 | 0.38 | 1ug9A | 0.53 | 0.51 |
| 1hx6B | 0.67 | 0.60 | 3t7jA | 0.65 | 0.63 | 1z1wA | 0.90 | 0.88 |
| 1iwaA | 0.92 | 0.89 | 3u07C | 0.45 | 0.41 | 2au3A | 0.65 | 0.54 |
| 1m5qH | 0.51 | 0.49 | 3u0oB | 0.79 | 0.74 | 2ii2A | 0.84 | 0.82 |
| 1mkfA | 0.43 | 0.42 | 3u9gA | 0.44 | 0.44 | 2olsA | 0.42 | 0.38 |
| 1mkmB | 0.65 | 0.58 | 3ub1D | 0.41 | 0.40 | 2ra1A | 0.42 | 0.41 |
| 1nh2D | 0.64 | 0.59 | 3uitD | 0.48 | 0.44 | 2v5dA | 0.73 | 0.58 |
| 1pprM | 0.50 | 0.48 | 3uo3A | 0.61 | 0.56 | 2xt6A | 0.62 | 0.50 |
| 1prrA | 0.49 | 0.48 | 3v7oB | 0.46 | 0.46 | 2zpaB | 0.42 | 0.36 |
| 1q19A | 0.59 | 0.48 | 3vr8B | 0.71 | 0.53 | 3apoA | 0.37 | 0.35 |
| 1qwrA | 0.83 | 0.67 | 3wkuA | 0.43 | 0.41 | 3b43A | 0.46 | 0.44 |
| 1r71B | 0.63 | 0.55 | 3zvmA | 0.42 | 0.41 | 3gf5B | 0.39 | 0.39 |
| 1rh1A | 0.41 | 0.39 | 4acoA | 0.42 | 0.37 | 3hjlA | 0.44 | 0.43 |
| 1rktA | 0.71 | 0.70 | 4ap5A | 0.69 | 0.63 | 3kq4B | 0.41 | 0.40 |
| 1s6lA | 0.45 | 0.45 | 4axdA | 0.44 | 0.40 | 3kwlA | 0.41 | 0.40 |
| 1sp3A | 0.39 | 0.38 | 4bfiB | 0.79 | 0.77 | 3ob8A | 0.89 | 0.66 |
| 1vz6A | 0.44 | 0.43 | 4bt9B | 0.54 | 0.52 | 3opfB | 0.42 | 0.43 |
| 1w3aA | 0.49 | 0.47 | 4cczA | 0.50 | 0.47 | 3p53A | 0.40 | 0.39 |
| 1wv3A | 0.50 | 0.49 | 4d0nB | 0.77 | 0.72 | 3pvlA | 0.63 | 0.56 |
| 1x7pA | 0.70 | 0.61 | 4d1iG | 0.65 | 0.64 | 3r05A | 0.33 | 0.31 |
| 1x9yA | 0.47 | 0.46 | 4dj3A | 0.60 | 0.46 | 3ubhA | 0.48 | 0.48 |
| 1y11A | 0.49 | 0.46 | 4dqaA | 0.47 | 0.46 | 3w2wA | 0.57 | 0.39 |
| 1yiqA | 0.62 | 0.58 | 4eo3A | 0.48 | 0.46 | 3zniA | 0.53 | 0.43 |
| 1zbuB | 0.56 | 0.56 | 4eogA | 0.60 | 0.42 | 4aimA | 0.73 | 0.47 |
| 1ze1A | 0.78 | 0.62 | 4etxA | 0.44 | 0.42 | 4ak1A | 0.38 | 0.38 |
| 2ablA | 0.59 | 0.56 | 4fguA | 0.90 | 0.47 | 4aq1A | 0.34 | 0.33 |
| 2ahvA | 0.74 | 0.67 | 4fkcA | 0.89 | 0.89 | 4fe9A | 0.39 | 0.38 |
| 2bkpA | 0.50 | 0.50 | 4fxkC | 0.72 | 0.58 | 4h2aA | 0.48 | 0.37 |
| 2c1yA | 0.57 | 0.52 | 4gbyA | 0.81 | 0.75 | 4i5sB | 0.49 | 0.47 |
| 2cxcA | 0.70 | 0.61 | 4ggmX | 0.46 | 0.44 | 4iggB | 0.56 | 0.36 |
| 2d1cA | 0.65 | 0.65 | 4gslA | 0.41 | 0.38 | 4j9vA | 0.58 | 0.54 |
| 2d7iA | 0.69 | 0.50 | 4gyjA | 0.80 | 0.79 | 4k3bA | 0.49 | 0.36 |
| 2e9hA | 0.49 | 0.48 | 4h3tA | 0.71 | 0.59 | 4kwuA | 0.67 | 0.66 |
| 2e9xB | 0.63 | 0.56 | 4hmoA | 0.74 | 0.74 | 4m00A | 0.39 | 0.37 |
| 2evrA | 0.66 | 0.56 | 4ie6A | 0.42 | 0.40 | 1bp1A | 0.48 | 0.45 |
| 2ew9A | 0.55 | 0.54 | 4l5gA | 0.50 | 0.50 | 1cjsA | 0.82 | 0.78 |
| 2fd5A | 0.78 | 0.78 | 4lpqA | 0.52 | 0.51 | 1ck1A | 0.88 | 0.77 |

| | | | | | | | | |
|---|---|---|---|---|---|---|---|---|
| 2gh8A | 0.56 | 0.46 | 4m8rA | 0.54 | 0.53 | 1ecrA | 0.40 | 0.38 |
| 2gt1A | 0.68 | 0.66 | 4n06B | 0.76 | 0.68 | 1f5qD | 0.80 | 0.77 |
| 2gzaC | 0.48 | 0.48 | 4nj5A | 0.51 | 0.41 | 1fa9A | 0.88 | 0.70 |
| 2hjqA | 0.52 | 0.50 | 4opaB | 0.46 | 0.45 | 1gu7A | 0.79 | 0.77 |
| 2hwjA | 0.47 | 0.46 | 4qkuB | 0.57 | 0.45 | 1itwA | 0.48 | 0.46 |
| 2ijd1 | 0.70 | 0.66 | 4up9A | 0.88 | 0.83 | 1jkiA | 0.63 | 0.54 |
| 2iu7A | 0.57 | 0.49 | 4w7sA | 0.55 | 0.53 | 1n80A | 0.37 | 0.36 |
| 2iw2A | 0.82 | 0.76 | 1bf2A | 0.79 | 0.77 | 1nzjA | 0.88 | 0.83 |
| 2jz4A | 0.49 | 0.47 | 1bhgA | 0.92 | 0.86 | 1qhdA | 0.43 | 0.40 |
| 2kdyA | 0.52 | 0.48 | 1f7uA | 0.83 | 0.70 | 1qz9A | 0.85 | 0.82 |
| 2kn4A | 0.49 | 0.48 | 1fx7A | 0.66 | 0.51 | 1sb7B | 0.50 | 0.49 |
| 2mbgA | 0.63 | 0.62 | 1griA | 0.52 | 0.48 | 1vk1A | 0.48 | 0.42 |
| 2nsfA | 0.49 | 0.47 | 1h88C | 0.57 | 0.54 | 1vrmA | 0.65 | 0.51 |
| 2nykA | 0.67 | 0.66 | 1m8pB | 0.69 | 0.56 | 1xvuA | 0.86 | 0.70 |
| 2o6yA | 0.93 | 0.81 | 1ni5A | 0.47 | 0.44 | 1yy3A | 0.52 | 0.44 |
| 2owbA | 0.94 | 0.93 | 1q25A | 0.40 | 0.38 | 1z87A | 0.40 | 0.40 |
| 2qfiA | 0.49 | 0.46 | 1uzjA | 0.48 | 0.46 | 2a1sC | 0.41 | 0.40 |
| 2qp2A | 0.50 | 0.46 | 1zpuA | 0.90 | 0.85 | 2a3lA | 0.44 | 0.44 |
| 2qygA | 0.89 | 0.88 | 1zy9A | 0.78 | 0.74 | 2bt1A | 0.50 | 0.44 |
| 2r5wB | 0.55 | 0.48 | 2b5uA | 0.37 | 0.36 | 2bydA | 0.51 | 0.46 |
| 2uu7A | 0.79 | 0.77 | 2ewfA | 0.39 | 0.37 | 2c43A | 0.51 | 0.45 |
| 2w4bA | 0.51 | 0.41 | 2piaA | 0.64 | 0.63 | 2dfyC | 0.50 | 0.46 |
| 2x7iA | 0.79 | 0.76 | 2r7dA | 0.84 | 0.60 | 2dlaA | 0.51 | 0.50 |
| 2x8kC | 0.53 | 0.50 | 2uwnA | 0.51 | 0.49 | 2g3pA | 0.41 | 0.40 |
| 2yilA | 0.56 | 0.51 | 2v0nA | 0.42 | 0.40 | 2gg6A | 0.90 | 0.86 |
| 2yrqA | 0.48 | 0.46 | 2vgmA | 0.57 | 0.52 | 2gsyE | 0.56 | 0.48 |
| 2zxcA | 0.45 | 0.45 | 2wqrB | 0.58 | 0.56 | 2gzoA | 0.49 | 0.42 |
| 3a1iA | 0.63 | 0.56 | 2y25B | 0.45 | 0.44 | 2j2cA | 0.47 | 0.49 |
| 3a45A | 0.80 | 0.77 | 2yk0A | 0.42 | 0.39 | 2kfwA | 0.44 | 0.43 |
| 3a56A | 0.43 | 0.42 | 2zzqA | 0.44 | 0.42 | 2l9yA | 0.43 | 0.41 |
| 3ajvA | 0.73 | 0.65 | 3bt1U | 0.42 | 0.42 | 2ntyB | 0.42 | 0.41 |
| 3aqkA | 0.74 | 0.67 | 3c1yA | 0.43 | 0.42 | 2r3vA | 0.70 | 0.68 |
| 3arbA | 0.86 | 0.83 | 3cw2C | 0.57 | 0.55 | 2r58A | 0.87 | 0.59 |
| 3aujG | 0.51 | 0.50 | 3f83A | 0.45 | 0.44 | 2w4mA | 0.74 | 0.72 |
| 3b2zF | 0.69 | 0.62 | 3fc3A | 0.45 | 0.44 | 2x0cA | 0.47 | 0.44 |
| 3b7wA | 0.93 | 0.86 | 3gbgA | 0.44 | 0.42 | 2y51A | 0.86 | 0.84 |
| 3bt3A | 0.74 | 0.72 | 3h5cB | 0.80 | 0.78 | 2yb0E | 0.54 | 0.47 |
| 3c4tA | 0.59 | 0.53 | 3ibjA | 0.45 | 0.42 | 2z86C | 0.37 | 0.37 |
| 3craA | 0.47 | 0.46 | 3ippB | 0.86 | 0.56 | 3afoA | 0.70 | 0.63 |
| 3d30A | 0.61 | 0.57 | 3jymB | 0.44 | 0.42 | 3bu2A | 0.50 | 0.47 |
| 3eo5A | 0.44 | 0.42 | 3kbgA | 0.86 | 0.73 | 3cvzA | 0.43 | 0.41 |
| 3errA | 0.70 | 0.62 | 3mc8A | 0.52 | 0.50 | 3dupA | 0.49 | 0.47 |
| 3g79A | 0.79 | 0.75 | 3npfA | 0.56 | 0.49 | 3eswA | 0.53 | 0.46 |
| 3h2tA | 0.41 | 0.39 | 3orjA | 0.47 | 0.41 | 3eukH | 0.59 | 0.57 |

| | | | | | | | | |
|---|---|---|---|---|---|---|---|---|
| 3hcsA | 0.48 | 0.48 | 3plaA | 0.54 | 0.43 | 3fi7A | 0.63 | 0.58 |
| 3hyiA | 0.50 | 0.49 | 3qe9Y | 0.72 | 0.66 | 3fvvA | 0.77 | 0.72 |
| 3i2dA | 0.55 | 0.47 | 3qjoA | 0.71 | 0.55 | 3gmsA | 0.88 | 0.88 |
| 3iam2 | 0.78 | 0.47 | 3qphA | 0.42 | 0.42 | 3hzzB | 0.76 | 0.74 |
| 3ifrA | 0.92 | 0.91 | 3qyeA | 0.78 | 0.76 | 3m1uA | 0.48 | 0.41 |
| 3isqA | 0.85 | 0.79 | 3rimA | 0.93 | 0.92 | 3mw8A | 0.64 | 0.63 |
| 3j7aK | 0.87 | 0.72 | 3rrpA | 0.93 | 0.85 | 3mwcA | 0.89 | 0.88 |
| 3k1rA | 0.46 | 0.46 | 3soaA | 0.65 | 0.62 | 3nsjA | 0.50 | 0.47 |
| 3k2iA | 0.59 | 0.58 | 3tixD | 0.41 | 0.40 | 3ntkA | 0.82 | 0.71 |
| 3kh5A | 0.83 | 0.57 | 3tp9A | 0.48 | 0.46 | 3oaaG | 0.76 | 0.57 |
| 3kjpA | 0.52 | 0.43 | 3ua3A | 0.44 | 0.43 | 3ptyA | 0.68 | 0.45 |
| 3kt1A | 0.43 | 0.42 | 3uj0A | 0.70 | 0.49 | 3rfyA | 0.39 | 0.38 |
| 3ktmE | 0.45 | 0.44 | 3vn4A | 0.41 | 0.39 | 3seoB | 0.46 | 0.45 |
| 3kzwA | 0.89 | 0.88 | 3vsmA | 0.76 | 0.71 | 3spgA | 0.76 | 0.50 |
| 3l76A | 0.57 | 0.50 | 3w1bA | 0.64 | 0.52 | 3u0kA | 0.56 | 0.42 |
| 3ld1A | 0.43 | 0.42 | 3zh9B | 0.53 | 0.50 | 3vlaA | 0.78 | 0.75 |
| 3lsgA | 0.85 | 0.79 | 4alzA | 0.47 | 0.47 | 3vstA | 0.49 | 0.47 |
| 3me4A | 0.63 | 0.57 | 4ax8A | 0.45 | 0.43 | 4aqfB | 0.43 | 0.42 |
| 3ml4C | 0.50 | 0.50 | 4b3iA | 0.81 | 0.58 | 4b21A | 0.85 | 0.81 |
| 3mx2B | 0.45 | 0.41 | 4bd9B | 0.44 | 0.43 | 4dt4A | 0.60 | 0.57 |
| 3mzfA | 0.78 | 0.69 | 4c0aB | 0.52 | 0.46 | 4dtfA | 0.40 | 0.39 |
| 3njaB | 0.50 | 0.49 | 4c0sA | 0.74 | 0.56 | 4ewtA | 0.81 | 0.80 |
| 3nqiA | 0.56 | 0.56 | 4dimA | 0.80 | 0.79 | 4f23A | 0.89 | 0.80 |
| 3nt8A | 0.41 | 0.39 | 4indA | 0.43 | 0.41 | 4fzbC | 0.69 | 0.50 |
| 3og5A | 0.73 | 0.63 | 4jdzB | 0.60 | 0.58 | 4g1pA | 0.78 | 0.75 |
| 3oh0A | 0.65 | 0.61 | 4kc3B | 0.72 | 0.71 | 4gfqA | 0.67 | 0.59 |
| 3pcsB | 0.47 | 0.41 | 4kikB | 0.63 | 0.42 | 4hvzA | 0.46 | 0.44 |
| 3po3S | 0.50 | 0.48 | 4lmfA | 0.63 | 0.55 | 4il6B | 0.54 | 0.45 |
| 3pxpA | 0.42 | 0.42 | 4lziA | 0.42 | 0.42 | 4jxkA | 0.90 | 0.86 |
| 3qavA | 0.81 | 0.81 | 4m9pA | 0.48 | 0.46 | 4m8mB | 0.73 | 0.58 |
| 3qf4B | 0.88 | 0.85 | 4pt5A | 0.86 | 0.63 | 4mzyA | 0.63 | 0.65 |
| 3qjjA | 0.73 | 0.64 | 4uwhA | 0.85 | 0.77 | 4onyA | 0.75 | 0.74 |
| 3qtdA | 0.44 | 0.43 | 1c1zA | 0.45 | 0.42 | 4pyhA | 0.58 | 0.54 |
| 3r6bA | 0.60 | 0.52 | 1d2pA | 0.39 | 0.39 | 4rg1A | 0.54 | 0.50 |
| 3rh7A | 0.47 | 0.44 | 1k7tA | 0.50 | 0.48 | | | |

**Table S7.** Under the sequence identity cutoff of 50%, average coverage scores of 2dom proteins, 3dom proteins, m4dom proteins, 2dis proteins and all test protein in MPDB and DEMO-lib.

| Database | 2dom (166) | 3dom (69) | m4dom (40) | 2dis (81) | Total (356) | *P*-value |
|---|---|---|---|---|---|---|
| MPDB | 0.60 | 0.60 | 0.53 | 0.62 | 0.60 | NA |
| DEMO-lib | 0.56 | 0.54 | 0.47 | 0.56 | 0.55 | 1.74E-54 |

**Table S8.** Coverage scores of each test protein in MPDB and DEMO-lib under the sequence identity cutoff of 70%.

| PDB ID | Coverage score | | PDB ID | Coverage score | | PDB ID | Coverage score | |
|---|---|---|---|---|---|---|---|---|
| | MPDB | DEMO-lib | | MPDB | DEMO-lib | | MPDB | DEMO-lib |
| 1cjyA | 0.47 | 0.41 | 3rwxA | 0.45 | 0.44 | 1kfqA | 0.92 | 0.77 |
| 1efdN | 0.85 | 0.83 | 3sb4A | 0.54 | 0.52 | 1ldjA | 0.61 | 0.47 |
| 1fjrA | 0.45 | 0.43 | 3swjA | 0.55 | 0.53 | 1nyqB | 0.62 | 0.60 |
| 1g87B | 0.78 | 0.62 | 3t58B | 0.47 | 0.38 | 1ug9A | 0.53 | 0.51 |
| 1hx6B | 0.67 | 0.60 | 3t7jA | 0.65 | 0.63 | 1z1wA | 0.90 | 0.88 |
| 1iwaA | 0.99 | 0.90 | 3u07C | 0.45 | 0.41 | 2au3A | 0.65 | 0.54 |
| 1m5qH | 0.51 | 0.49 | 3u0oB | 0.79 | 0.74 | 2ii2A | 0.84 | 0.82 |
| 1mkfA | 0.43 | 0.42 | 3u9gA | 0.44 | 0.44 | 2olsA | 0.42 | 0.38 |
| 1mkmB | 0.65 | 0.58 | 3ub1D | 0.41 | 0.40 | 2ra1A | 0.42 | 0.41 |
| 1nh2D | 0.64 | 0.59 | 3uitD | 0.48 | 0.44 | 2v5dA | 0.73 | 0.58 |
| 1pprM | 0.50 | 0.48 | 3uo3A | 0.61 | 0.56 | 2xt6A | 0.62 | 0.50 |
| 1prrA | 0.49 | 0.48 | 3v7oB | 0.46 | 0.46 | 2zpaB | 0.42 | 0.36 |
| 1q19A | 0.59 | 0.48 | 3vr8B | 0.86 | 0.63 | 3apoA | 0.37 | 0.35 |
| 1qwrA | 0.83 | 0.67 | 3wkuA | 0.43 | 0.41 | 3b43A | 0.46 | 0.44 |
| 1r71B | 0.63 | 0.55 | 3zvmA | 0.42 | 0.41 | 3gf5B | 0.39 | 0.39 |
| 1rh1A | 0.41 | 0.39 | 4acoA | 0.42 | 0.37 | 3hjlA | 0.44 | 0.43 |
| 1rktA | 0.71 | 0.70 | 4ap5A | 0.69 | 0.63 | 3kq4B | 0.41 | 0.40 |
| 1s6lA | 0.45 | 0.45 | 4axdA | 0.44 | 0.40 | 3kwlA | 0.41 | 0.40 |
| 1sp3A | 0.39 | 0.38 | 4bfiB | 0.79 | 0.77 | 3ob8A | 0.89 | 0.66 |
| 1vz6A | 0.46 | 0.43 | 4bt9B | 0.54 | 0.52 | 3opfB | 0.44 | 0.45 |
| 1w3aA | 0.49 | 0.47 | 4cczA | 0.50 | 0.47 | 3p53A | 0.40 | 0.39 |
| 1wv3A | 0.50 | 0.49 | 4d0nB | 0.77 | 0.72 | 3pvlA | 0.63 | 0.56 |
| 1x7pA | 0.70 | 0.61 | 4d1iG | 0.67 | 0.67 | 3r05A | 0.33 | 0.31 |
| 1x9yA | 0.47 | 0.46 | 4dj3A | 0.69 | 0.55 | 3ubhA | 0.48 | 0.48 |
| 1y11A | 0.49 | 0.46 | 4dqaA | 0.47 | 0.46 | 3w2wA | 0.57 | 0.39 |
| 1yiqA | 0.67 | 0.62 | 4eo3A | 0.48 | 0.46 | 3zniA | 0.74 | 0.43 |
| 1zbuB | 0.56 | 0.56 | 4eogA | 0.60 | 0.42 | 4aimA | 0.75 | 0.51 |
| 1ze1A | 0.78 | 0.62 | 4etxA | 0.44 | 0.42 | 4ak1A | 0.38 | 0.38 |
| 2ablA | 0.59 | 0.56 | 4fguA | 0.90 | 0.47 | 4aq1A | 0.34 | 0.33 |
| 2ahvA | 0.74 | 0.67 | 4fkcA | 0.90 | 0.89 | 4fe9A | 0.39 | 0.38 |
| 2bkpA | 0.50 | 0.50 | 4fxkC | 0.72 | 0.58 | 4h2aA | 0.50 | 0.37 |
| 2c1yA | 0.57 | 0.52 | 4gbyA | 0.81 | 0.75 | 4i5sB | 0.49 | 0.47 |
| 2cxcA | 0.70 | 0.61 | 4ggmX | 0.46 | 0.44 | 4iggB | 0.61 | 0.39 |
| 2d1cA | 0.65 | 0.65 | 4gslA | 0.41 | 0.38 | 4j9vA | 0.58 | 0.54 |
| 2d7iA | 0.69 | 0.50 | 4gyjA | 0.80 | 0.79 | 4k3bA | 0.49 | 0.36 |
| 2e9hA | 0.49 | 0.48 | 4h3tA | 0.71 | 0.59 | 4kwuA | 0.67 | 0.66 |
| 2e9xB | 0.63 | 0.56 | 4hmoA | 0.74 | 0.74 | 4m00A | 0.41 | 0.37 |
| 2evrA | 0.66 | 0.56 | 4ie6A | 0.42 | 0.40 | 1bp1A | 0.48 | 0.45 |
| 2ew9A | 0.55 | 0.54 | 4l5gA | 0.50 | 0.50 | 1cjsA | 0.82 | 0.78 |
| 2fd5A | 0.78 | 0.78 | 4lpqA | 0.52 | 0.51 | 1ck1A | 0.91 | 0.81 |

| | | | | | | | | |
|---|---|---|---|---|---|---|---|---|
| 2gh8A | 0.69 | 0.51 | 4m8rA | 0.54 | 0.53 | 1ecrA | 0.40 | 0.38 |
| 2gt1A | 0.68 | 0.66 | 4n06B | 0.76 | 0.68 | 1f5qD | 0.80 | 0.77 |
| 2gzaC | 0.48 | 0.48 | 4nj5A | 0.51 | 0.41 | 1fa9A | 0.92 | 0.75 |
| 2hjqA | 0.52 | 0.50 | 4opaB | 0.46 | 0.45 | 1gu7A | 0.79 | 0.77 |
| 2hwjA | 0.47 | 0.46 | 4qkuB | 0.64 | 0.51 | 1itwA | 0.48 | 0.56 |
| 2ijd1 | 0.71 | 0.66 | 4up9A | 0.88 | 0.87 | 1jkiA | 0.63 | 0.54 |
| 2iu7A | 0.72 | 0.54 | 4w7sA | 0.55 | 0.53 | 1n80A | 0.37 | 0.36 |
| 2iw2A | 0.82 | 0.76 | 1bf2A | 0.79 | 0.77 | 1nzjA | 0.88 | 0.83 |
| 2jz4A | 0.49 | 0.47 | 1bhgA | 0.92 | 0.86 | 1qhdA | 0.43 | 0.40 |
| 2kdyA | 0.56 | 0.52 | 1f7uA | 0.83 | 0.70 | 1qz9A | 0.85 | 0.82 |
| 2kn4A | 0.49 | 0.48 | 1fx7A | 0.80 | 0.59 | 1sb7B | 0.50 | 0.49 |
| 2mbgA | 0.63 | 0.62 | 1griA | 0.52 | 0.48 | 1vk1A | 0.48 | 0.42 |
| 2nsfA | 0.49 | 0.47 | 1h88C | 0.60 | 0.54 | 1vrmA | 0.65 | 0.51 |
| 2nykA | 0.67 | 0.66 | 1m8pB | 0.73 | 0.59 | 1xvuA | 0.86 | 0.70 |
| 2o6yA | 0.93 | 0.81 | 1ni5A | 0.47 | 0.44 | 1yy3A | 0.52 | 0.44 |
| 2owbA | 0.94 | 0.94 | 1q25A | 0.40 | 0.38 | 1z87A | 0.40 | 0.40 |
| 2qfiA | 0.49 | 0.46 | 1uzjA | 0.48 | 0.46 | 2a1sC | 0.41 | 0.40 |
| 2qp2A | 0.50 | 0.46 | 1zpuA | 0.90 | 0.85 | 2a3lA | 0.44 | 0.44 |
| 2qygA | 0.92 | 0.89 | 1zy9A | 0.78 | 0.74 | 2bt1A | 0.50 | 0.44 |
| 2r5wB | 0.55 | 0.48 | 2b5uA | 0.37 | 0.36 | 2bydA | 0.51 | 0.46 |
| 2uu7A | 0.84 | 0.81 | 2ewfA | 0.39 | 0.37 | 2c43A | 0.51 | 0.45 |
| 2w4bA | 0.51 | 0.41 | 2piaA | 0.64 | 0.63 | 2dfyC | 0.50 | 0.46 |
| 2x7iA | 0.79 | 0.76 | 2r7dA | 0.84 | 0.60 | 2dlaA | 0.51 | 0.50 |
| 2x8kC | 0.53 | 0.50 | 2uwnA | 0.53 | 0.49 | 2g3pA | 0.41 | 0.40 |
| 2yilA | 0.56 | 0.51 | 2v0nA | 0.42 | 0.40 | 2gg6A | 0.90 | 0.86 |
| 2yrqA | 0.49 | 0.47 | 2vgmA | 0.57 | 0.52 | 2gsyE | 0.56 | 0.48 |
| 2zxcA | 0.45 | 0.45 | 2wqrB | 0.62 | 0.57 | 2gzoA | 0.49 | 0.42 |
| 3a1iA | 0.63 | 0.56 | 2y25B | 0.47 | 0.45 | 2j2cA | 0.47 | 0.49 |
| 3a45A | 0.80 | 0.77 | 2yk0A | 0.44 | 0.42 | 2kfwA | 0.44 | 0.43 |
| 3a56A | 0.43 | 0.42 | 2zzqA | 0.44 | 0.42 | 2l9yA | 0.43 | 0.41 |
| 3ajvA | 0.73 | 0.65 | 3bt1U | 0.51 | 0.51 | 2ntyB | 0.42 | 0.41 |
| 3aqkA | 0.74 | 0.67 | 3c1yA | 0.43 | 0.42 | 2r3vA | 0.70 | 0.68 |
| 3arbA | 0.96 | 0.92 | 3cw2C | 0.57 | 0.55 | 2r58A | 0.92 | 0.70 |
| 3aujG | 0.56 | 0.55 | 3f83A | 0.48 | 0.46 | 2w4mA | 0.74 | 0.72 |
| 3b2zF | 0.78 | 0.66 | 3fc3A | 0.45 | 0.44 | 2x0cA | 0.47 | 0.44 |
| 3b7wA | 0.93 | 0.86 | 3gbgA | 0.44 | 0.42 | 2y51A | 0.86 | 0.84 |
| 3bt3A | 0.74 | 0.72 | 3h5cB | 0.80 | 0.78 | 2yb0E | 0.59 | 0.52 |
| 3c4tA | 0.59 | 0.53 | 3ibjA | 0.45 | 0.42 | 2z86C | 0.37 | 0.37 |
| 3craA | 0.47 | 0.46 | 3ippB | 0.86 | 0.56 | 3afoA | 0.70 | 0.63 |
| 3d30A | 0.61 | 0.57 | 3jymB | 0.44 | 0.42 | 3bu2A | 0.50 | 0.47 |
| 3eo5A | 0.44 | 0.42 | 3kbgA | 0.86 | 0.73 | 3cvzA | 0.43 | 0.41 |
| 3errA | 0.70 | 0.65 | 3mc8A | 0.56 | 0.55 | 3dupA | 0.49 | 0.47 |
| 3g79A | 0.79 | 0.75 | 3npfA | 0.56 | 0.49 | 3eswA | 0.53 | 0.46 |
| 3h2tA | 0.41 | 0.39 | 3orjA | 0.47 | 0.41 | 3eukH | 0.59 | 0.57 |

| | | | | | | | | |
|---|---|---|---|---|---|---|---|---|
| 3hcsA | 0.52 | 0.48 | 3plaA | 0.56 | 0.45 | 3fi7A | 0.63 | 0.58 |
| 3hyiA | 0.50 | 0.49 | 3qe9Y | 0.72 | 0.66 | 3fvvA | 0.77 | 0.72 |
| 3i2dA | 0.55 | 0.47 | 3qjoA | 0.71 | 0.55 | 3gmsA | 0.88 | 0.88 |
| 3iam2 | 0.78 | 0.47 | 3qphA | 0.45 | 0.44 | 3hzzB | 0.79 | 0.76 |
| 3ifrA | 0.92 | 0.91 | 3qyeA | 0.79 | 0.78 | 3m1uA | 0.48 | 0.41 |
| 3isqA | 0.85 | 0.79 | 3rimA | 0.93 | 0.92 | 3mw8A | 0.64 | 0.63 |
| 3j7aK | 0.92 | 0.76 | 3rrpA | 0.93 | 0.89 | 3mwcA | 0.89 | 0.88 |
| 3k1rA | 0.46 | 0.46 | 3soaA | 0.66 | 0.63 | 3nsjA | 0.50 | 0.47 |
| 3k2iA | 0.66 | 0.62 | 3tixD | 0.41 | 0.40 | 3ntkA | 0.82 | 0.71 |
| 3kh5A | 0.83 | 0.57 | 3tp9A | 0.48 | 0.46 | 3oaaG | 0.76 | 0.57 |
| 3kjpA | 0.52 | 0.43 | 3ua3A | 0.44 | 0.43 | 3ptyA | 0.68 | 0.45 |
| 3kt1A | 0.43 | 0.42 | 3uj0A | 0.70 | 0.50 | 3rfyA | 0.39 | 0.38 |
| 3ktmE | 0.45 | 0.44 | 3vn4A | 0.41 | 0.39 | 3seoB | 0.46 | 0.45 |
| 3kzwA | 0.89 | 0.88 | 3vsmA | 0.76 | 0.71 | 3spgA | 0.83 | 0.56 |
| 3l76A | 0.57 | 0.50 | 3w1bA | 0.64 | 0.52 | 3u0kA | 0.57 | 0.46 |
| 3ld1A | 0.43 | 0.42 | 3zh9B | 0.53 | 0.50 | 3vlaA | 0.78 | 0.75 |
| 3lsgA | 0.85 | 0.79 | 4alzA | 0.47 | 0.47 | 3vstA | 0.49 | 0.47 |
| 3me4A | 0.63 | 0.57 | 4ax8A | 0.45 | 0.43 | 4aqfB | 0.63 | 0.57 |
| 3ml4C | 0.50 | 0.50 | 4b3iA | 0.81 | 0.58 | 4b21A | 0.85 | 0.81 |
| 3mx2B | 0.45 | 0.41 | 4bd9B | 0.44 | 0.43 | 4dt4A | 0.62 | 0.57 |
| 3mzfA | 0.83 | 0.76 | 4c0aB | 0.52 | 0.46 | 4dtfA | 0.40 | 0.39 |
| 3njaB | 0.50 | 0.49 | 4c0sA | 0.76 | 0.64 | 4ewtA | 0.81 | 0.80 |
| 3nqiA | 0.56 | 0.56 | 4dimA | 0.80 | 0.79 | 4f23A | 0.89 | 0.80 |
| 3nt8A | 0.41 | 0.39 | 4indA | 0.43 | 0.41 | 4fzbC | 0.69 | 0.50 |
| 3og5A | 0.76 | 0.68 | 4jdzB | 0.61 | 0.59 | 4g1pA | 0.81 | 0.78 |
| 3oh0A | 0.65 | 0.61 | 4kc3B | 0.75 | 0.71 | 4gfqA | 0.68 | 0.59 |
| 3pcsB | 0.47 | 0.41 | 4kikB | 0.67 | 0.51 | 4hvzA | 0.46 | 0.44 |
| 3po3S | 0.50 | 0.48 | 4lmfA | 0.64 | 0.56 | 4il6B | 0.54 | 0.45 |
| 3pxpA | 0.42 | 0.42 | 4lziA | 0.42 | 0.42 | 4jxkA | 0.90 | 0.86 |
| 3qavA | 0.81 | 0.81 | 4m9pA | 0.48 | 0.46 | 4m8mB | 0.73 | 0.58 |
| 3qf4B | 0.88 | 0.85 | 4pt5A | 0.86 | 0.63 | 4mzyA | 0.63 | 0.65 |
| 3qjjA | 0.73 | 0.64 | 4uwhA | 0.86 | 0.81 | 4onyA | 0.75 | 0.74 |
| 3qtdA | 0.44 | 0.43 | 1c1zA | 0.45 | 0.42 | 4pyhA | 0.58 | 0.54 |
| 3r6bA | 0.63 | 0.54 | 1d2pA | 0.41 | 0.39 | 4rg1A | 0.54 | 0.50 |
| 3rh7A | 0.47 | 0.44 | 1k7tA | 0.50 | 0.48 | | | |

**Table S9.** Under the sequence identity cutoff of 70%, average coverage scores of 2dom proteins, 3dom proteins, m4dom proteins, 2dis proteins and all test protein in MPDB and DEMO-lib.

| Database | 2dom (166) | 3dom (69) | m4dom (40) | 2dis (81) | Total (356) | *P*-value |
|---|---|---|---|---|---|---|
| MPDB | 0.61 | 0.61 | 0.54 | 0.62 | 0.61 | NA |
| DEMO-lib | 0.56 | 0.55 | 0.47 | 0.57 | 0.55 | 7.81E-54 |

**Table S10.** TM-score between the structural analogue with the highest $LS_{score}$ and the native structure of test proteins under the sequence identity cutoff of 30%, 50% and 70%, respectively.

| PDB ID | TM-score | | | PDB ID | TM-score | | | PDB ID | TM-score | | |
|---|---|---|---|---|---|---|---|---|---|---|---|
| | 30% | 50% | 70% | | 30% | 50% | 70% | | 30% | 50% | 70% |
| 1cjyA | 0.25 | 0.81 | 0.81 | 3rwxA | 0.31 | 0.31 | 0.31 | 1kfqA | 0.77 | 0.89 | 0.91 |
| 1efdN | 0.90 | 0.91 | 0.91 | 3sb4A | 0.45 | 0.45 | 0.45 | 1ldjA | 0.42 | 0.87 | 0.87 |
| 1fjrA | 0.33 | 0.36 | 0.36 | 3swjA | 0.53 | 0.53 | 0.91 | 1nyqB | 0.29 | 0.87 | 0.87 |
| 1g87B | 0.61 | 0.94 | 0.94 | 3t58B | 0.48 | 0.48 | 0.48 | 1ug9A | 0.52 | 0.52 | 0.52 |
| 1hx6B | 0.70 | 0.70 | 0.70 | 3t7jA | 0.69 | 0.69 | 0.69 | 1z1wA | 0.87 | 0.87 | 0.87 |
| 1iwaA | 0.87 | 0.93 | 0.99 | 3u07C | 0.67 | 0.67 | 0.67 | 2au3A | 0.65 | 0.65 | 0.65 |
| 1m5qH | 0.39 | 0.39 | 0.39 | 3u0oB | 0.80 | 0.89 | 0.89 | 2ii2A | 0.81 | 0.81 | 0.81 |
| 1mkfA | 0.59 | 0.59 | 0.59 | 3u9gA | 0.33 | 0.27 | 0.27 | 2olsA | 0.41 | 0.41 | 0.41 |
| 1mkmB | 0.64 | 0.64 | 0.64 | 3ub1D | 0.34 | 0.34 | 0.34 | 2ra1A | 0.26 | 0.33 | 0.33 |
| 1nh2D | 0.47 | 0.95 | 0.95 | 3uitD | 0.31 | 0.48 | 0.48 | 2v5dA | 0.58 | 0.72 | 0.72 |
| 1pprM | 0.39 | 0.90 | 0.90 | 3uo3A | 0.76 | 0.76 | 0.76 | 2xt6A | 0.26 | 0.75 | 0.75 |
| 1prrA | 0.43 | 0.43 | 0.43 | 3v7oB | 0.40 | 0.40 | 0.40 | 2zpaB | 0.82 | 0.82 | 0.82 |
| 1q19A | 0.84 | 0.84 | 0.84 | 3vr8B | 0.87 | 0.89 | 0.94 | 3apoA | 0.28 | 0.28 | 0.28 |
| 1qwrA | 0.84 | 0.84 | 0.84 | 3wkuA | 0.29 | 0.29 | 0.29 | 3b43A | 0.35 | 0.35 | 0.35 |
| 1r71B | 0.78 | 0.78 | 0.78 | 3zvmA | 0.32 | 0.32 | 0.32 | 3gf5B | 0.21 | 0.30 | 0.30 |
| 1rh1A | 0.45 | 0.45 | 0.45 | 4acoA | 0.72 | 0.72 | 0.72 | 3hjlA | 0.33 | 0.33 | 0.33 |
| 1rktA | 0.71 | 0.71 | 0.71 | 4ap5A | 0.69 | 0.93 | 0.93 | 3kq4B | 0.36 | 0.36 | 0.36 |
| 1s6lA | 0.32 | 0.38 | 0.38 | 4axdA | 0.32 | 0.77 | 0.77 | 3kwlA | 0.41 | 0.35 | 0.35 |
| 1sp3A | 0.35 | 0.35 | 0.35 | 4bfiB | 0.78 | 0.78 | 0.78 | 3ob8A | 0.52 | 0.89 | 0.89 |
| 1vz6A | 0.33 | 0.33 | 0.52 | 4bt9B | 0.40 | 0.40 | 0.40 | 3opfB | 0.26 | 0.55 | 0.55 |
| 1w3aA | 0.39 | 0.39 | 0.39 | 4cczA | 0.75 | 0.75 | 0.75 | 3p53A | 0.43 | 0.43 | 0.43 |
| 1wv3A | 0.76 | 0.76 | 0.76 | 4d0nB | 0.77 | 0.82 | 0.82 | 3pvlA | 0.40 | 0.84 | 0.84 |
| 1x7pA | 0.70 | 0.79 | 0.79 | 4d1iG | 0.69 | 0.69 | 0.90 | 3r05A | 0.29 | 0.29 | 0.29 |
| 1x9yA | 0.37 | 0.37 | 0.37 | 4dj3A | 0.79 | 0.79 | 0.85 | 3ubhA | 0.46 | 0.47 | 0.47 |
| 1y11A | 0.49 | 0.49 | 0.49 | 4dqaA | 0.35 | 0.35 | 0.35 | 3w2wA | 0.62 | 0.62 | 0.62 |
| 1yiqA | 0.45 | 0.92 | 0.94 | 4eo3A | 0.43 | 0.43 | 0.43 | 3zniA | 0.27 | 0.71 | 0.71 |
| 1zbuB | 0.46 | 0.55 | 0.55 | 4eogA | 0.77 | 0.77 | 0.77 | 4aimA | 0.31 | 0.31 | 0.77 |
| 1ze1A | 0.81 | 0.78 | 0.78 | 4etxA | 0.33 | 0.33 | 0.33 | 4ak1A | 0.18 | 0.18 | 0.18 |
| 2ablA | 0.48 | 0.58 | 0.58 | 4fguA | 0.39 | 0.91 | 0.91 | 4aq1A | 0.25 | 0.25 | 0.25 |
| 2ahvA | 0.77 | 0.77 | 0.77 | 4fkcA | 0.88 | 0.89 | 0.93 | 4fe9A | 0.30 | 0.33 | 0.33 |
| 2bkpA | 0.44 | 0.41 | 0.41 | 4fxkC | 0.83 | 0.83 | 0.83 | 4h2aA | 0.29 | 0.70 | 0.70 |
| 2c1yA | 0.58 | 0.57 | 0.57 | 4gbyA | 0.87 | 0.87 | 0.87 | 4i5sB | 0.44 | 0.44 | 0.44 |
| 2cxcA | 0.74 | 0.74 | 0.74 | 4ggmX | 0.38 | 0.37 | 0.37 | 4iggB | 0.62 | 0.62 | 0.62 |
| 2d1cA | 0.59 | 0.59 | 0.59 | 4gslA | 0.37 | 0.37 | 0.37 | 4j9vA | 0.61 | 0.61 | 0.61 |
| 2d7iA | 0.35 | 0.74 | 0.74 | 4gyjA | 0.82 | 0.82 | 0.82 | 4k3bA | 0.38 | 0.54 | 0.54 |
| 2e9hA | 0.45 | 0.42 | 0.42 | 4h3tA | 0.74 | 0.90 | 0.90 | 4kwuA | 0.63 | 0.63 | 0.63 |
| 2e9xB | 0.73 | 0.73 | 0.73 | 4hmoA | 0.76 | 0.76 | 0.76 | 4m00A | 0.31 | 0.31 | 0.31 |
| 2evrA | 0.76 | 0.81 | 0.81 | 4ie6A | 0.35 | 0.35 | 0.35 | 1bp1A | 0.72 | 0.74 | 0.74 |
| 2ew9A | 0.54 | 0.54 | 0.54 | 4l5gA | 0.57 | 0.57 | 0.57 | 1cjsA | 0.83 | 0.83 | 0.83 |
| 2fd5A | 0.82 | 0.82 | 0.82 | 4lpqA | 0.49 | 0.49 | 0.49 | 1ck1A | 0.83 | 0.89 | 0.92 |

| | | | | | | | | | | | |
|---|---|---|---|---|---|---|---|---|---|---|---|
| 2gh8A | 0.59 | 0.77 | 0.94 | 4m8rA | 0.48 | 0.48 | 0.48 | 1ecrA | 0.33 | 0.43 | 0.43 |
| 2gt1A | 0.84 | 0.84 | 0.84 | 4n06B | 0.86 | 0.86 | 0.86 | 1f5qD | 0.83 | 0.83 | 0.83 |
| 2gzaC | 0.48 | 0.51 | 0.51 | 4nj5A | 0.25 | 0.86 | 0.86 | 1fa9A | 0.52 | 0.95 | 0.96 |
| 2hjqA | 0.33 | 0.38 | 0.38 | 4opaB | 0.27 | 0.38 | 0.38 | 1gu7A | 0.79 | 0.86 | 0.86 |
| 2hwjA | 0.45 | 0.37 | 0.37 | 4qkuB | 0.67 | 0.67 | 0.94 | 1itwA | 0.48 | 0.48 | 0.48 |
| 2ijd1 | 0.67 | 0.70 | 0.70 | 4up9A | 0.73 | 0.88 | 0.90 | 1jkiA | 0.70 | 0.94 | 0.94 |
| 2iu7A | 0.49 | 0.84 | 0.98 | 4w7sA | 0.52 | 0.52 | 0.52 | 1n80A | 0.30 | 0.30 | 0.30 |
| 2iw2A | 0.82 | 0.82 | 0.82 | 1bf2A | 0.76 | 0.85 | 0.85 | 1nzjA | 0.75 | 0.89 | 0.89 |
| 2jz4A | 0.30 | 0.47 | 0.47 | 1bhgA | 0.84 | 0.92 | 0.92 | 1qhdA | 0.52 | 0.52 | 0.52 |
| 2kdyA | 0.52 | 0.52 | 0.64 | 1f7uA | 0.85 | 0.84 | 0.84 | 1qz9A | 0.92 | 0.92 | 0.92 |
| 2kn4A | 0.47 | 0.35 | 0.46 | 1fx7A | 0.75 | 0.75 | 0.93 | 1sb7B | 0.78 | 0.78 | 0.78 |
| 2mbgA | 0.61 | 0.61 | 0.61 | 1griA | 0.42 | 0.42 | 0.42 | 1vk1A | 0.73 | 0.73 | 0.73 |
| 2nsfA | 0.31 | 0.31 | 0.31 | 1h88C | 0.35 | 0.64 | 0.64 | 1vrmA | 0.86 | 0.91 | 0.91 |
| 2nykA | 0.71 | 0.71 | 0.71 | 1m8pB | 0.62 | 0.62 | 0.62 | 1xvuA | 0.86 | 0.86 | 0.86 |
| 2o6yA | 0.87 | 0.95 | 0.95 | 1ni5A | 0.61 | 0.61 | 0.61 | 1yy3A | 0.30 | 0.78 | 0.78 |
| 2owbA | 0.91 | 0.92 | 0.95 | 1q25A | 0.44 | 0.44 | 0.44 | 1z87A | 0.30 | 0.30 | 0.30 |
| 2qfiA | 0.36 | 0.66 | 0.66 | 1uzjA | 0.38 | 0.63 | 0.63 | 2a1sC | 0.37 | 0.39 | 0.39 |
| 2qp2A | 0.50 | 0.50 | 0.50 | 1zpuA | 0.91 | 0.90 | 0.90 | 2a3lA | 0.51 | 0.51 | 0.51 |
| 2qygA | 0.90 | 0.90 | 0.99 | 1zy9A | 0.77 | 0.77 | 0.77 | 2bt1A | 0.70 | 0.70 | 0.70 |
| 2r5wB | 0.30 | 0.91 | 0.91 | 2b5uA | 0.47 | 0.47 | 0.47 | 2bydA | 0.72 | 0.72 | 0.72 |
| 2uu7A | 0.79 | 0.88 | 0.92 | 2ewfA | 0.35 | 0.35 | 0.35 | 2c43A | 0.74 | 0.74 | 0.74 |
| 2w4bA | 0.30 | 0.87 | 0.87 | 2piaA | 0.61 | 0.70 | 0.70 | 2dfyC | 0.46 | 0.52 | 0.52 |
| 2x7iA | 0.82 | 0.82 | 0.82 | 2r7dA | 0.86 | 0.86 | 0.86 | 2dlaA | 0.62 | 0.62 | 0.62 |
| 2x8kC | 0.82 | 0.82 | 0.82 | 2uwnA | 0.45 | 0.45 | 0.45 | 2g3pA | 0.31 | 0.34 | 0.34 |
| 2yilA | 0.46 | 0.57 | 0.57 | 2v0nA | 0.27 | 0.35 | 0.35 | 2gg6A | 0.88 | 0.95 | 0.95 |
| 2yrqA | 0.37 | 0.37 | 0.37 | 2vgmA | 0.51 | 0.57 | 0.57 | 2gsyE | 0.31 | 0.90 | 0.90 |
| 2zxcA | 0.26 | 0.93 | 0.93 | 2wqrB | 0.58 | 0.53 | 0.53 | 2gzoA | 0.61 | 0.65 | 0.65 |
| 3a1iA | 0.75 | 0.75 | 0.75 | 2y25B | 0.49 | 0.49 | 0.49 | 2j2cA | 0.43 | 0.82 | 0.82 |
| 3a45A | 0.80 | 0.80 | 0.80 | 2yk0A | 0.45 | 0.37 | 0.37 | 2kfwA | 0.43 | 0.43 | 0.43 |
| 3a56A | 0.43 | 0.43 | 0.43 | 2zzqA | 0.41 | 0.41 | 0.41 | 2l9yA | 0.36 | 0.36 | 0.36 |
| 3ajvA | 0.73 | 0.78 | 0.78 | 3bt1U | 0.35 | 0.35 | 0.86 | 2ntyB | 0.38 | 0.38 | 0.38 |
| 3aqkA | 0.78 | 0.78 | 0.78 | 3c1yA | 0.44 | 0.44 | 0.44 | 2r3vA | 0.69 | 0.69 | 0.69 |
| 3arbA | 0.86 | 0.93 | 0.97 | 3cw2C | 0.66 | 0.65 | 0.65 | 2r58A | 0.25 | 0.64 | 0.98 |
| 3aujG | 0.35 | 0.42 | 0.95 | 3f83A | 0.39 | 0.55 | 0.55 | 2w4mA | 0.71 | 0.71 | 0.71 |
| 3b2zF | 0.64 | 0.85 | 0.92 | 3fc3A | 0.27 | 0.35 | 0.35 | 2x0cA | 0.59 | 0.59 | 0.59 |
| 3b7wA | 0.78 | 0.78 | 0.78 | 3gbgA | 0.36 | 0.36 | 0.36 | 2y51A | 0.85 | 0.94 | 0.94 |
| 3bt3A | 0.74 | 0.74 | 0.74 | 3h5cB | 0.79 | 0.79 | 0.79 | 2yb0E | 0.36 | 0.81 | 0.81 |
| 3c4tA | 0.59 | 0.59 | 0.59 | 3ibjA | 0.47 | 0.47 | 0.47 | 2z86C | 0.33 | 0.33 | 0.33 |
| 3craA | 0.35 | 0.35 | 0.35 | 3ippB | 0.39 | 0.92 | 0.92 | 3afoA | 0.75 | 0.74 | 0.74 |
| 3d30A | 0.81 | 0.84 | 0.84 | 3jymB | 0.51 | 0.51 | 0.51 | 3bu2A | 0.41 | 0.50 | 0.50 |
| 3eo5A | 0.27 | 0.36 | 0.36 | 3kbgA | 0.25 | 0.90 | 0.90 | 3cvzA | 0.32 | 0.32 | 0.32 |
| 3errA | 0.70 | 0.70 | 0.70 | 3mc8A | 0.49 | 0.49 | 0.74 | 3dupA | 0.48 | 0.49 | 0.49 |
| 3g79A | 0.80 | 0.75 | 0.75 | 3npfA | 0.75 | 0.75 | 0.75 | 3eswA | 0.42 | 0.77 | 0.77 |
| 3h2tA | 0.35 | 0.34 | 0.34 | 3orjA | 0.57 | 0.57 | 0.57 | 3eukH | 0.61 | 0.61 | 0.61 |

| | | | | | | | | | | | |
|---|---|---|---|---|---|---|---|---|---|---|---|
| 3hcsA | 0.45 | 0.36 | 0.88 | 3plaA | 0.58 | 0.60 | 0.60 | 3fi7A | 0.66 | 0.77 | 0.77 |
| 3hyiA | 0.51 | 0.51 | 0.51 | 3qe9Y | 0.75 | 0.75 | 0.75 | 3fvvA | 0.81 | 0.81 | 0.81 |
| 3i2dA | 0.73 | 0.77 | 0.77 | 3qjoA | 0.66 | 0.84 | 0.84 | 3gmsA | 0.90 | 0.90 | 0.90 |
| 3iam2 | 0.74 | 0.89 | 0.89 | 3qphA | 0.41 | 0.41 | 0.67 | 3hzzB | 0.74 | 0.84 | 0.93 |
| 3ifrA | 0.92 | 0.93 | 0.93 | 3qyeA | 0.75 | 0.82 | 0.92 | 3m1uA | 0.54 | 0.90 | 0.90 |
| 3isqA | 0.86 | 0.88 | 0.88 | 3rimA | 0.83 | 0.94 | 0.94 | 3mw8A | 0.63 | 0.67 | 0.67 |
| 3j7aK | 0.86 | 0.96 | 0.98 | 3rrpA | 0.77 | 0.93 | 0.93 | 3mwcA | 0.89 | 0.89 | 0.89 |
| 3k1rA | 0.41 | 0.41 | 0.41 | 3soaA | 0.56 | 0.56 | 0.56 | 3nsjA | 0.37 | 0.37 | 0.37 |
| 3k2iA | 0.49 | 0.49 | 0.95 | 3tixD | 0.32 | 0.44 | 0.44 | 3ntkA | 0.80 | 0.80 | 0.80 |
| 3kh5A | 0.84 | 0.84 | 0.84 | 3tp9A | 0.38 | 0.38 | 0.38 | 3oaaG | 0.80 | 0.85 | 0.85 |
| 3kjpA | 0.57 | 0.57 | 0.57 | 3ua3A | 0.41 | 0.48 | 0.48 | 3ptyA | 0.36 | 0.91 | 0.91 |
| 3kt1A | 0.70 | 0.70 | 0.70 | 3uj0A | 0.71 | 0.74 | 0.74 | 3rfyA | 0.34 | 0.35 | 0.35 |
| 3ktmE | 0.32 | 0.32 | 0.32 | 3vn4A | 0.36 | 0.36 | 0.36 | 3seoB | 0.32 | 0.32 | 0.32 |
| 3kzwA | 0.89 | 0.90 | 0.90 | 3vsmA | 0.76 | 0.76 | 0.76 | 3spgA | 0.78 | 0.80 | 0.96 |
| 3l76A | 0.58 | 0.63 | 0.63 | 3w1bA | 0.64 | 0.64 | 0.64 | 3u0kA | 0.51 | 0.52 | 0.55 |
| 3ld1A | 0.34 | 0.34 | 0.34 | 3zh9B | 0.58 | 0.58 | 0.58 | 3vlaA | 0.68 | 0.87 | 0.87 |
| 3lsgA | 0.80 | 0.94 | 0.94 | 4alzA | 0.60 | 0.60 | 0.60 | 3vstA | 0.56 | 0.56 | 0.56 |
| 3me4A | 0.60 | 0.65 | 0.65 | 4ax8A | 0.36 | 0.36 | 0.36 | 4aqfB | 0.34 | 0.74 | 0.90 |
| 3ml4C | 0.47 | 0.47 | 0.47 | 4b3iA | 0.84 | 0.88 | 0.88 | 4b21A | 0.83 | 0.90 | 0.90 |
| 3mx2B | 0.29 | 0.32 | 0.32 | 4bd9B | 0.32 | 0.32 | 0.32 | 4dt4A | 0.75 | 0.75 | 0.77 |
| 3mzfA | 0.79 | 0.82 | 0.92 | 4c0aB | 0.46 | 0.59 | 0.59 | 4dtfA | 0.32 | 0.32 | 0.32 |
| 3njaB | 0.45 | 0.45 | 0.45 | 4c0sA | 0.49 | 0.81 | 0.53 | 4ewtA | 0.76 | 0.81 | 0.81 |
| 3nqiA | 0.68 | 0.68 | 0.68 | 4dimA | 0.78 | 0.78 | 0.78 | 4f23A | 0.59 | 0.90 | 0.90 |
| 3nt8A | 0.34 | 0.34 | 0.34 | 4indA | 0.40 | 0.40 | 0.40 | 4fzbC | 0.82 | 0.82 | 0.82 |
| 3og5A | 0.80 | 0.81 | 0.81 | 4jdzB | 0.59 | 0.59 | 0.59 | 4g1pA | 0.81 | 0.94 | 0.97 |
| 3oh0A | 0.58 | 0.58 | 0.58 | 4kc3B | 0.72 | 0.72 | 0.72 | 4gfqA | 0.63 | 0.65 | 0.65 |
| 3pcsB | 0.66 | 0.66 | 0.66 | 4kikB | 0.65 | 0.65 | 0.86 | 4hvzA | 0.28 | 0.28 | 0.28 |
| 3po3S | 0.37 | 0.72 | 0.72 | 4lmfA | 0.46 | 0.65 | 0.65 | 4il6B | 0.48 | 0.48 | 0.48 |
| 3pxpA | 0.29 | 0.29 | 0.29 | 4lziA | 0.27 | 0.27 | 0.27 | 4jxkA | 0.84 | 0.93 | 0.93 |
| 3qavA | 0.81 | 0.81 | 0.81 | 4m9pA | 0.32 | 0.32 | 0.32 | 4m8mB | 0.23 | 0.80 | 0.80 |
| 3qf4B | 0.86 | 0.86 | 0.86 | 4pt5A | 0.77 | 0.87 | 0.87 | 4mzyA | 0.73 | 0.83 | 0.83 |
| 3qjjA | 0.85 | 0.88 | 0.88 | 4uwhA | 0.86 | 0.86 | 0.90 | 4onyA | 0.64 | 0.93 | 0.93 |
| 3qtdA | 0.94 | 0.94 | 0.94 | 1c1zA | 0.49 | 0.44 | 0.44 | 4pyhA | 0.53 | 0.53 | 0.53 |
| 3r6bA | 0.32 | 0.78 | 0.78 | 1d2pA | 0.38 | 0.38 | 0.38 | 4rg1A | 0.34 | 0.80 | 0.80 |
| 3rh7A | 0.45 | 0.45 | 0.45 | 1k7tA | 0.39 | 0.39 | 0.39 | | | | |

**Table S11.** Under the sequence identity (SeqId) cutoff of 30%, 50% and 70%, average TM-score of the structural analogue with the highest $LS_{score}$. #TM-score⩾0.5 represents the number of the structural analogues with TM-score⩾0.5.

| SeqId cutoff | 2dom (166) | 3dom (69) | m4dom (40) | 2dis (81) | Total (356) | #TM-score⩾0.5 |
|---|---|---|---|---|---|---|
| 30% | 0.58 | 0.55 | 0.44 | 0.59 | 0.56 | 192 |
| 50% | 0.65 | 0.61 | 0.53 | 0.69 | 0.64 | 239 |
| 70% | 0.67 | 0.63 | 0.55 | 0.70 | 0.65 | 247 |

**Table S12.** The $LS_{\text{score}}$ of the top 1 structural analogue detected under the sequence identity cutoff of 30%, 50% and 70%, respectively.

| PDB ID | $LS_{\text{score}}$ | | | PDB ID | $LS_{\text{score}}$ | | | PDB ID | $LS_{\text{score}}$ | | |
|---|---|---|---|---|---|---|---|---|---|---|---|
| | 30% | 50% | 70% | | 30% | 50% | 70% | | 30% | 50% | 70% |
| 1cjyA | 0.46 | 0.87 | 0.87 | 3rwxA | 0.61 | 0.61 | 0.61 | 1kfqA | 0.68 | 0.90 | 0.91 |
| 1efdN | 0.86 | 0.87 | 0.87 | 3sb4A | 0.67 | 0.67 | 0.67 | 1ldjA | 0.60 | 0.82 | 0.82 |
| 1fjrA | 0.41 | 0.43 | 0.43 | 3swjA | 0.79 | 0.79 | 0.85 | 1nyqB | 0.49 | 0.90 | 0.90 |
| 1g87B | 0.60 | 0.93 | 0.93 | 3t58B | 0.60 | 0.60 | 0.60 | 1ug9A | 0.45 | 0.45 | 0.45 |
| 1hx6B | 0.69 | 0.69 | 0.69 | 3t7jA | 0.64 | 0.64 | 0.64 | 1z1wA | 0.88 | 0.88 | 0.88 |
| 1iwaA | 0.82 | 0.92 | 0.99 | 3u07C | 0.79 | 0.79 | 0.79 | 2au3A | 0.55 | 0.55 | 0.55 |
| 1m5qH | 0.59 | 0.59 | 0.59 | 3u0oB | 0.79 | 0.86 | 0.86 | 2ii2A | 0.84 | 0.84 | 0.84 |
| 1mkfA | 0.53 | 0.53 | 0.53 | 3u9gA | 0.50 | 0.53 | 0.53 | 2olsA | 0.73 | 0.73 | 0.73 |
| 1mkmB | 0.84 | 0.84 | 0.84 | 3ub1D | 0.60 | 0.60 | 0.60 | 2ra1A | 0.54 | 0.58 | 0.58 |
| 1nh2D | 0.65 | 0.92 | 0.92 | 3uitD | 0.52 | 0.74 | 0.74 | 2v5dA | 0.62 | 0.75 | 0.75 |
| 1pprM | 0.46 | 0.87 | 0.87 | 3uo3A | 0.75 | 0.75 | 0.75 | 2xt6A | 0.45 | 0.55 | 0.55 |
| 1prrA | 0.73 | 0.73 | 0.73 | 3v7oB | 0.58 | 0.58 | 0.58 | 2zpaB | 0.88 | 0.88 | 0.88 |
| 1q19A | 0.76 | 0.76 | 0.76 | 3vr8B | 0.83 | 0.87 | 0.94 | 3apoA | 0.40 | 0.40 | 0.40 |
| 1qwrA | 0.79 | 0.79 | 0.79 | 3wkuA | 0.43 | 0.43 | 0.43 | 3b43A | 0.82 | 0.82 | 0.82 |
| 1r71B | 0.71 | 0.71 | 0.71 | 3zvmA | 0.57 | 0.57 | 0.57 | 3gf5B | 0.41 | 0.44 | 0.44 |
| 1rh1A | 0.40 | 0.40 | 0.40 | 4acoA | 0.63 | 0.63 | 0.63 | 3hjlA | 0.50 | 0.50 | 0.50 |
| 1rktA | 0.74 | 0.74 | 0.74 | 4ap5A | 0.66 | 0.92 | 0.92 | 3kq4B | 0.61 | 0.65 | 0.65 |
| 1s6lA | 0.49 | 0.49 | 0.49 | 4axdA | 0.40 | 0.71 | 0.71 | 3kwlA | 0.45 | 0.44 | 0.44 |
| 1sp3A | 0.39 | 0.39 | 0.39 | 4bfiB | 0.81 | 0.81 | 0.81 | 3ob8A | 0.49 | 0.82 | 0.82 |
| 1vz6A | 0.43 | 0.43 | 0.48 | 4bt9B | 0.61 | 0.61 | 0.61 | 3opfB | 0.40 | 0.83 | 0.83 |
| 1w3aA | 0.55 | 0.55 | 0.55 | 4cczA | 0.80 | 0.80 | 0.80 | 3p53A | 0.84 | 0.84 | 0.84 |
| 1wv3A | 0.78 | 0.78 | 0.78 | 4d0nB | 0.86 | 0.86 | 0.86 | 3pvlA | 0.45 | 0.88 | 0.88 |
| 1x7pA | 0.78 | 0.84 | 0.84 | 4d1iG | 0.54 | 0.54 | 0.83 | 3r05A | 0.36 | 0.36 | 0.36 |
| 1x9yA | 0.47 | 0.47 | 0.47 | 4dj3A | 0.74 | 0.74 | 0.85 | 3ubhA | 0.80 | 0.82 | 0.82 |
| 1y11A | 0.52 | 0.52 | 0.52 | 4dqaA | 0.64 | 0.64 | 0.64 | 3w2wA | 0.52 | 0.52 | 0.52 |
| 1yiqA | 0.58 | 0.83 | 0.90 | 4eo3A | 0.48 | 0.48 | 0.48 | 3zniA | 0.47 | 0.48 | 0.48 |
| 1zbuB | 0.52 | 0.54 | 0.54 | 4eogA | 0.72 | 0.72 | 0.72 | 4aimA | 0.42 | 0.42 | 0.81 |
| 1ze1A | 0.80 | 0.81 | 0.81 | 4etxA | 0.64 | 0.64 | 0.64 | 4ak1A | 0.55 | 0.55 | 0.55 |
| 2ablA | 0.75 | 0.83 | 0.83 | 4fguA | 0.47 | 0.86 | 0.86 | 4aq1A | 0.54 | 0.54 | 0.54 |
| 2ahvA | 0.77 | 0.77 | 0.77 | 4fkcA | 0.85 | 0.88 | 0.90 | 4fe9A | 0.44 | 0.46 | 0.46 |
| 2bkpA | 0.57 | 0.59 | 0.59 | 4fxkC | 0.80 | 0.81 | 0.81 | 4h2aA | 0.41 | 0.71 | 0.71 |
| 2c1yA | 0.55 | 0.83 | 0.83 | 4gbyA | 0.84 | 0.84 | 0.84 | 4i5sB | 0.55 | 0.55 | 0.55 |
| 2cxcA | 0.75 | 0.75 | 0.75 | 4ggmX | 0.56 | 0.57 | 0.57 | 4iggB | 0.72 | 0.72 | 0.72 |
| 2d1cA | 0.48 | 0.48 | 0.48 | 4gslA | 0.42 | 0.42 | 0.42 | 4j9vA | 0.72 | 0.72 | 0.72 |
| 2d7iA | 0.46 | 0.84 | 0.84 | 4gyjA | 0.77 | 0.77 | 0.77 | 4k3bA | 0.49 | 0.86 | 0.86 |
| 2e9hA | 0.48 | 0.51 | 0.51 | 4h3tA | 0.73 | 0.82 | 0.82 | 4kwuA | 0.47 | 0.47 | 0.47 |
| 2e9xB | 0.71 | 0.71 | 0.71 | 4hmoA | 0.77 | 0.77 | 0.77 | 4m00A | 0.54 | 0.54 | 0.54 |
| 2evrA | 0.71 | 0.75 | 0.75 | 4ie6A | 0.50 | 0.50 | 0.50 | 1bp1A | 0.66 | 0.84 | 0.84 |
| 2ew9A | 0.82 | 0.82 | 0.82 | 4l5gA | 0.80 | 0.80 | 0.80 | 1cjsA | 0.98 | 0.98 | 0.98 |
| 2fd5A | 0.78 | 0.78 | 0.78 | 4lpqA | 0.77 | 0.77 | 0.77 | 1ck1A | 0.80 | 0.87 | 0.90 |

| | | | | | | | | | | | |
|---|---|---|---|---|---|---|---|---|---|---|---|
| 2gh8A | 0.72 | 0.83 | 0.94 | 4m8rA | 0.65 | 0.65 | 0.65 | 1ecrA | 0.50 | 0.50 | 0.50 |
| 2gt1A | 0.80 | 0.80 | 0.80 | 4n06B | 0.82 | 0.82 | 0.82 | 1f5qD | 0.80 | 0.80 | 0.80 |
| 2gzaC | 0.72 | 0.82 | 0.82 | 4nj5A | 0.40 | 0.84 | 0.84 | 1fa9A | 0.52 | 0.94 | 0.95 |
| 2hjqA | 0.55 | 0.55 | 0.55 | 4opaB | 0.50 | 0.51 | 0.51 | 1gu7A | 0.77 | 0.86 | 0.86 |
| 2hwjA | 0.49 | 0.49 | 0.49 | 4qkuB | 0.54 | 0.54 | 0.88 | 1itwA | 0.47 | 0.47 | 0.47 |
| 2ijd1 | 0.48 | 0.49 | 0.49 | 4up9A | 0.70 | 0.89 | 0.90 | 1jkiA | 0.73 | 0.93 | 0.93 |
| 2iu7A | 0.50 | 0.81 | 0.98 | 4w7sA | 0.74 | 0.82 | 0.82 | 1n80A | 0.39 | 0.39 | 0.39 |
| 2iw2A | 0.79 | 0.79 | 0.79 | 1bf2A | 0.65 | 0.77 | 0.77 | 1nzjA | 0.71 | 0.85 | 0.85 |
| 2jz4A | 0.50 | 0.74 | 0.74 | 1bhgA | 0.77 | 0.87 | 0.87 | 1qhdA | 0.51 | 0.51 | 0.51 |
| 2kdyA | 0.51 | 0.51 | 0.71 | 1f7uA | 0.77 | 0.79 | 0.79 | 1qz9A | 0.90 | 0.90 | 0.90 |
| 2kn4A | 0.57 | 0.57 | 0.73 | 1fx7A | 0.74 | 0.74 | 0.91 | 1sb7B | 0.83 | 0.83 | 0.83 |
| 2mbgA | 0.53 | 0.53 | 0.53 | 1griA | 0.76 | 0.76 | 0.76 | 1vk1A | 0.73 | 0.73 | 0.73 |
| 2nsfA | 0.47 | 0.47 | 0.47 | 1h88C | 0.56 | 0.72 | 0.72 | 1vrmA | 0.84 | 0.87 | 0.87 |
| 2nykA | 0.72 | 0.72 | 0.72 | 1m8pB | 0.88 | 0.88 | 0.88 | 1xvuA | 0.76 | 0.76 | 0.76 |
| 2o6yA | 0.88 | 0.92 | 0.92 | 1ni5A | 0.76 | 0.76 | 0.76 | 1yy3A | 0.48 | 0.77 | 0.77 |
| 2owbA | 0.91 | 0.92 | 0.93 | 1q25A | 0.80 | 0.80 | 0.80 | 1z87A | 0.54 | 0.54 | 0.54 |
| 2qfiA | 0.48 | 0.71 | 0.71 | 1uzjA | 0.45 | 0.67 | 0.67 | 2a1sC | 0.49 | 0.50 | 0.50 |
| 2qp2A | 0.58 | 0.58 | 0.58 | 1zpuA | 0.86 | 0.87 | 0.87 | 2a3lA | 0.45 | 0.45 | 0.45 |
| 2qygA | 0.82 | 0.82 | 0.97 | 1zy9A | 0.69 | 0.69 | 0.69 | 2bt1A | 0.69 | 0.69 | 0.69 |
| 2r5wB | 0.50 | 0.90 | 0.90 | 2b5uA | 0.43 | 0.43 | 0.43 | 2bydA | 0.70 | 0.70 | 0.70 |
| 2uu7A | 0.72 | 0.81 | 0.89 | 2ewfA | 0.50 | 0.50 | 0.50 | 2c43A | 0.71 | 0.71 | 0.71 |
| 2w4bA | 0.42 | 0.86 | 0.86 | 2piaA | 0.75 | 0.90 | 0.90 | 2dfyC | 0.65 | 0.81 | 0.81 |
| 2x7iA | 0.76 | 0.77 | 0.77 | 2r7dA | 0.70 | 0.70 | 0.70 | 2dlaA | 0.64 | 0.64 | 0.64 |
| 2x8kC | 0.77 | 0.77 | 0.77 | 2uwnA | 0.64 | 0.64 | 0.64 | 2g3pA | 0.40 | 0.42 | 0.42 |
| 2yilA | 0.71 | 0.72 | 0.72 | 2v0nA | 0.50 | 0.52 | 0.52 | 2gg6A | 0.85 | 0.94 | 0.94 |
| 2yrqA | 0.64 | 0.64 | 0.64 | 2vgmA | 0.76 | 0.76 | 0.76 | 2gsyE | 0.48 | 0.86 | 0.86 |
| 2zxcA | 0.38 | 0.91 | 0.91 | 2wqrB | 0.82 | 0.82 | 0.82 | 2gzoA | 0.56 | 0.60 | 0.60 |
| 3a1iA | 0.58 | 0.59 | 0.59 | 2y25B | 0.71 | 0.71 | 0.71 | 2j2cA | 0.43 | 0.81 | 0.81 |
| 3a45A | 0.78 | 0.78 | 0.78 | 2yk0A | 0.49 | 0.53 | 0.53 | 2kfwA | 0.59 | 0.60 | 0.60 |
| 3a56A | 0.46 | 0.46 | 0.46 | 2zzqA | 0.51 | 0.51 | 0.51 | 2l9yA | 0.50 | 0.50 | 0.50 |
| 3ajvA | 0.67 | 0.70 | 0.70 | 3bt1U | 0.42 | 0.42 | 0.84 | 2ntyB | 0.49 | 0.49 | 0.49 |
| 3aqkA | 0.73 | 0.73 | 0.73 | 3c1yA | 0.47 | 0.47 | 0.47 | 2r3vA | 0.73 | 0.73 | 0.73 |
| 3arbA | 0.88 | 0.93 | 0.96 | 3cw2C | 0.82 | 0.83 | 0.83 | 2r58A | 0.48 | 0.85 | 0.98 |
| 3aujG | 0.46 | 0.47 | 0.89 | 3f83A | 0.55 | 0.69 | 0.69 | 2w4mA | 0.73 | 0.73 | 0.73 |
| 3b2zF | 0.78 | 0.86 | 0.89 | 3fc3A | 0.47 | 0.47 | 0.47 | 2x0cA | 0.86 | 0.86 | 0.86 |
| 3b7wA | 0.91 | 0.91 | 0.91 | 3gbgA | 0.49 | 0.49 | 0.49 | 2y51A | 0.82 | 0.92 | 0.92 |
| 3bt3A | 0.70 | 0.70 | 0.70 | 3h5cB | 0.73 | 0.73 | 0.73 | 2yb0E | 0.48 | 0.79 | 0.79 |
| 3c4tA | 0.77 | 0.77 | 0.77 | 3ibjA | 0.75 | 0.75 | 0.75 | 2z86C | 0.48 | 0.48 | 0.48 |
| 3craA | 0.55 | 0.55 | 0.55 | 3ippB | 0.63 | 0.87 | 0.87 | 3afoA | 0.72 | 0.74 | 0.74 |
| 3d30A | 0.80 | 0.84 | 0.84 | 3jymB | 0.65 | 0.65 | 0.65 | 3bu2A | 0.52 | 0.57 | 0.57 |
| 3eo5A | 0.45 | 0.51 | 0.51 | 3kbgA | 0.51 | 0.85 | 0.85 | 3cvzA | 0.50 | 0.50 | 0.50 |
| 3errA | 0.58 | 0.58 | 0.58 | 3mc8A | 0.72 | 0.72 | 0.84 | 3dupA | 0.49 | 0.50 | 0.50 |
| 3g79A | 0.83 | 0.85 | 0.85 | 3npfA | 0.73 | 0.73 | 0.73 | 3eswA | 0.47 | 0.75 | 0.75 |
| 3h2tA | 0.45 | 0.46 | 0.46 | 3orjA | 0.73 | 0.73 | 0.73 | 3eukH | 0.55 | 0.55 | 0.55 |

| | | | | | | | | | | | |
|---|---|---|---|---|---|---|---|---|---|---|---|
| 3hcsA | 0.45 | 0.50 | 0.80 | 3plaA | 0.49 | 0.88 | 0.88 | 3fi7A | 0.67 | 0.82 | 0.82 |
| 3hyiA | 0.52 | 0.52 | 0.52 | 3qe9Y | 0.49 | 0.49 | 0.49 | 3fvvA | 0.75 | 0.75 | 0.75 |
| 3i2dA | 0.79 | 0.83 | 0.83 | 3qjoA | 0.46 | 0.63 | 0.63 | 3gmsA | 0.88 | 0.88 | 0.88 |
| 3iam2 | 0.73 | 0.85 | 0.85 | 3qphA | 0.48 | 0.48 | 0.49 | 3hzzB | 0.73 | 0.84 | 0.93 |
| 3ifrA | 0.90 | 0.91 | 0.91 | 3qyeA | 0.66 | 0.67 | 0.90 | 3m1uA | 0.52 | 0.87 | 0.87 |
| 3isqA | 0.83 | 0.86 | 0.86 | 3rimA | 0.80 | 0.92 | 0.92 | 3mw8A | 0.80 | 0.88 | 0.88 |
| 3j7aK | 0.81 | 0.93 | 0.95 | 3rrpA | 0.69 | 0.88 | 0.88 | 3mwcA | 0.86 | 0.87 | 0.87 |
| 3k1rA | 0.55 | 0.55 | 0.55 | 3soaA | 0.54 | 0.54 | 0.54 | 3nsjA | 0.45 | 0.45 | 0.45 |
| 3k2iA | 0.55 | 0.55 | 0.95 | 3tixD | 0.43 | 0.44 | 0.44 | 3ntkA | 0.78 | 0.78 | 0.78 |
| 3kh5A | 0.81 | 0.81 | 0.81 | 3tp9A | 0.48 | 0.48 | 0.48 | 3oaaG | 0.76 | 0.82 | 0.82 |
| 3kjpA | 0.67 | 0.67 | 0.67 | 3ua3A | 0.46 | 0.82 | 0.82 | 3ptyA | 0.48 | 0.88 | 0.88 |
| 3kt1A | 0.69 | 0.69 | 0.69 | 3uj0A | 0.71 | 0.72 | 0.72 | 3rfyA | 0.46 | 0.47 | 0.47 |
| 3ktmE | 0.50 | 0.50 | 0.50 | 3vn4A | 0.42 | 0.42 | 0.42 | 3seoB | 0.49 | 0.49 | 0.49 |
| 3kzwA | 0.83 | 0.84 | 0.84 | 3vsmA | 0.69 | 0.69 | 0.69 | 3spgA | 0.75 | 0.80 | 0.94 |
| 3l76A | 0.58 | 0.60 | 0.60 | 3w1bA | 0.81 | 0.81 | 0.81 | 3u0kA | 0.55 | 0.84 | 0.90 |
| 3ld1A | 0.41 | 0.41 | 0.41 | 3zh9B | 0.75 | 0.75 | 0.75 | 3vlaA | 0.66 | 0.83 | 0.83 |
| 3lsgA | 0.84 | 0.90 | 0.90 | 4alzA | 0.69 | 0.69 | 0.69 | 3vstA | 0.54 | 0.54 | 0.54 |
| 3me4A | 0.75 | 0.79 | 0.79 | 4ax8A | 0.48 | 0.48 | 0.48 | 4aqfB | 0.42 | 0.81 | 0.91 |
| 3ml4C | 0.78 | 0.78 | 0.78 | 4b3iA | 0.84 | 0.88 | 0.88 | 4b21A | 0.80 | 0.87 | 0.87 |
| 3mx2B | 0.39 | 0.39 | 0.39 | 4bd9B | 0.47 | 0.53 | 0.53 | 4dt4A | 0.78 | 0.78 | 0.78 |
| 3mzfA | 0.82 | 0.88 | 0.95 | 4c0aB | 0.51 | 0.74 | 0.74 | 4dtfA | 0.44 | 0.44 | 0.44 |
| 3njaB | 0.74 | 0.74 | 0.74 | 4c0sA | 0.77 | 0.85 | 0.88 | 4ewtA | 0.83 | 0.92 | 0.92 |
| 3nqiA | 0.68 | 0.68 | 0.68 | 4dimA | 0.79 | 0.79 | 0.79 | 4f23A | 0.51 | 0.90 | 0.90 |
| 3nt8A | 0.41 | 0.41 | 0.41 | 4indA | 0.62 | 0.62 | 0.62 | 4fzbC | 0.79 | 0.79 | 0.79 |
| 3og5A | 0.82 | 0.88 | 0.88 | 4jdzB | 0.78 | 0.78 | 0.78 | 4g1pA | 0.84 | 0.94 | 0.96 |
| 3oh0A | 0.59 | 0.59 | 0.59 | 4kc3B | 0.83 | 0.83 | 0.83 | 4gfqA | 0.79 | 0.92 | 0.92 |
| 3pcsB | 0.66 | 0.66 | 0.66 | 4kikB | 0.79 | 0.79 | 0.90 | 4hvzA | 0.51 | 0.52 | 0.52 |
| 3po3S | 0.51 | 0.65 | 0.65 | 4lmfA | 0.54 | 0.88 | 0.88 | 4il6B | 0.44 | 0.44 | 0.44 |
| 3pxpA | 0.47 | 0.47 | 0.47 | 4lziA | 0.55 | 0.55 | 0.55 | 4jxkA | 0.83 | 0.93 | 0.93 |
| 3qavA | 0.83 | 0.83 | 0.83 | 4m9pA | 0.75 | 0.75 | 0.75 | 4m8mB | 0.39 | 0.80 | 0.80 |
| 3qf4B | 0.87 | 0.89 | 0.89 | 4pt5A | 0.82 | 0.87 | 0.87 | 4mzyA | 0.73 | 0.83 | 0.83 |
| 3qjjA | 0.82 | 0.86 | 0.86 | 4uwhA | 0.82 | 0.83 | 0.90 | 4onyA | 0.75 | 0.94 | 0.94 |
| 3qtdA | 0.92 | 0.92 | 0.92 | 1c1zA | 0.66 | 0.69 | 0.69 | 4pyhA | 0.70 | 0.70 | 0.70 |
| 3r6bA | 0.51 | 0.82 | 0.82 | 1d2pA | 0.61 | 0.61 | 0.61 | 4rg1A | 0.44 | 0.75 | 0.75 |
| 3rh7A | 0.47 | 0.47 | 0.47 | 1k7tA | 0.36 | 0.41 | 0.41 | | | | |

**Table S13.** The domain assembly results of SADA and SADA-w/o-D on 356 test proteins.

| PDB ID | TM-score | | PDB ID | TM-score | | PDB ID | TM-score | |
|---|---|---|---|---|---|---|---|---|
| | SADA | SADA-w/o-D | | SADA | SADA-w/o-D | | SADA | SADA-w/o-D |
| 1cjyA | 0.81 | 0.82 | 3rwxA | 0.60 | 0.59 | 1kfqA | 0.97 | 0.93 |
| 1efdN | 0.97 | 0.98 | 3sb4A | 0.61 | 0.55 | 1ldjA | 0.62 | 0.31 |
| 1fjrA | 0.86 | 0.68 | 3swjA | 0.71 | 0.73 | 1nyqB | 0.87 | 0.65 |
| 1g87B | 0.76 | 0.78 | 3t58B | 0.58 | 0.56 | 1ug9A | 0.65 | 0.47 |
| 1hx6B | 0.98 | 0.99 | 3t7jA | 0.97 | 0.95 | 1z1wA | 0.93 | 0.88 |
| 1iwaA | 0.99 | 0.99 | 3u07C | 0.88 | 0.80 | 2au3A | 0.81 | 0.69 |
| 1m5qH | 0.58 | 0.67 | 3u0oB | 0.99 | 0.98 | 2ii2A | 0.87 | 0.87 |
| 1mkfA | 0.99 | 0.99 | 3u9gA | 0.77 | 0.73 | 2olsA | 0.49 | 0.47 |
| 1mkmB | 0.70 | 0.70 | 3ub1D | 0.62 | 0.61 | 2ra1A | 0.49 | 0.37 |
| 1nh2D | 0.58 | 0.54 | 3uitD | 0.57 | 0.55 | 2v5dA | 0.69 | 0.80 |
| 1pprM | 0.57 | 0.61 | 3uo3A | 0.99 | 0.96 | 2xt6A | 0.54 | 0.51 |
| 1prrA | 0.60 | 0.72 | 3v7oB | 0.65 | 0.68 | 2zpaB | 0.90 | 0.90 |
| 1q19A | 0.99 | 0.99 | 3vr8B | 1.00 | 1.00 | 3apoA | 0.45 | 0.39 |
| 1qwrA | 1.00 | 0.99 | 3wkuA | 0.76 | 0.75 | 3b43A | 0.36 | 0.21 |
| 1r71B | 0.88 | 0.92 | 3zvmA | 0.59 | 0.57 | 3gf5B | 0.41 | 0.30 |
| 1rh1A | 0.88 | 0.59 | 4acoA | 0.91 | 0.93 | 3hjlA | 0.36 | 0.42 |
| 1rktA | 0.97 | 0.89 | 4ap5A | 0.99 | 0.98 | 3kq4B | 0.36 | 0.45 |
| 1s6lA | 0.76 | 0.73 | 4axdA | 0.99 | 0.66 | 3kwlA | 0.58 | 0.35 |
| 1sp3A | 1.00 | 0.93 | 4bfiB | 0.83 | 0.84 | 3ob8A | 0.97 | 0.37 |
| 1vz6A | 0.95 | 0.71 | 4bt9B | 0.61 | 0.61 | 3opfB | 0.68 | 0.40 |
| 1w3aA | 0.62 | 0.58 | 4cczA | 0.96 | 0.89 | 3p53A | 0.37 | 0.45 |
| 1wv3A | 0.93 | 0.92 | 4d0nB | 0.91 | 0.91 | 3pvlA | 0.68 | 0.38 |
| 1x7pA | 0.99 | 0.80 | 4d1iG | 0.96 | 0.80 | 3r05A | 0.32 | 0.28 |
| 1x9yA | 0.63 | 0.58 | 4dj3A | 0.98 | 0.98 | 3ubhA | 0.68 | 0.36 |
| 1y11A | 0.60 | 0.56 | 4dqaA | 0.64 | 0.62 | 3w2wA | 0.85 | 0.54 |
| 1yiqA | 0.88 | 0.88 | 4eo3A | 0.58 | 0.58 | 3zniA | 0.41 | 0.35 |
| 1zbuB | 0.79 | 0.77 | 4eogA | 0.98 | 0.99 | 4aimA | 0.91 | 0.52 |
| 1ze1A | 0.99 | 0.99 | 4etxA | 0.55 | 0.55 | 4ak1A | 0.29 | 0.29 |
| 2ablA | 0.70 | 0.68 | 4fguA | 0.92 | 0.70 | 4aq1A | 0.45 | 0.31 |
| 2ahvA | 0.82 | 0.80 | 4fkcA | 0.94 | 0.94 | 4fe9A | 0.47 | 0.48 |
| 2bkpA | 0.83 | 0.59 | 4fxkC | 0.97 | 0.97 | 4h2aA | 0.61 | 0.45 |
| 2c1yA | 0.56 | 0.58 | 4gbyA | 1.00 | 0.99 | 4i5sB | 0.55 | 0.44 |
| 2cxcA | 0.96 | 0.85 | 4ggmX | 0.61 | 0.60 | 4iggB | 0.31 | 0.31 |
| 2d1cA | 0.84 | 0.83 | 4gslA | 0.55 | 0.54 | 4j9vA | 0.87 | 0.58 |
| 2d7iA | 0.74 | 0.74 | 4gyjA | 1.00 | 0.99 | 4k3bA | 0.56 | 0.51 |
| 2e9hA | 0.97 | 0.73 | 4h3tA | 0.98 | 0.90 | 4kwuA | 0.78 | 0.51 |
| 2e9xB | 0.99 | 0.92 | 4hmoA | 0.99 | 0.93 | 4m00A | 0.52 | 0.52 |
| 2evrA | 0.99 | 0.99 | 4ie6A | 0.70 | 0.69 | 1bp1A | 0.99 | 0.97 |
| 2ew9A | 0.54 | 0.55 | 4l5gA | 0.97 | 0.68 | 1cjsA | 0.84 | 0.86 |
| 2fd5A | 0.99 | 0.99 | 4lpqA | 0.91 | 0.64 | 1ck1A | 0.99 | 0.99 |
| 2gh8A | 0.72 | 0.71 | 4m8rA | 0.77 | 0.77 | 1ecrA | 0.88 | 0.86 |

| | | | | | | | | |
|---|---|---|---|---|---|---|---|---|
| 2gt1A | 0.97 | 0.97 | 4n06B | 0.99 | 0.99 | 1f5qD | 0.99 | 0.99 |
| 2gzaC | 0.98 | 0.85 | 4nj5A | 0.69 | 0.61 | 1fa9A | 1.00 | 0.95 |
| 2hjqA | 0.55 | 0.53 | 4opaB | 0.71 | 0.82 | 1gu7A | 1.00 | 0.96 |
| 2hwjA | 0.83 | 0.69 | 4qkuB | 1.00 | 0.97 | 1itwA | 0.99 | 0.96 |
| 2ijd1 | 0.77 | 0.77 | 4up9A | 0.96 | 0.95 | 1jkiA | 1.00 | 0.99 |
| 2iu7A | 0.60 | 0.59 | 4w7sA | 0.64 | 0.64 | 1n80A | 1.00 | 0.84 |
| 2iw2A | 0.98 | 0.98 | 1bf2A | 0.99 | 0.99 | 1nzjA | 0.99 | 0.99 |
| 2jz4A | 0.59 | 0.57 | 1bhgA | 1.00 | 1.00 | 1qhdA | 0.90 | 0.93 |
| 2kdyA | 0.59 | 0.59 | 1f7uA | 0.97 | 0.94 | 1qz9A | 0.99 | 0.99 |
| 2kn4A | 0.65 | 0.61 | 1fx7A | 0.87 | 0.92 | 1sb7B | 0.87 | 0.89 |
| 2mbgA | 0.78 | 0.79 | 1griA | 0.68 | 0.61 | 1vk1A | 0.85 | 0.85 |
| 2nsfA | 0.94 | 0.67 | 1h88C | 0.71 | 0.45 | 1vrmA | 0.99 | 0.96 |
| 2nykA | 0.95 | 0.92 | 1m8pB | 0.68 | 0.41 | 1xvuA | 0.98 | 0.97 |
| 2o6yA | 0.99 | 0.96 | 1ni5A | 0.75 | 0.66 | 1yy3A | 0.70 | 0.70 |
| 2owbA | 0.98 | 0.97 | 1q25A | 0.49 | 0.47 | 1z87A | 0.67 | 0.55 |
| 2qfiA | 0.87 | 0.72 | 1uzjA | 0.78 | 0.52 | 2a1sC | 0.86 | 0.85 |
| 2qp2A | 0.74 | 0.74 | 1zpuA | 0.99 | 1.00 | 2a3lA | 0.97 | 0.99 |
| 2qygA | 0.97 | 0.98 | 1zy9A | 0.92 | 0.88 | 2bt1A | 0.99 | 0.90 |
| 2r5wB | 0.80 | 0.59 | 2b5uA | 0.48 | 0.47 | 2bydA | 0.98 | 0.66 |
| 2uu7A | 0.99 | 0.96 | 2ewfA | 0.49 | 0.57 | 2c43A | 0.93 | 0.68 |
| 2w4bA | 0.83 | 0.81 | 2piaA | 0.86 | 0.74 | 2dfyC | 0.66 | 0.54 |
| 2x7iA | 0.98 | 1.00 | 2r7dA | 0.98 | 0.93 | 2dlaA | 1.00 | 0.97 |
| 2x8kC | 0.98 | 0.96 | 2uwnA | 0.77 | 0.42 | 2g3pA | 0.63 | 0.60 |
| 2yilA | 0.63 | 0.64 | 2v0nA | 0.45 | 0.53 | 2gg6A | 0.99 | 1.00 |
| 2yrqA | 0.52 | 0.52 | 2vgmA | 0.76 | 0.60 | 2gsyE | 0.92 | 0.70 |
| 2zxcA | 0.82 | 0.82 | 2wqrB | 0.49 | 0.62 | 2gzoA | 0.93 | 0.86 |
| 3a1iA | 0.93 | 0.93 | 2y25B | 0.57 | 0.45 | 2j2cA | 0.96 | 0.77 |
| 3a45A | 0.99 | 0.98 | 2yk0A | 0.72 | 0.74 | 2kfwA | 0.78 | 0.77 |
| 3a56A | 0.66 | 0.66 | 2zzqA | 0.46 | 0.44 | 2l9yA | 0.69 | 0.69 |
| 3ajvA | 0.98 | 0.85 | 3bt1U | 0.61 | 0.49 | 2ntyB | 0.82 | 0.76 |
| 3aqkA | 0.96 | 0.98 | 3c1yA | 0.63 | 0.61 | 2r3vA | 1.00 | 0.96 |
| 3arbA | 0.94 | 0.94 | 3cw2C | 0.70 | 0.70 | 2r58A | 0.65 | 0.55 |
| 3aujG | 0.99 | 0.70 | 3f83A | 0.46 | 0.39 | 2w4mA | 0.89 | 0.85 |
| 3b2zF | 0.76 | 0.76 | 3fc3A | 0.54 | 0.49 | 2x0cA | 0.66 | 0.66 |
| 3b7wA | 0.83 | 0.83 | 3gbgA | 0.64 | 0.60 | 2y51A | 0.99 | 0.99 |
| 3bt3A | 0.95 | 0.96 | 3h5cB | 0.97 | 0.98 | 2yb0E | 0.97 | 0.79 |
| 3c4tA | 0.74 | 0.74 | 3ibjA | 0.58 | 0.50 | 2z86C | 0.69 | 0.68 |
| 3craA | 0.61 | 0.57 | 3ippB | 0.74 | 0.44 | 3afoA | 0.67 | 0.68 |
| 3d30A | 0.96 | 0.94 | 3jymB | 0.43 | 0.43 | 3bu2A | 0.91 | 0.91 |
| 3eo5A | 0.64 | 0.59 | 3kbgA | 0.62 | 0.49 | 3cvzA | 0.87 | 0.76 |
| 3errA | 0.79 | 0.80 | 3mc8A | 0.83 | 0.43 | 3dupA | 0.98 | 0.87 |
| 3g79A | 0.90 | 0.90 | 3npfA | 0.93 | 0.92 | 3eswA | 0.89 | 0.88 |
| 3h2tA | 0.81 | 0.81 | 3orjA | 0.66 | 0.74 | 3eukH | 0.92 | 0.90 |
| 3hcsA | 0.80 | 0.67 | 3plaA | 0.65 | 0.53 | 3fi7A | 0.99 | 0.93 |

| | | | | | | | | |
|---|---|---|---|---|---|---|---|---|
| 3hyiA | 0.71 | 0.69 | 3qe9Y | 0.94 | 0.72 | 3fvvA | 0.98 | 0.96 |
| 3i2dA | 0.86 | 0.84 | 3qjoA | 0.80 | 0.67 | 3gmsA | 1.00 | 0.99 |
| 3iam2 | 0.92 | 0.93 | 3qphA | 0.48 | 0.47 | 3hzzB | 1.00 | 0.99 |
| 3ifrA | 0.99 | 0.98 | 3qyeA | 0.99 | 0.95 | 3m1uA | 0.98 | 0.99 |
| 3isqA | 1.00 | 0.99 | 3rimA | 0.98 | 0.98 | 3mw8A | 0.75 | 0.64 |
| 3j7aK | 0.96 | 0.98 | 3rrpA | 0.98 | 0.95 | 3mwcA | 1.00 | 0.99 |
| 3k1rA | 0.66 | 0.60 | 3soaA | 0.73 | 0.56 | 3nsjA | 0.82 | 0.82 |
| 3k2iA | 0.78 | 0.75 | 3tixD | 0.63 | 0.60 | 3ntkA | 0.98 | 0.92 |
| 3kh5A | 0.94 | 0.96 | 3tp9A | 0.87 | 0.57 | 3oaaG | 0.98 | 0.98 |
| 3kjpA | 0.91 | 0.58 | 3ua3A | 0.58 | 0.50 | 3ptyA | 0.75 | 0.71 |
| 3kt1A | 0.91 | 0.90 | 3uj0A | 0.94 | 0.91 | 3rfyA | 0.95 | 0.90 |
| 3ktmE | 0.63 | 0.64 | 3vn4A | 0.47 | 0.43 | 3seoB | 0.77 | 0.81 |
| 3kzwA | 1.00 | 0.97 | 3vsmA | 0.80 | 0.87 | 3spgA | 0.98 | 0.98 |
| 3l76A | 0.89 | 0.89 | 3w1bA | 0.72 | 0.73 | 3u0kA | 0.61 | 0.62 |
| 3ld1A | 0.70 | 0.69 | 3zh9B | 0.62 | 0.59 | 3vlaA | 1.00 | 0.98 |
| 3lsgA | 0.86 | 0.85 | 4alzA | 0.69 | 0.69 | 3vstA | 1.00 | 0.98 |
| 3me4A | 0.90 | 0.77 | 4ax8A | 0.64 | 0.52 | 4aqfB | 0.88 | 0.87 |
| 3ml4C | 0.65 | 0.63 | 4b3iA | 0.94 | 0.95 | 4b21A | 0.99 | 0.90 |
| 3mx2B | 0.92 | 0.70 | 4bd9B | 0.61 | 0.40 | 4dt4A | 0.98 | 0.93 |
| 3mzfA | 0.93 | 0.86 | 4c0aB | 0.57 | 0.48 | 4dtfA | 0.95 | 0.85 |
| 3njaB | 0.61 | 0.61 | 4c0sA | 0.63 | 0.61 | 4ewtA | 0.84 | 0.87 |
| 3nqiA | 0.94 | 0.95 | 4dimA | 0.89 | 0.86 | 4f23A | 1.00 | 0.97 |
| 3nt8A | 0.76 | 0.57 | 4indA | 0.46 | 0.45 | 4fzbC | 0.99 | 0.99 |
| 3og5A | 0.97 | 0.89 | 4jdzB | 0.70 | 0.67 | 4g1pA | 0.93 | 0.93 |
| 3oh0A | 0.97 | 0.79 | 4kc3B | 0.76 | 0.75 | 4gfqA | 0.76 | 0.70 |
| 3pcsB | 0.86 | 0.80 | 4kikB | 0.73 | 0.72 | 4hvzA | 0.75 | 0.57 |
| 3po3S | 0.96 | 0.56 | 4lmfA | 0.49 | 0.45 | 4il6B | 0.73 | 0.72 |
| 3pxpA | 0.74 | 0.68 | 4lziA | 0.67 | 0.43 | 4jxkA | 0.98 | 0.96 |
| 3qavA | 0.98 | 0.93 | 4m9pA | 0.51 | 0.49 | 4m8mB | 0.87 | 0.85 |
| 3qf4B | 0.93 | 0.93 | 4pt5A | 0.91 | 0.88 | 4mzyA | 1.00 | 0.99 |
| 3qjjA | 0.97 | 0.98 | 4uwhA | 0.99 | 1.00 | 4onyA | 0.81 | 0.80 |
| 3qtdA | 1.00 | 1.00 | 1c1zA | 0.46 | 0.28 | 4pyhA | 0.77 | 0.75 |
| 3r6bA | 0.54 | 0.52 | 1d2pA | 0.36 | 0.47 | 4rg1A | 0.77 | 0.66 |
| 3rh7A | 0.71 | 0.58 | 1k7tA | 0.67 | 0.32 | | | |

**Table S14.** Summary of the domain assembly results obtained by SADA and SADA-w/o-D.

| Method | 2dom (166) | 3dom (69) | m4dom (40) | 2dis (81) | Total (356) |
|---|---|---|---|---|---|
| SADA | 0.83 | 0.72 | 0.60 | 0.89 | 0.80 |
| SADA-w/o-D | 0.79 | 0.65 | 0.48 | 0.85 | 0.74 |

**Table S15.** Under the sequence identity cutoff of 30%, 50% and 70%, the TM-score between the native structure of test proteins and top 1 homologous templates of the test protein detected by JackHMMER in MPDB. "-" represents the homologous template of the test protein cannot be detected by JackHMMER with default parameters in MPDB.

| PDB ID | TM-score | | | PDB ID | TM-score | | | PDB ID | TM-score | | |
|---|---|---|---|---|---|---|---|---|---|---|---|
| | 30% | 50% | 70% | | 30% | 50% | 70% | | 30% | 50% | 70% |
| 1cjyA | 0.25 | 0.81 | 0.81 | 3rwxA | - | - | - | 1kfqA | 0.77 | 0.87 | 0.93 |
| 1efdN | 0.79 | 0.79 | 0.79 | 3sb4A | 0.41 | 0.41 | 0.41 | 1ldjA | 0.43 | 0.80 | 0.80 |
| 1fjrA | - | - | - | 3swjA | 0.55 | 0.55 | 0.91 | 1nyqB | 0.59 | 0.87 | 0.87 |
| 1g87B | 0.61 | 0.94 | 0.94 | 3t58B | 0.48 | 0.48 | 0.48 | 1ug9A | 0.63 | 0.63 | 0.63 |
| 1hx6B | - | - | - | 3t7jA | 0.69 | 0.69 | 0.69 | 1z1wA | 0.87 | 0.87 | 0.87 |
| 1iwaA | 0.86 | 0.93 | 0.98 | 3u07C | - | - | - | 2au3A | 0.66 | 0.66 | 0.66 |
| 1m5qH | - | - | - | 3u0oB | 0.80 | 0.89 | 0.89 | 2ii2A | 0.80 | 0.80 | 0.80 |
| 1mkfA | - | - | - | 3u9gA | - | - | - | 2olsA | 0.40 | 0.40 | 0.40 |
| 1mkmB | 0.73 | 0.73 | 0.73 | 3ub1D | - | - | - | 2ra1A | - | - | - |
| 1nh2D | - | 0.88 | 0.88 | 3uitD | - | 0.48 | 0.48 | 2v5dA | 0.71 | 0.72 | 0.72 |
| 1pprM | - | 0.90 | 0.90 | 3uo3A | 0.76 | 0.76 | 0.76 | 2xt6A | - | 0.75 | 0.75 |
| 1prrA | 0.41 | 0.41 | 0.41 | 3v7oB | 0.33 | 0.33 | 0.33 | 2zpaB | 0.82 | 0.82 | 0.82 |
| 1q19A | 0.84 | 0.84 | 0.84 | 3vr8B | 0.87 | 0.89 | 0.94 | 3apoA | 0.21 | 0.21 | 0.21 |
| 1qwrA | 0.82 | 0.82 | 0.82 | 3wkuA | - | - | - | 3b43A | 0.20 | 0.20 | 0.20 |
| 1r71B | 0.71 | 0.71 | 0.71 | 3zvmA | 0.33 | 0.33 | 0.33 | 3gf5B | 0.18 | 0.18 | 0.18 |
| 1rh1A | 0.48 | 0.48 | 0.48 | 4acoA | 0.72 | 0.72 | 0.72 | 3hjlA | 0.34 | 0.34 | 0.34 |
| 1rktA | 0.63 | 0.63 | 0.63 | 4ap5A | 0.69 | 0.93 | 0.93 | 3kq4B | 0.28 | 0.28 | 0.28 |
| 1s6lA | - | - | - | 4axdA | 0.64 | 0.77 | 0.77 | 3kwlA | 0.49 | 0.49 | 0.49 |
| 1sp3A | - | - | - | 4bfiB | 0.51 | 0.51 | 0.51 | 3ob8A | 0.49 | 0.88 | 0.88 |
| 1vz6A | 0.50 | 0.50 | 0.52 | 4bt9B | 0.49 | 0.49 | 0.49 | 3opfB | 0.53 | 0.55 | 0.55 |
| 1w3aA | 0.52 | 0.52 | 0.52 | 4cczA | 0.73 | 0.73 | 0.73 | 3p53A | - | - | - |
| 1wv3A | - | - | - | 4d0nB | 0.77 | 0.77 | 0.77 | 3pvlA | 0.57 | 0.84 | 0.84 |
| 1x7pA | 0.82 | 0.79 | 0.79 | 4d1iG | 0.62 | 0.62 | 0.90 | 3r05A | 0.23 | 0.25 | 0.25 |
| 1x9yA | - | - | - | 4dj3A | 0.78 | 0.78 | 0.85 | 3ubhA | 0.47 | 0.47 | 0.47 |
| 1y11A | - | 0.48 | 0.48 | 4dqaA | - | - | - | 3w2wA | 0.62 | 0.62 | 0.62 |
| 1yiqA | 0.42 | 0.92 | 0.94 | 4eo3A | 0.45 | 0.45 | 0.45 | 3zniA | - | 0.71 | 0.76 |
| 1zbuB | - | - | - | 4eogA | 0.77 | 0.77 | 0.77 | 4aimA | 0.31 | 0.75 | 0.82 |
| 1ze1A | 0.77 | 0.74 | 0.74 | 4etxA | - | - | - | 4ak1A | - | 0.46 | 0.46 |
| 2ablA | 0.47 | 0.58 | 0.58 | 4fguA | - | 0.90 | 0.90 | 4aq1A | - | - | - |
| 2ahvA | 0.75 | 0.75 | 0.75 | 4fkcA | 0.88 | 0.89 | 0.89 | 4fe9A | 0.42 | 0.42 | 0.42 |
| 2bkpA | 0.44 | 0.46 | 0.46 | 4fxkC | 0.54 | 0.59 | 0.59 | 4h2aA | 0.65 | 0.71 | 0.71 |
| 2c1yA | - | 0.57 | 0.57 | 4gbyA | 0.78 | 0.86 | 0.86 | 4i5sB | 0.60 | 0.60 | 0.60 |
| 2cxcA | 0.74 | 0.74 | 0.74 | 4ggmX | - | - | - | 4iggB | 0.65 | 0.65 | 0.65 |
| 2d1cA | 0.66 | 0.66 | 0.66 | 4gslA | 0.41 | 0.47 | 0.47 | 4j9vA | 0.61 | 0.61 | 0.61 |
| 2d7iA | 0.42 | 0.64 | 0.64 | 4gyjA | 0.79 | 0.79 | 0.79 | 4k3bA | 0.46 | 0.54 | 0.54 |
| 2e9hA | 0.65 | 0.65 | 0.65 | 4h3tA | 0.76 | 0.90 | 0.90 | 4kwuA | 0.40 | 0.40 | 0.40 |
| 2e9xB | - | - | - | 4hmoA | 0.66 | 0.66 | 0.66 | 4m00A | - | 0.33 | 0.63 |
| 2evrA | 0.80 | 0.50 | 0.50 | 4ie6A | - | - | - | 1bp1A | 0.72 | 0.74 | 0.74 |

| | | | | | | | | | | | |
|---|---|---|---|---|---|---|---|---|---|---|---|
| 2ew9A | 0.54 | 0.43 | 0.43 | 4l5gA | 0.58 | 0.58 | 0.58 | 1cjsA | 0.70 | 0.81 | 0.81 |
| 2fd5A | 0.67 | 0.67 | 0.67 | 4lpqA | 0.62 | 0.62 | 0.62 | 1ck1A | 0.79 | 0.89 | 0.92 |
| 2gh8A | 0.60 | 0.77 | 0.94 | 4m8rA | - | - | - | 1ecrA | - | - | - |
| 2gt1A | 0.84 | 0.84 | 0.84 | 4n06B | 0.81 | 0.81 | 0.81 | 1f5qD | 0.78 | 0.78 | 0.78 |
| 2gzaC | 0.47 | 0.51 | 0.51 | 4nj5A | 0.27 | 0.86 | 0.86 | 1fa9A | - | 0.92 | 0.92 |
| 2hjqA | - | - | - | 4opaB | - | - | - | 1gu7A | 0.70 | 0.70 | 0.70 |
| 2hwjA | - | - | - | 4qkuB | - | - | 0.94 | 1itwA | - | - | - |
| 2ijd1 | 0.67 | 0.71 | 0.71 | 4up9A | 0.73 | 0.88 | 0.87 | 1jkiA | 0.70 | 0.94 | 0.94 |
| 2iu7A | - | 0.84 | 0.97 | 4w7sA | 0.52 | 0.52 | 0.52 | 1n80A | - | - | - |
| 2iw2A | 0.80 | 0.82 | 0.82 | 1bf2A | 0.74 | 0.85 | 0.85 | 1nzjA | 0.75 | 0.89 | 0.89 |
| 2jz4A | - | 0.47 | 0.47 | 1bhgA | 0.83 | 0.92 | 0.92 | 1qhdA | 0.14 | 0.14 | 0.14 |
| 2kdyA | - | - | 0.64 | 1f7uA | 0.85 | 0.84 | 0.84 | 1qz9A | 0.82 | 0.82 | 0.82 |
| 2kn4A | 0.46 | 0.44 | 0.46 | 1fx7A | 0.76 | 0.76 | 0.89 | 1sb7B | 0.78 | 0.82 | 0.82 |
| 2mbgA | 0.62 | 0.62 | 0.62 | 1griA | 0.48 | 0.48 | 0.48 | 1vk1A | 0.36 | 0.36 | 0.36 |
| 2nsfA | - | - | - | 1h88C | 0.37 | 0.64 | 0.66 | 1vrmA | 0.86 | 0.87 | 0.87 |
| 2nykA | - | - | - | 1m8pB | 0.62 | 0.62 | 0.62 | 1xvuA | 0.86 | 0.86 | 0.86 |
| 2o6yA | 0.86 | 0.95 | 0.95 | 1ni5A | 0.61 | 0.61 | 0.61 | 1yy3A | - | 0.78 | 0.78 |
| 2owbA | 0.85 | 0.80 | 0.80 | 1q25A | 0.44 | 0.44 | 0.44 | 1z87A | 0.33 | 0.38 | 0.38 |
| 2qfiA | - | 0.66 | 0.66 | 1uzjA | 0.33 | 0.32 | 0.32 | 2a1sC | - | - | - |
| 2qp2A | 0.65 | 0.65 | 0.65 | 1zpuA | 0.87 | 0.87 | 0.87 | 2a3lA | - | - | - |
| 2qygA | 0.89 | 0.89 | 0.99 | 1zy9A | 0.77 | 0.77 | 0.77 | 2bt1A | - | - | - |
| 2r5wB | 0.38 | 0.91 | 0.91 | 2b5uA | 0.47 | 0.47 | 0.47 | 2bydA | 0.72 | 0.72 | 0.72 |
| 2uu7A | 0.78 | 0.88 | 0.92 | 2ewfA | - | - | - | 2c43A | 0.67 | 0.67 | 0.67 |
| 2w4bA | - | 0.87 | 0.87 | 2piaA | 0.60 | 0.69 | 0.69 | 2dfyC | 0.29 | 0.29 | 0.29 |
| 2x7iA | 0.74 | 0.77 | 0.77 | 2r7dA | 0.83 | 0.83 | 0.83 | 2dlaA | - | - | - |
| 2x8kC | 0.82 | 0.82 | 0.82 | 2uwnA | 0.45 | 0.49 | 0.49 | 2g3pA | - | 0.41 | 0.41 |
| 2yilA | - | 0.56 | 0.56 | 2v0nA | 0.43 | 0.43 | 0.43 | 2gg6A | 0.79 | 0.79 | 0.79 |
| 2yrqA | 0.38 | 0.38 | 0.52 | 2vgmA | 0.66 | 0.57 | 0.57 | 2gsyE | - | 0.90 | 0.90 |
| 2zxcA | - | 0.93 | 0.93 | 2wqrB | 0.41 | 0.53 | 0.53 | 2gzoA | 0.61 | 0.65 | 0.65 |
| 3a1iA | 0.75 | 0.73 | 0.73 | 2y25B | 0.30 | 0.30 | 0.30 | 2j2cA | - | 0.82 | 0.82 |
| 3a45A | 0.80 | 0.80 | 0.80 | 2yk0A | 0.38 | 0.37 | 0.60 | 2kfwA | 0.39 | 0.45 | 0.45 |
| 3a56A | - | - | - | 2zzqA | - | - | - | 2l9yA | 0.34 | 0.34 | 0.34 |
| 3ajvA | - | - | - | 3bt1U | - | - | 0.86 | 2ntyB | - | - | - |
| 3aqkA | 0.72 | 0.72 | 0.72 | 3c1yA | 0.33 | 0.33 | 0.33 | 2r3vA | 0.69 | 0.69 | 0.69 |
| 3arbA | 0.71 | 0.71 | 0.71 | 3cw2C | 0.56 | 0.65 | 0.65 | 2r58A | - | 0.70 | 0.70 |
| 3aujG | - | - | 0.95 | 3f83A | - | 0.55 | 0.55 | 2w4mA | 0.75 | 0.75 | 0.75 |
| 3b2zF | 0.63 | 0.63 | 0.92 | 3fc3A | - | - | - | 2x0cA | 0.59 | 0.59 | 0.59 |
| 3b7wA | 0.72 | 0.70 | 0.70 | 3gbgA | 0.36 | 0.36 | 0.36 | 2y51A | 0.83 | 0.83 | 0.83 |
| 3bt3A | - | - | - | 3h5cB | 0.70 | 0.70 | 0.70 | 2yb0E | - | 0.81 | 0.74 |
| 3c4tA | 0.56 | 0.60 | 0.60 | 3ibjA | 0.47 | 0.47 | 0.47 | 2z86C | 0.37 | 0.37 | 0.37 |
| 3craA | 0.35 | 0.35 | 0.35 | 3ippB | 0.60 | 0.92 | 0.92 | 3afoA | 0.71 | 0.74 | 0.74 |
| 3d30A | - | 0.84 | 0.84 | 3jymB | 0.24 | 0.24 | 0.24 | 3bu2A | 0.50 | 0.51 | 0.51 |
| 3eo5A | - | - | - | 3kbgA | - | 0.85 | 0.85 | 3cvzA | - | - | - |
| 3errA | 0.72 | 0.72 | 0.69 | 3mc8A | 0.52 | 0.47 | 0.47 | 3dupA | - | - | - |

| | | | | | | | | | | | |
|---|---|---|---|---|---|---|---|---|---|---|---|
| 3g79A | 0.73 | 0.73 | 0.73 | 3npfA | 0.75 | 0.75 | 0.75 | 3eswA | 0.54 | 0.77 | 0.77 |
| 3h2tA | - | - | - | 3orjA | 0.57 | 0.57 | 0.57 | 3eukH | - | - | - |
| 3hcsA | 0.28 | 0.28 | 0.28 | 3plaA | 0.57 | 0.60 | 0.60 | 3fi7A | 0.66 | 0.77 | 0.77 |
| 3hyiA | - | - | - | 3qe9Y | 0.75 | 0.75 | 0.75 | 3fvvA | 0.69 | 0.69 | 0.69 |
| 3i2dA | 0.68 | 0.77 | 0.77 | 3qjoA | 0.59 | 0.84 | 0.84 | 3gmsA | 0.82 | 0.82 | 0.82 |
| 3iam2 | 0.74 | 0.89 | 0.89 | 3qphA | 0.35 | 0.35 | 0.67 | 3hzzB | 0.71 | 0.84 | 0.93 |
| 3ifrA | 0.82 | 0.82 | 0.82 | 3qyeA | 0.76 | 0.82 | 0.92 | 3m1uA | 0.90 | 0.90 | 0.90 |
| 3isqA | 0.86 | 0.88 | 0.88 | 3rimA | 0.82 | 0.92 | 0.92 | 3mw8A | 0.82 | 0.67 | 0.67 |
| 3j7aK | 0.86 | 0.86 | 0.98 | 3rrpA | 0.77 | 0.92 | 0.92 | 3mwcA | 0.88 | 0.89 | 0.89 |
| 3k1rA | 0.45 | 0.47 | 0.47 | 3soaA | 0.53 | 0.66 | 0.67 | 3nsjA | 0.56 | 0.56 | 0.56 |
| 3k2iA | - | - | 0.96 | 3tixD | - | - | - | 3ntkA | 0.80 | 0.80 | 0.80 |
| 3kh5A | 0.82 | 0.82 | 0.82 | 3tp9A | 0.47 | 0.70 | 0.70 | 3oaaG | 0.80 | 0.80 | 0.80 |
| 3kjpA | 0.43 | 0.43 | 0.43 | 3ua3A | 0.41 | 0.48 | 0.48 | 3ptyA | - | 0.91 | 0.91 |
| 3kt1A | 0.69 | 0.69 | 0.69 | 3uj0A | 0.71 | 0.43 | 0.52 | 3rfyA | - | - | - |
| 3ktmE | - | - | - | 3vn4A | 0.17 | 0.17 | 0.17 | 3seoB | - | - | - |
| 3kzwA | 0.89 | 0.89 | 0.89 | 3vsmA | - | - | - | 3spgA | 0.72 | 0.80 | 0.96 |
| 3l76A | 0.58 | 0.60 | 0.60 | 3w1bA | 0.65 | 0.65 | 0.65 | 3u0kA | 0.51 | 0.51 | 0.51 |
| 3ld1A | - | - | - | 3zh9B | - | - | - | 3vlaA | 0.79 | 0.87 | 0.87 |
| 3lsgA | 0.88 | 0.94 | 0.94 | 4alzA | - | - | - | 3vstA | - | - | - |
| 3me4A | 0.60 | 0.48 | 0.48 | 4ax8A | 0.37 | 0.37 | 0.37 | 4aqfB | - | 0.74 | 0.90 |
| 3ml4C | - | - | - | 4b3iA | 0.84 | 0.88 | 0.88 | 4b21A | 0.84 | 0.90 | 0.90 |
| 3mx2B | - | - | - | 4bd9B | 0.42 | 0.42 | 0.42 | 4dt4A | 0.53 | 0.68 | 0.77 |
| 3mzfA | 0.77 | 0.82 | 0.92 | 4c0aB | 0.40 | 0.40 | 0.40 | 4dtfA | - | - | - |
| 3njaB | 0.45 | 0.45 | 0.45 | 4c0sA | 0.70 | 0.81 | 0.53 | 4ewtA | 0.77 | 0.77 | 0.77 |
| 3nqiA | - | - | - | 4dimA | 0.77 | 0.67 | 0.67 | 4f23A | 0.59 | 0.87 | 0.87 |
| 3nt8A | 0.42 | 0.42 | 0.42 | 4indA | - | - | - | 4fzbC | 0.83 | 0.83 | 0.83 |
| 3og5A | 0.78 | 0.83 | 0.83 | 4jdzB | 0.59 | 0.59 | 0.59 | 4g1pA | 0.72 | 0.93 | 0.97 |
| 3oh0A | 0.55 | 0.55 | 0.55 | 4kc3B | 0.34 | 0.34 | 0.34 | 4gfqA | 0.63 | 0.68 | 0.68 |
| 3pcsB | - | - | - | 4kikB | 0.18 | 0.18 | 0.86 | 4hvzA | - | - | - |
| 3po3S | 0.30 | 0.72 | 0.72 | 4lmfA | 0.41 | 0.42 | 0.42 | 4il6B | 0.63 | 0.63 | 0.63 |
| 3pxpA | - | - | - | 4lziA | 0.42 | 0.42 | 0.42 | 4jxkA | 0.80 | 0.80 | 0.80 |
| 3qavA | 0.69 | 0.69 | 0.69 | 4m9pA | 0.34 | 0.34 | 0.34 | 4m8mB | 0.10 | 0.82 | 0.82 |
| 3qf4B | 0.76 | 0.85 | 0.85 | 4pt5A | 0.77 | 0.86 | 0.86 | 4mzyA | 0.36 | 0.83 | 0.83 |
| 3qjjA | 0.84 | 0.88 | 0.88 | 4uwhA | 0.84 | 0.84 | 0.84 | 4onyA | 0.63 | 0.93 | 0.93 |
| 3qtdA | 0.94 | 0.94 | 0.94 | 1c1zA | 0.51 | 0.44 | 0.44 | 4pyhA | 0.51 | 0.51 | 0.51 |
| 3r6bA | 0.38 | 0.69 | 0.76 | 1d2pA | - | - | 0.50 | 4rg1A | 0.80 | 0.80 | 0.80 |
| 3rh7A | - | 0.59 | 0.59 | 1k7tA | 0.33 | 0.39 | 0.39 | | | | |

**Table S16.** On the 256 test proteins that JackHMMER can detect homologues under the sequence identity cutoff of 30%, the summary of the quality of analogues and homologues. TM-score represents the average TM-score in the detected templates. #TM-score⩾0.5 is the number of analogue (homologue) with TM-score⩾0.5.

| Domain | Template | TM-score | TM-score⩾0.5 |
|---|---|---|---|
| 2dom | Analogue | 0.65 | 78 |
| (*N* = 110) | Homologue | 0.65 | 82 |
| 3dom | Analogue | 0.58 | 33 |
| (*N* = 58) | Homologue | 0.56 | 32 |
| m4dom | Analogue | 0.47 | 12 |
| (*N* = 32) | Homologue | 0.51 | 16 |
| 2dis | Analogue | 0.67 | 45 |
| (*N* = 56) | Homologue | 0.65 | 47 |
| All | Analogue | 0.62 | 168 |
| (*N* = 256) | Homologue | 0.61 | 177 |

**Table S17.** On the 285 test proteins that JackHMMER can detect homologues under the sequence identity cutoff of 50%, the summary of the quality of analogues and homologues. TM-score represents the average TM-score in the detected templates. #TM-score⩾0.5 is the number of analogue (homologue) with TM-score⩾0.5.

| Domain | Template | TM-score | TM-score⩾0.5 |
|---|---|---|---|
| 2dom | Analogue | 0.71 | 98 |
| (*N* = 124) | Homologue | 0.69 | 100 |
| 3dom | Analogue | 0.63 | 41 |
| (*N* = 60) | Homologue | 0.59 | 35 |
| m4dom | Analogue | 0.56 | 20 |
| (*N* = 36) | Homologue | 0.57 | 21 |
| 2dis | Analogue | 0.75 | 58 |
| (*N* = 65) | Homologue | 0.72 | 57 |
| All | Analogue | 0.68 | 217 |
| (*N* = 285) | Homologue | 0.66 | 213 |

**Table S18.** On the 291 test proteins that JackHMMER can detect homologues under the sequence identity cutoff of 70%, the summary of the quality of analogues and homologues.

| Domain | Template | TM-score | TM-score⩾0.5 |
|---|---|---|---|
| 2dom | Analogue | 0.73 | 104 |
| ($N$ = 128) | Homologue | 0.72 | 106 |
| 3dom | Analogue | 0.64 | 44 |
| ($N$ = 61) | Homologue | 0.62 | 40 |
| m4dom | Analogue | 0.56 | 21 |
| ($N$ = 37) | Homologue | 0.58 | 22 |
| 2dis | Analogue | 0.76 | 58 |
| ($N$ = 65) | Homologue | 0.73 | 57 |
| All | Analogue | 0.70 | 227 |
| ($N$ = 291) | Homologue | 0.68 | 225 |

**Table S19.** On 356 test proteins, the TM-score of the results assembled by SADA-w/o-D and SADA-w/o-D using homologue, where SADA-w/o-D-H represents the full-chain models generated by SADA-w/o-D using homologue and "-" represents the protein cannot be detected homologue by JackHMMER.

| PDB ID | TM-score | | PDB ID | TM-score | | PDB ID | TM-score | |
|---|---|---|---|---|---|---|---|---|
| | SADA-w/o-D-H | SADA-w/o-D | | SADA-w/o-D-H | SADA-w/o-D | | SADA-w/o-D-H | SADA-w/o-D |
| 1cjyA | 0.81 | 0.82 | 3rwxA | - | 0.59 | 1kfqA | 0.93 | 0.93 |
| 1efdN | 0.98 | 0.98 | 3sb4A | 0.60 | 0.55 | 1ldjA | 0.27 | 0.31 |
| 1fjrA | - | 0.68 | 3swjA | 0.72 | 0.73 | 1nyqB | 0.62 | 0.65 |
| 1g87B | 0.75 | 0.78 | 3t58B | 0.56 | 0.56 | 1ug9A | 0.43 | 0.47 |
| 1hx6B | - | 0.99 | 3t7jA | 0.95 | 0.95 | 1z1wA | 0.90 | 0.88 |
| 1iwaA | 0.99 | 0.99 | 3u07C | - | 0.80 | 2au3A | 0.39 | 0.69 |
| 1m5qH | - | 0.67 | 3u0oB | 0.98 | 0.98 | 2ii2A | 0.46 | 0.87 |
| 1mkfA | - | 0.99 | 3u9gA | - | 0.73 | 2olsA | 0.46 | 0.47 |
| 1mkmB | 0.71 | 0.70 | 3ub1D | - | 0.61 | 2ra1A | - | 0.37 |
| 1nh2D | - | 0.54 | 3uitD | - | 0.55 | 2v5dA | 0.69 | 0.80 |
| 1pprM | - | 0.61 | 3uo3A | 0.96 | 0.96 | 2xt6A | - | 0.51 |
| 1prrA | 0.60 | 0.72 | 3v7oB | 0.63 | 0.68 | 2zpaB | 0.90 | 0.90 |
| 1q19A | 0.99 | 0.99 | 3vr8B | 1.00 | 1.00 | 3apoA | 0.27 | 0.39 |
| 1qwrA | 1.00 | 0.99 | 3wkuA | - | 0.75 | 3b43A | 0.23 | 0.21 |
| 1r71B | 0.88 | 0.92 | 3zvmA | 0.53 | 0.57 | 3gf5B | 0.23 | 0.30 |
| 1rh1A | 0.55 | 0.59 | 4acoA | 0.92 | 0.93 | 3hjlA | 0.39 | 0.42 |
| 1rktA | 0.81 | 0.89 | 4ap5A | 0.98 | 0.98 | 3kq4B | 0.37 | 0.45 |
| 1s6lA | - | 0.73 | 4axdA | 0.68 | 0.66 | 3kwlA | 0.52 | 0.35 |
| 1sp3A | - | 0.93 | 4bfiB | 0.61 | 0.84 | 3ob8A | 0.38 | 0.37 |
| 1vz6A | 0.72 | 0.71 | 4bt9B | 0.59 | 0.61 | 3opfB | 0.49 | 0.40 |
| 1w3aA | 0.64 | 0.58 | 4cczA | 0.87 | 0.89 | 3p53A | - | 0.45 |
| 1wv3A | - | 0.92 | 4d0nB | 0.88 | 0.91 | 3pvlA | 0.47 | 0.38 |
| 1x7pA | 0.98 | 0.80 | 4d1iG | 0.81 | 0.80 | 3r05A | 0.26 | 0.28 |
| 1x9yA | - | 0.58 | 4dj3A | 0.98 | 0.98 | 3ubhA | 0.33 | 0.36 |
| 1y11A | - | 0.56 | 4dqaA | - | 0.62 | 3w2wA | 0.52 | 0.54 |
| 1yiqA | 0.94 | 0.88 | 4eo3A | 0.56 | 0.58 | 3zniA | - | 0.35 |
| 1zbuB | - | 0.77 | 4eogA | 0.99 | 0.99 | 4aimA | 0.60 | 0.52 |
| 1ze1A | 0.97 | 0.99 | 4etxA | - | 0.55 | 4ak1A | - | 0.29 |
| 2ablA | 0.67 | 0.68 | 4fguA | - | 0.70 | 4aq1A | - | 0.31 |
| 2ahvA | 0.79 | 0.80 | 4fkcA | 0.99 | 0.94 | 4fe9A | 0.35 | 0.48 |
| 2bkpA | 0.57 | 0.59 | 4fxkC | 0.65 | 0.97 | 4h2aA | 0.68 | 0.45 |
| 2c1yA | - | 0.58 | 4gbyA | 0.93 | 0.99 | 4i5sB | 0.42 | 0.44 |
| 2cxcA | 0.86 | 0.85 | 4ggmX | - | 0.60 | 4iggB | 0.29 | 0.31 |
| 2d1cA | 0.78 | 0.83 | 4gslA | 0.55 | 0.54 | 4j9vA | 0.47 | 0.58 |
| 2d7iA | 0.80 | 0.74 | 4gyjA | 0.99 | 0.99 | 4k3bA | 0.54 | 0.51 |
| 2e9hA | 0.97 | 0.73 | 4h3tA | 0.91 | 0.90 | 4kwuA | 0.48 | 0.51 |
| 2e9xB | - | 0.92 | 4hmoA | 0.91 | 0.93 | 4m00A | - | 0.52 |

| | | | | | | | | |
|---|---|---|---|---|---|---|---|---|
| 2evrA | 0.99 | 0.99 | 4ie6A | - | 0.69 | 1bp1A | 0.97 | 0.97 |
| 2ew9A | 0.58 | 0.55 | 4l5gA | 0.73 | 0.68 | 1cjsA | 0.79 | 0.86 |
| 2fd5A | 0.92 | 0.99 | 4lpqA | 0.70 | 0.64 | 1ck1A | 0.98 | 0.99 |
| 2gh8A | 0.72 | 0.71 | 4m8rA | - | 0.77 | 1ecrA | - | 0.86 |
| 2gt1A | 0.97 | 0.97 | 4n06B | 0.95 | 0.99 | 1f5qD | 0.97 | 0.99 |
| 2gzaC | 0.95 | 0.85 | 4nj5A | 0.62 | 0.61 | 1fa9A | - | 0.95 |
| 2hjqA | - | 0.53 | 4opaB | - | 0.82 | 1gu7A | 0.88 | 0.96 |
| 2hwjA | - | 0.69 | 4qkuB | - | 0.97 | 1itwA | - | 0.96 |
| 2ijd1 | 0.76 | 0.77 | 4up9A | 0.98 | 0.95 | 1jkiA | 1.00 | 0.99 |
| 2iu7A | - | 0.59 | 4w7sA | 0.64 | 0.64 | 1n80A | - | 0.84 |
| 2iw2A | 0.98 | 0.98 | 1bf2A | 0.95 | 0.99 | 1nzjA | 0.99 | 0.99 |
| 2jz4A | - | 0.57 | 1bhgA | 1.00 | 1.00 | 1qhdA | 0.63 | 0.93 |
| 2kdyA | - | 0.59 | 1f7uA | 0.94 | 0.94 | 1qz9A | 0.98 | 0.99 |
| 2kn4A | 0.62 | 0.61 | 1fx7A | 0.86 | 0.92 | 1sb7B | 0.87 | 0.89 |
| 2mbgA | 0.78 | 0.79 | 1griA | 0.52 | 0.61 | 1vk1A | 0.93 | 0.85 |
| 2nsfA | - | 0.67 | 1h88C | 0.41 | 0.45 | 1vrmA | 0.97 | 0.96 |
| 2nykA | - | 0.92 | 1m8pB | 0.45 | 0.41 | 1xvuA | 0.97 | 0.97 |
| 2o6yA | 0.97 | 0.96 | 1ni5A | 0.66 | 0.66 | 1yy3A | - | 0.70 |
| 2owbA | 0.99 | 0.97 | 1q25A | 0.45 | 0.47 | 1z87A | 0.56 | 0.55 |
| 2qfiA | - | 0.72 | 1uzjA | 0.53 | 0.52 | 2a1sC | - | 0.85 |
| 2qp2A | 0.87 | 0.74 | 1zpuA | 1.00 | 1.00 | 2a3lA | - | 0.99 |
| 2qygA | 0.99 | 0.98 | 1zy9A | 0.89 | 0.88 | 2bt1A | - | 0.90 |
| 2r5wB | 0.59 | 0.59 | 2b5uA | 0.49 | 0.47 | 2bydA | 0.67 | 0.66 |
| 2uu7A | 0.95 | 0.96 | 2ewfA | - | 0.57 | 2c43A | 0.66 | 0.68 |
| 2w4bA | - | 0.81 | 2piaA | 0.71 | 0.74 | 2dfyC | 0.52 | 0.54 |
| 2x7iA | 0.94 | 1.00 | 2r7dA | 0.95 | 0.93 | 2dlaA | - | 0.97 |
| 2x8kC | 0.96 | 0.96 | 2uwnA | 0.43 | 0.42 | 2g3pA | - | 0.60 |
| 2yilA | - | 0.64 | 2v0nA | 0.60 | 0.53 | 2gg6A | 0.93 | 1.00 |
| 2yrqA | 0.53 | 0.52 | 2vgmA | 0.77 | 0.60 | 2gsyE | - | 0.70 |
| 2zxcA | - | 0.82 | 2wqrB | 0.43 | 0.62 | 2gzoA | 0.85 | 0.86 |
| 3a1iA | 0.91 | 0.93 | 2y25B | 0.38 | 0.45 | 2j2cA | - | 0.77 |
| 3a45A | 0.97 | 0.98 | 2yk0A | 0.47 | 0.74 | 2kfwA | 0.77 | 0.77 |
| 3a56A | - | 0.66 | 2zzqA | - | 0.44 | 2l9yA | 0.69 | 0.69 |
| 3ajvA | - | 0.85 | 3bt1U | - | 0.49 | 2ntyB | - | 0.76 |
| 3aqkA | 0.98 | 0.98 | 3c1yA | 0.48 | 0.61 | 2r3vA | 0.95 | 0.96 |
| 3arbA | 0.84 | 0.94 | 3cw2C | 0.62 | 0.70 | 2r58A | - | 0.55 |
| 3aujG | - | 0.70 | 3f83A | - | 0.39 | 2w4mA | 0.91 | 0.85 |
| 3b2zF | 0.76 | 0.76 | 3fc3A | - | 0.49 | 2x0cA | 0.66 | 0.66 |
| 3b7wA | 0.82 | 0.83 | 3gbgA | 0.64 | 0.60 | 2y51A | 1.00 | 0.99 |
| 3bt3A | - | 0.96 | 3h5cB | 0.56 | 0.98 | 2yb0E | - | 0.79 |
| 3c4tA | 0.75 | 0.74 | 3ibjA | 0.57 | 0.50 | 2z86C | 0.67 | 0.68 |
| 3craA | 0.65 | 0.57 | 3ippB | 0.45 | 0.44 | 3afoA | 0.67 | 0.68 |
| 3d30A | - | 0.94 | 3jymB | 0.47 | 0.43 | 3bu2A | 0.74 | 0.91 |
| 3eo5A | - | 0.59 | 3kbgA | - | 0.49 | 3cvzA | - | 0.76 |

| | | | | | | | | |
|---|---|---|---|---|---|---|---|---|
| 3errA | 0.88 | 0.80 | 3mc8A | 0.41 | 0.43 | 3dupA | - | 0.87 |
| 3g79A | 0.81 | 0.90 | 3npfA | 0.92 | 0.92 | 3eswA | 0.88 | 0.88 |
| 3h2tA | - | 0.81 | 3orjA | 0.73 | 0.74 | 3eukH | - | 0.90 |
| 3hcsA | 0.70 | 0.67 | 3plaA | 0.49 | 0.53 | 3fi7A | 0.92 | 0.93 |
| 3hyiA | - | 0.69 | 3qe9Y | 0.71 | 0.72 | 3fvvA | 0.91 | 0.96 |
| 3i2dA | 0.79 | 0.84 | 3qjoA | 0.68 | 0.67 | 3gmsA | 0.94 | 0.99 |
| 3iam2 | 0.93 | 0.93 | 3qphA | 0.54 | 0.47 | 3hzzB | 1.00 | 0.99 |
| 3ifrA | 0.91 | 0.98 | 3qyeA | 0.96 | 0.95 | 3m1uA | 0.99 | 0.99 |
| 3isqA | 0.99 | 0.99 | 3rimA | 0.99 | 0.98 | 3mw8A | 0.98 | 0.64 |
| 3j7aK | 0.98 | 0.98 | 3rrpA | 0.95 | 0.95 | 3mwcA | 0.99 | 0.99 |
| 3k1rA | 0.59 | 0.60 | 3soaA | 0.59 | 0.56 | 3nsjA | 0.80 | 0.82 |
| 3k2iA | - | 0.75 | 3tixD | - | 0.60 | 3ntkA | 0.92 | 0.92 |
| 3kh5A | 0.96 | 0.96 | 3tp9A | 0.60 | 0.57 | 3oaaG | 0.98 | 0.98 |
| 3kjpA | 0.94 | 0.58 | 3ua3A | 0.51 | 0.50 | 3ptyA | - | 0.71 |
| 3kt1A | 0.89 | 0.90 | 3uj0A | 0.91 | 0.91 | 3rfyA | - | 0.90 |
| 3ktmE | - | 0.64 | 3vn4A | 0.43 | 0.43 | 3seoB | - | 0.81 |
| 3kzwA | 0.98 | 0.97 | 3vsmA | - | 0.87 | 3spgA | 0.95 | 0.98 |
| 3l76A | 0.88 | 0.89 | 3w1bA | 0.72 | 0.73 | 3u0kA | 0.62 | 0.62 |
| 3ld1A | - | 0.69 | 3zh9B | - | 0.59 | 3vlaA | 0.99 | 0.98 |
| 3lsgA | 0.99 | 0.85 | 4alzA | - | 0.69 | 3vstA | - | 0.98 |
| 3me4A | 0.77 | 0.77 | 4ax8A | 0.51 | 0.52 | 4aqfB | - | 0.87 |
| 3ml4C | - | 0.63 | 4b3iA | 0.95 | 0.95 | 4b21A | 0.98 | 0.90 |
| 3mx2B | - | 0.70 | 4bd9B | 0.39 | 0.40 | 4dt4A | 0.71 | 0.93 |
| 3mzfA | 0.86 | 0.86 | 4c0aB | 0.46 | 0.48 | 4dtfA | - | 0.85 |
| 3njaB | 0.61 | 0.61 | 4c0sA | 0.90 | 0.61 | 4ewtA | 0.92 | 0.87 |
| 3nqiA | - | 0.95 | 4dimA | 0.87 | 0.86 | 4f23A | 0.97 | 0.97 |
| 3nt8A | 0.78 | 0.57 | 4indA | - | 0.45 | 4fzbC | 0.95 | 0.99 |
| 3og5A | 0.61 | 0.89 | 4jdzB | 0.69 | 0.67 | 4g1pA | 0.80 | 0.93 |
| 3oh0A | 0.79 | 0.79 | 4kc3B | 0.43 | 0.75 | 4gfqA | 0.70 | 0.70 |
| 3pcsB | - | 0.80 | 4kikB | 0.54 | 0.72 | 4hvzA | - | 0.57 |
| 3po3S | 0.56 | 0.56 | 4lmfA | 0.45 | 0.45 | 4il6B | 0.83 | 0.72 |
| 3pxpA | - | 0.68 | 4lziA | 0.41 | 0.43 | 4jxkA | 0.93 | 0.96 |
| 3qavA | 0.93 | 0.93 | 4m9pA | 0.42 | 0.49 | 4m8mB | 0.82 | 0.85 |
| 3qf4B | 0.65 | 0.93 | 4pt5A | 0.93 | 0.88 | 4mzyA | 0.73 | 0.99 |
| 3qjjA | 0.97 | 0.98 | 4uwhA | 0.98 | 1.00 | 4onyA | 0.81 | 0.80 |
| 3qtdA | 1.00 | 1.00 | 1c1zA | 0.29 | 0.28 | 4pyhA | 0.95 | 0.75 |
| 3r6bA | 0.52 | 0.52 | 1d2pA | - | 0.47 | 4rg1A | 0.92 | 0.66 |
| 3rh7A | - | 0.58 | 1k7tA | 0.42 | 0.32 | | | |

**Table S20.** On 20 human proteins, the TM-score of full-chain models assembled by SADA and predicted by AlphaFold2. #Domain represents the number of domains. SADA1 represents that the domain models for SADA assembly were decomposed from the full-chain structures of AlphaFold2. SADA2 represents that the domain models for SADA assembly were predicted by AlphaFold2. AF2 represents the full-chain model predicted by AlphaFold2.

| PDB ID | Length | #Domains | TM-score | | |
|---|---|---|---|---|---|
| | | | AF2 | SADA1 | SADA2 |
| 1b3uA | 589 | 3 | 0.72 | 0.75 | 0.81 |
| 1ggzA | 149 | 2 | 0.59 | 0.49 | 0.71 |
| 1lfsA | 340 | 3 | 0.79 | 0.79 | 0.73 |
| 1qbkB | 898 | 3 | 0.79 | 0.84 | 0.84 |
| 1s8oA | 555 | 2 | 0.72 | 0.97 | 0.92 |
| 1st0A | 337 | 2 | 0.78 | 0.77 | 0.84 |
| 1xa6A | 468 | 4 | 0.54 | 0.70 | 0.81 |
| 1ytqA | 205 | 2 | 0.49 | 0.50 | 0.49 |
| 2ar7A | 223 | 2 | 0.75 | 0.82 | 0.81 |
| 2hxyA | 411 | 2 | 0.53 | 0.53 | 0.54 |
| 2kdoA | 250 | 3 | 0.43 | 0.54 | 0.59 |
| 2l6lA | 149 | 2 | 0.38 | 0.39 | 0.46 |
| 2nn6I | 195 | 2 | 0.80 | 0.74 | 0.63 |
| 2wzbA | 417 | 2 | 0.77 | 0.72 | 0.72 |
| 3ajmA | 212 | 2 | 0.69 | 0.68 | 0.67 |
| 3t5oA | 934 | 3 | 0.61 | 0.51 | 0.51 |
| 5ivw2 | 462 | 2 | 0.40 | 0.39 | 0.37 |
| 5t7cA | 193 | 2 | 0.43 | 0.43 | 0.35 |
| 5ztfA | 1042 | 2 | 0.65 | 0.65 | 0.61 |
| 6nr8A | 556 | 2 | 0.70 | 0.69 | 0.69 |

### Supplementary Texts

**Text S1. The used parameters in CD-HIT.**

The sequence identity threshold (c) was set to 1.0, the alignment coverage for the longer sequence (aL) was set to 1.0, and other parameters were set to default.

**Text S2. The details of culling module.**

Users need to input the criteria, including protein chain length, resolution, number of domains, R-factor and sequence identity of multi-domain protein, and then the module retrieves whole MPDB according to the criteria. After the retrieval was finished, the module sends the user an email that includes a link to the resulting files. From these files, the user can download the multi-domain proteins that satisfy the criteria and the corresponding information file recording the protein name, chain length, experimental measurement methods, resolution, R-factor, number of domains, and methods of defining domain boundary and domain boundary. In the culling module, these criteria were applied first the list of MPDB entries. The result is a list of MPDB chains that fit the criteria that can then be culled according to mutual sequence identity.

**Text S3. The details of structural analogue detection module.**

In the structural analogue detection module, the mandatory inputs for the web server are individual domain structure files in PDB format and target sequence, where the sequence is used to calculate sequence identity.

Upon job completion, the user will be notified via email that contains two links to the results display page and results download page on the structural analogues, respectively. The results display page shows an ordered list of the top 10 structural analogues with the highest $LS_{\text{score}}$ from the MPDB. The structural analogues are displayed in three dimensions, allowing users to rotate the pictures, and a link is provided to the URL addresses for downloading the analogue structure from PDB. The structural analogue is shown together with the $LS_{\text{score}}$, TM-score of each input domain aligned on the analogue, aligned length between input sequence and structural analogue sequence, sequence identical length and sequence identity, where the aligned length, identical length and sequence identity are calculated using the NW-align program (http://zhanglab.dcmb.med.umich.edu/NW-align). In the results download page, a compressed file containing structures of top 200 multi-domain protein structural analogues with the highest $LS_{\text{score}}$ and an analogues information file including the information mentioned above can be downloaded.

**Text S4. Description of the two-stage differential evolution algorithm and parameters setting.**

The proposed two-stage differential evolution algorithm is described as follows:

**Exploration stage:**

1. ***Initialization***. The population size *NP* and maximum generation $G_{\text{explore}}$ were set. The initial population should maximise the coverage of the entire search space by uniformly randomising individuals within the search space constrained by the prescribed minimum and maximum parameter bounds $BD_{\text{explore}}^{\min} = (x_1^{\min}, y_1^{\min}, z_1^{\min}, \phi_1^{\min}, \psi_1^{\min}, \omega_1^{\min}, ..., x_N^{\min}, y_N^{\min}, z_N^{\min}, \phi_N^{\min}, \psi_N^{\min}, \omega_N^{\min})$ and $BD_{\text{explore}}^{\max} = (x_1^{\max}, y_1^{\max}, z_1^{\max}, \phi_1^{\max}, \psi_1^{\max}, \omega_1^{\max}, ..., x_N^{\max}, y_N^{\max}, z_N^{\max}, \phi_N^{\max}, \psi_N^{\max}, \omega_N^{\max})$, where *N* is the number of domains. Thus, each solution $S_{\text{explore},i}$ in initial solution population $P1 = \{S_{\text{explore},1}, S_{\text{explore},2}, ..., S_{\text{explore},NP}\}$ are generated as follows:

$$S_{\text{explore},i} = BD_{\text{explore}}^{\min} + \text{rand}(0,1) \times (BD_{\text{explore}}^{\max} - BD_{\text{explore}}^{\min})$$

where $i = \{1, 2, ..., NP\}$ and rand(0,1) represents a uniformly distributed random variable within the range (0, 1). The generation counter of exploration stage ($g_{\text{explore}}$) was set to 1.

2. ***Mutation operation***. The *i*-th solution $S_{\text{explore},i}$ in the population of the $g_{\text{explore}}$-th generation was set to the target solution $S_{\text{explore,target}}$, and performed the following operations:

   **2.1.** Three mutually different solutions were randomly selected in current solution population, namely, $S_{\text{explore},rand1}$, $S_{\text{explore},rand2}$ and $S_{\text{explore},rand3}$. The mutant vector $V_{\text{explore},i}$ was generated as follows:

$$V_{\text{explore},i} = S_{\text{explore},rand1} + F_{\text{explore},i} \times (S_{\text{explore},rand2} - S_{\text{explore},rand3})$$

   where $F_{\text{explore},i}$ is scale factor of exploration stage in the *i*-th solution. If the values of $F_{\text{explore},i} \times (S_{\text{explore},rand2} - S_{\text{explore},rand3})$ exceed the corresponding $BD_{\text{explore}}^{\max}$ and $BD_{\text{explore}}^{\min}$, we randomly and uniformly reinitialise them within the prespecified range.

3. ***Crossover operation***. After the mutation phase, crossover operation was applied to each pair of the target vector $S_{\text{explore,target}}$ and its corresponding mutant vector $V_{\text{explore},i}$ to generate a trial vector $U_{\text{explore},i}$ as follows:

$$U_{\text{explore},i}^{j} = \begin{cases} V_{\text{explore},i}^{j}, & \text{if } (\text{rand}[0,1) < CR_{\text{explore},i}) \text{ or } (j = j_{\text{rand}}) \\ S_{\text{explore,target}}^{j}, & \text{otherwise} \end{cases}, \quad j = 1, 2, 3, ..., 6*N.$$

   where $CR_{\text{explore},i}$ is the crossover rate of exploration stage in the *i*-th solution, which controls the fraction of parameter values copied from the mutant vector. *j* is the dimension of the corresponding vector. $j_{\text{rand}}$ is a randomly chosen integer in the range [1, 6**N*].

4. ***Selection operation***. The energy function value of trial vector was evaluated. The selection operation can be expressed as follows:

$$S_{\text{explore},i} = \begin{cases} U_{\text{explore},i}, & \text{if } E_{\text{total}}(U_{\text{explore},i}) \le E_{\text{total}}(S_{\text{explore,target}}) \\ S_{\text{explore,target}}, & \text{otherwise} \end{cases}$$

5. ***Iteration***. The following steps were performed to check iteration:
   **5.1.** If $i$ is equal to the population size $NP$ then execute step 5.2; Otherwise $i = i + 1$ and go to step 2~5.
   **5.2.** Every $LP$ generation, the solution with the lowest energy is selected from $P1$ into a new population $P2$, and go to step 5.3.
   **5.3.** If the solution with the lowest energy was not replaced in consecutive 1000 generations, then go to exploitation stage; Otherwise $g_{\text{explore}} = g_{\text{explore}} + 1$, $i = 1$, and execute steps 2~5.

**Exploitation stage:**

1. ***Initialization***. The new population $P2$ generated in exploration stage is regarded as the initial population of exploitation stage. The initial generation counter of exploration stage ($g_{\text{exploit}}$) was set to 1.

2. ***Mutation operation.*** The $i$-th solution $S_{\text{exploit},i}$ in the population $P2$ of the $g_{\text{exploit}}$-th generation was set to the target conformation. The following operations were performed:
   **2.1.** Two mutually different solutions were randomly selected in $P2$, namely, $S_{\text{exploit},rand1}$ and $S_{\text{exploit},rand2}$. The mutant vector $V_{\text{exploit},i}$ was generated as follows:

$$V_{\text{exploit},i} = S_{\text{exploit,target}} + F_{\text{exploit}} \times (S_{\text{exploit},rand1} - S_{\text{exploit},rand2})$$

   where $F_{\text{exploit}}$ is scale factor of exploitation stage. In this stage, if the values of $F_{\text{exploit}} \times (S_{\text{exploit},rand1} - S_{\text{exploit},rand2})$ exceed the corresponding $BD^{\max}_{\text{exploit}}$ and $BD^{\min}_{\text{exploit}}$, we randomly and uniformly reinitialise them within the prespecified range.

3. ***Crossover operation***. After the mutation phase, crossover operation was applied to each pair of the target vector $S_{\text{exploit,target}}$ and its corresponding mutant vector $V_{\text{exploit},i}$ to generate a trial vector $U_{\text{exploit},i}$ as follows:

$$U^{j}_{\text{exploit},i} = \begin{cases} V^{j}_{\text{exploit},i}, & \text{if } (\text{rand}[0,1) < CR_{\text{exploit}}) \text{ or } (j = j_{\text{rand}}) \\ S^{j}_{\text{exploit,target}}, & \text{otherwise} \end{cases}, \quad j = 1, 2, 3, \ldots, 6*N.$$

   where $CR_{\text{exploit}}$ represents the crossover rate of exploitation stage.

4. ***Selection operation***. If the trial vector has less or equal objective function value than the corresponding target vector, the trial vector will replace the target vector and enter the population of the next generation. Otherwise, the target vector will remain in the population for the next generation.

5. ***Iteration***. The following steps were performed to check the iteration:
   **5.1.** If $i$ is equal to the size of $P2$ then execute step 5.2; Otherwise $i = i + 1$ and go to step 2~5.
   **5.2.** If the solution with the lowest energy was not replaced in consecutive 500 generations, the model with the lowest $E_{\text{total}}$ in $P2$ was output; Otherwise $g_{\text{exploit}} = g_{\text{exploit}} + 1$, $i = 1$, and go to step 2~5.

**Parameters setting:**

Population size:

$NP = 50$;

Scale factor of exploration stage in the *i*-th solution:

$F_{\text{explore},i} = 0.1 + 0.9 \times rand(0,1),\ i = 1, 2, 3, ..., NP$;

Crossover rate of exploration stage in the *i*-th solution:

$CR_{\text{explore},i} = rand(0,1),\ i = 1, 2, 3, ..., NP$;

Learning period:

$LP = 100$;

The prescribed minimum parameter bounds of exploration stage:

$BD^{\min}_{\text{explore}} = (-5, -5, -5, 0, 0, 0, ..., -5, -5, -5, 0, 0, 0)$;

The prescribed maximum parameter bounds of exploration stage:

$BD^{\max}_{\text{explore}} = (5, 5, 5, 2\pi, \pi, 2\pi, ..., 5, 5, 5, 2\pi, \pi, 2\pi)$;

Scale factor of exploitation stage:

$F_{\text{exploit}} = 0.5$;

Crossover rate of exploitation stage:

$CR_{\text{exploit}} = 0.5$;

The prescribed minimum parameter bounds of exploitation stage:

$BD^{\min}_{\text{exploit}} = (-0.2, -0.2, -0.2, 0, 0, 0, ..., -0.2, -0.2, -0.2, 0, 0, 0)$;

The prescribed maximum parameter bounds of exploitation stage:

$BD^{\max}_{\text{exploit}} = (0.2, 0.2, 0.2, 0.5, 0.25, 0.5, ..., 0.2, 0.2, 0.2, 0.5, 0.25, 0.5)$
